# TGS1 mediates mRNA 5’-cap trimethylation to promote oxidative phosphorylation in acute myeloid leukaemia

**DOI:** 10.64898/2026.03.18.712389

**Authors:** Valentina Miano, Giovanna Carrà, Maxim Bouvet, Jan Lj Miljkovic, Vincent Paupe, Chak Shun Yu, Domenico Ignoti, Sara Milioni, Lorenzo Manfreda, Daniele Viavattene, Anna M. Benedetti, Angelo D’angelo, Shi-Lu Luan, Davide Acquarone, Valeria Poli, Paolo E. Porporato, Stefano Gustincich, Suzanne D. Turner, Maria Paola Martelli, Adriano Setti, Alessio Menga, Lorenzo Brunetti, Angela Maria Savino, Michael P. Murphy, Luca Pandolfini, Isaia Barbieri

## Abstract

The Trimethylguanosine Synthase (TGS1) is a highly conserved enzyme mediating di-methylation of the 5’-cap 7-methylguanosine of RNA to generate 2,2,7-trimethylguanosine (m^2,2,7^G). Known TGS1 targets include snRNAs, snoRNAs, the telomeric RNA component and a limited number of mRNAs encoding selenoproteins.

TGS1 is highly expressed in acute myeloid leukaemia (AML) and correlates with poorly differentiated leukaemic subtypes, including P53-mutant leukaemias. Here, we report that TGS1 directly methylates the cap of more than 500 mRNAs in AML cells. Specifically, we demonstrate that TGS1 modifies nuclear mRNAs encoding mitochondrial proteins, including critical components of complexes involved in both the TCA cycle and oxidative phosphorylation, promoting their translation mediated by mitochondria associated cytosolic ribosomes. Functionally, we report that TGS1 depletion impairs mitochondrial respiration and increases oxidative stress. This, in turn, impairs the growth of AML cells causing differentiation and cell cycle arrest *in vitro* and *in vivo*. Finally, we demonstrate that TGS1 depletion sensitizes AML cells to RSL3, a small molecule promoting ferroptosis. Taken together, our findings establish TGS1 as a key regulator of oxidative phosphorylation and mitochondrial redox status of AML cells and highlight its potential as a therapeutic target in leukaemia.

## Main

The vast majority of primary RNA transcripts undergo post transcriptional maturation processes. These include splicing, 5’-capping, 3’ polyadenylation and the introduction of a large variety of heterogeneous chemical moieties within nucleotides^1^. Overall, the mechanisms of RNA post-transcriptional modification represent an additional layer to gene expression control that can impact all aspects of RNA biology such as nuclear export, stability, activity and translation^1^. Eukaryotic mRNAs and nuclear non-coding RNAs such as snRNAs and snoRNAs are co-transcriptionally capped by addition of a guanosine nucleotide at the 5’-end of the RNA through a non-canonical 5’-triphosphate-5’ bond^2^. Next, the guanosine methyltransferase RNMT introduces a methyl group on the guanosine cap to generate 7-methylguanosine^3^.

In the nucleus the main role of the 5’ RNA-cap is to protect newly synthesized RNA from degradation by 5’-3’ exonucleases. The m^7^G cap on Pol II generated transcripts is also necessary for recruitment of the cap-binding complex, which, in turn, mediates mRNA processing and export. Finally, in the cytoplasm, the presence of m^7^G is required for efficient initiation of cap-dependent translation^2^.

In vertebrates, mRNA 5’ ends can be further modified through ribose 2′-O-methylation on the first (cap1) and second (cap2) transcribed nucleotides. These modifications are mediated by cap methyltransferase 1 and 2 (CMTR1/2), respectively^2^. Little is known about the specific functions of these cap variants, but they appear to be positively correlated with mRNA half-life^4^.

Several less abundant types of alternative 5’ RNA caps have also been identified in both prokaryotic and eukaryotic cells including NAD, FAD and acetyl-CoA. In bacteria NAD-cap protects RNA from degradation^5^. Similarly, mitochondrial mRNAs are NAD-capped in response to specific metabolic conditions, to regulate gene expression. In contrast to prokaryotes, NAD-capped cytoplasmic mRNAs are rapidly degraded, in both yeast and human cells^6^.

5’-cap hypermethylation was originally identified in yeast snoRNAs and snRNAs, where a specific guanosine N^2^-methyltransferase, Trimethyl guanosine Synthase 1 (TGS1), introduces two methyl groups to the m^7^G-cap, forming 2,2,7-trimethylguanosine (m^2,2,7^G)^7^. It was reported that m^2,2,7^G is required for nucleus-cytoplasmic shuffling of snRNAs, an essential step for their maturation^8^. More recently, this specific cap hypermethylation was reported on the telomeric RNA component (*TERC*) in both yeast and mammalian cells, where it protects *TERC* stability^9^, and on a few mRNAs encoding selenoproteins, where it promotes the specific translation mechanisms required for selenocysteine incorporation^10^. Despite recent advances in RNA biology and epitranscriptomic mechanisms, the roles of *TGS1* and m^2,2,7^G under physiological conditions remain poorly understood, and its involvement in disease, including cancer, is largely unknown.

Acute myeloid leukaemia (AML) is one of the most aggressive types of blood cancers, with a 5-year survival rate of just 20%^11^. The disease is characterized by an overwhelming proliferation of myeloid blasts that rapidly become the predominant population in both the bone marrow and peripheral blood. If not immediately treated, AML leads to severe anaemia and death. Several therapeutic strategies are used to treat AML, including chemotherapy and bone marrow transplantation^11^. Unfortunately, most patients, after initial remission, undergo relapse, and once all therapeutic options have been used, the prognosis is extremely bleak. It is therefore of utmost urgency to develop new therapeutic strategies for AML, to extend the range of available treatments and increase survival.

AML can be driven by heterogeneous oncogenic mutations, including MLL-fusions, driving the expression of genes promoting proliferation and stemness of myeloid precursors, such as the *HOX* genes^12^. The majority of AML patients carry mutations of the Nucleophosmin gene (*NPM*) which inactivates the nuclear localization signal of the encoded protein, causing its translocation to the cytoplasm^13^.

Recently, epitranscriptomic factors involved in m^6^A RNA modification pathways were shown to play a crucial role in several cancer types^14^. In particular, both m^6^A writers, METTL3^15^, METTL16^16^ and m^6^A erasers, such as FTO^17^, were identified as potential therapeutic targets in AML together with the m^6^A reader proteins, such as YTHDF2^18^.

Leukaemia stem cells, differently from many types of solid tumours, shift their metabolic machinery towards oxidative phosphorylation (OXPHOS) compared to normal haematopoietic stem cells. Importantly, this metabolic shift appears to be one of the main drivers of transformation in AML, rather than being an adaptive response^19^. Conversely, reducing OXPHOS in AML cells induces cell cycle arrest and terminal differentiation^20^. These findings underpin the rationale for using OXPHOS inhibitors to specifically target leukaemic cells and treat leukaemia patients.

The mitochondrial inner membrane harbours 5 large metabolic complexes driving OXPHOS. Complexes I, II, III, and IV generate a proton electrochemical potential gradient across the mitochondrial inner membrane, using oxygen as the terminal electron acceptor. In turn, complex V (ATP synthase) utilizes this electrochemical gradient to drive ATP production^21^. These complexes comprise many subunits encoded by both mitochondrial and nuclear transcripts. Their assembly is a highly coordinated process involving the translocation of nuclear encoded subunits through the outer mitochondrial membrane and the insertion of mitochondrial encoded subunits within the inner mitochondrial membrane^22^. It is well known that translation of nuclear encoded mitochondrial complexes’ subunits occurs in close proximity to the mitochondrial outer membrane and the newly synthesized peptides are rapidly translocated across the membrane^23^. The molecular mechanisms involved in both mRNA sub-localization and the regulation of mitochondrial protein translation are largely unknown. The efficiency of mitochondrial protein translation and import may have relevant effects on cellular metabolism and its dysregulation could potentially be involved in disease.

Mitochondria are also involved in triggering several different types of controlled cell death. In particular, release of cytochrome *c* from mitochondrial membranes is one of the main pathways leading to apoptosis^24^. Moreover, it has been shown that aberrant mitochondrial metabolism is involved in ferroptosis^25^, a form of non-apoptotic cell death initiated through iron-dependent oxidation of lipids via Fenton chemistry^26^, causing accumulation of intracellular lipid peroxides. The propagation of lipid peroxidation ultimately compromises the integrity of cellular membranes, culminating in cell death. Dysfunction, mutation, or aberrant expression of *GPX4*, a selenoprotein belonging to the class of antioxidative defence enzymes, alongside dysregulation of CoQ_10_ redox status, iron homeostasis and glutathione biosynthesis, constitute key molecular drivers of ferroptotic cell death^27^. The development of different ferroptosis-inducers, such as RSL3 and Erastin, as well as inhibitors, such as Ferrostatin-1 and Liproxstatin-1, paved the way for regulation of ferroptotic pathways in many diseases, including their potential application as anticancer treatments^28^.

Here, we identify a new role for *TGS1* in AML cells. By directly hyper-methylating a subset of nuclear-transcribed mRNAs, including transcripts encoding for mitochondrial proteins, TGS1 mediates their translation. In turn, efficient translation of OXPHOS subunits promotes AML metabolism and maintains leukaemia cells’ growth. Conversely, we report that TGS1 depletion suppresses this elevated mitochondrial OXPHOS, induces oxidative stress affecting proliferation of AML cells *in vivo* and *in vitro*, and sensitizes AML cells to ferroptosis. Taken together, our data uncover *TGS1* as a key regulator of leukaemia metabolism and highlighting its potential relevance as a possible new therapeutic target in AML either alone or in combination with other metabolic inhibitors.

## Results

### TGS1 affects growth of AML cells

In our previous study, we performed both a genome wide and a targeted CRISPR-Cas9 dropout screen in RN2C mouse primary AML cells, identifying several RNA enzymes as potential therapeutic targets in AML^15^. *Tgs1* ranked amongst the top candidates in the targeted screen, specifically designed with a gRNA-library targeting the catalytic domain of RNA enzymes (Fig. 1a). To validate the findings of the original screens, we first performed a growth competition assay in RN2C cells, using two independent gRNAs targeting the catalytic domain of Tgs1 (Fig. 1b). After 8 days of culture *in vitro*, RN2C cells expressing *Tgs1*-targeting gRNAs, together with GFP, were significantly depleted compared to a negative control targeting the non-coding *Rosa26* locus. Remarkably, cellular depletion following CRISPR-mediated *Tgs1* knock-out was significantly higher than for cells lacking the positive control gene replication protein A (*Rpa3*).

**Figure 1.**
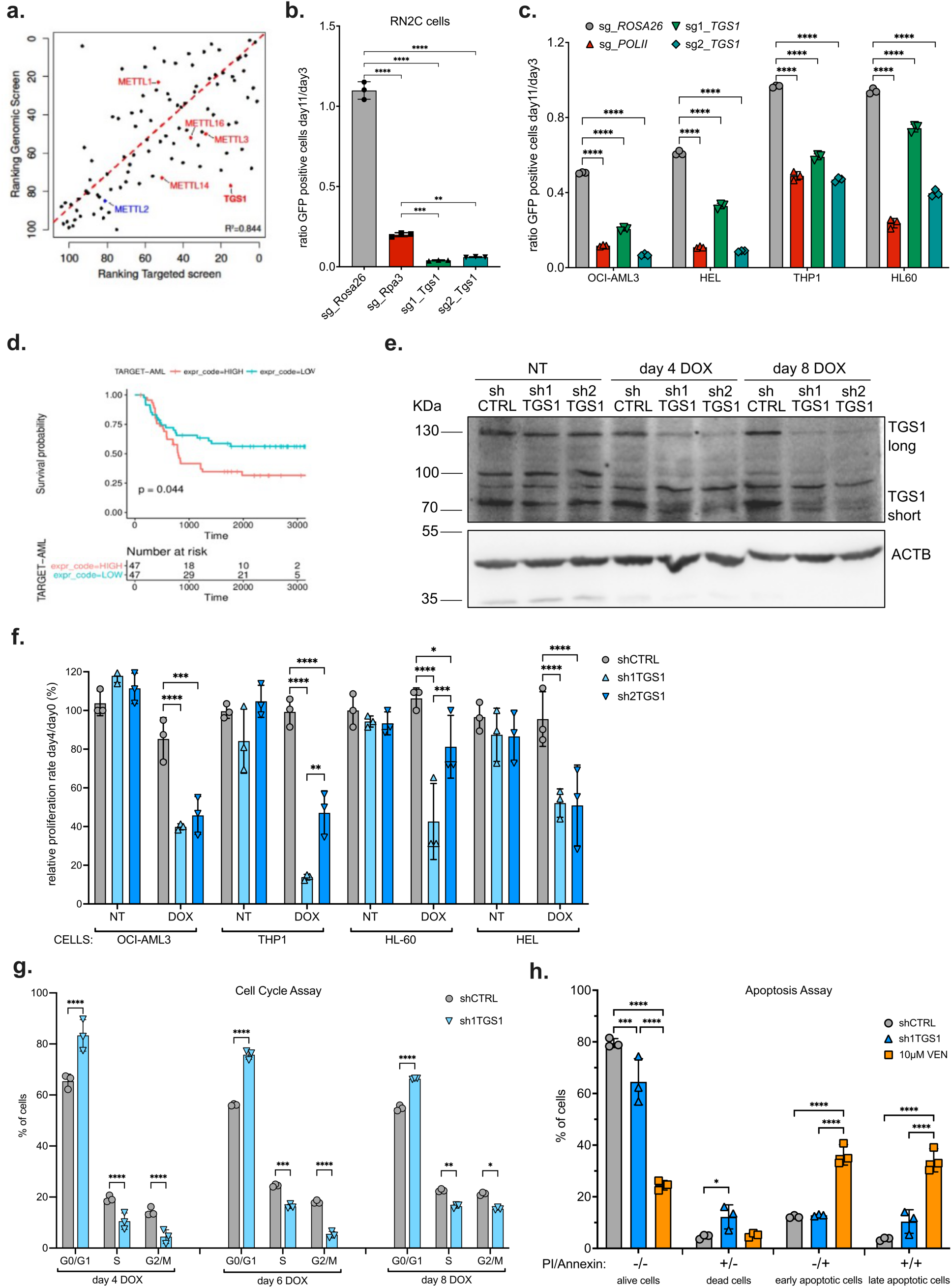
TGS1 is essential for the proliferation of AML cells. **a**. Correlation between genome-wide and RNA methyltransferase catalytic domain-targeted CRISPR dropout screens performed in RN2C cells, as previously reported^15^. The plot reports the ranking of the top 100 targets detected by each screen. **b.** Competitive co-culture assay showing negative selection of GFP-positive RN2C cells upon CRISPR-CAS9 targeting of *Tgs1*, *Rpa3* (positive control) or Rosa26 locus (negative control), between days 3 and 11 after infection. **c.** Competitive co-culture assay showing negative selection of GFP-positive cells in four different human leukaemia cell lines upon targeting *TGS1*, *POLII* (positive control) or Rosa26 locus (negative control), between days 3 and 11 after infection. **d.** Kaplan-Meier plot showing data from the TARGET-AML dataset comparing the top 75th percentile of patients (High) and 25th percentile (Low) for *TGS1* expression. **e.** Representative immunoblot of TGS1 protein levels in shCTRL, sh1TGS1 and sh2TGS1 OCI-AML3 cells untreated (NT) and following DOX treatment, at the indicated time points. ACTB was used as a loading control. **f.** Proliferation assay of the indicated human AML cell lines measured between day 0 and day 4; for DOX treated cells, these time points correspond to 4 days and 8 days of treatment, respectively. **g.** Flow cytometric assessment of cell cycle stage by Propidium Iodide (PI) staining of shCTRL and sh1TGS1 OCI-AML3 cells after indicated DOX treatment. **h.** Flow cytometric detection of PI/Annexin V stained shCTRL or sh1TGS1 OCI-AML3 cells following 8 days of DOX treatment. Values in **b**, **c**, **f**, **g** and **h** are mean ± SD of *n* = 3 independent experiments. **P<0.01 and ***P<0.001, one-way ANOVA test with Tukey’s multiple comparisons was used in **b**. *P<0.05, **P<0.01, ***P<0.001 and ****P<0.0001 two-way ANOVA test with Tukey’s (**c**, **f** and **g**) or Šidák’s (**h**) multiple comparisons.

Next, we confirmed the lethal effect of targeting *TGS1* in human AML cell lines, demonstrating that CRISPR-mediated *TGS1* depletion indeed affected the proliferation of leukaemia cell lines comprising different AML subtypes (Fig. 1c). Similar to mouse leukaemia cells, we observed a strong specific depletion of GFP-positive, *TGS1*-targeted AML cells in growth competition assays. Our data confirmed the high dependence of both mouse and human AML cells upon *TGS1*.

To evaluate TGS1 as a potential therapeutic target in leukaemia, we interrogated publicly available gene expression datasets in primary AML samples. *TGS1* is highly expressed in AML compared to other cancer types (Supplementary Fig. 1a) and its high expression levels significantly correlate with poor prognosis in the TARGET-AML dataset including pediatric refractory AML samples (Fig. 1d). Additionally, we report that *TGS1* expression shows a positive correlation with poor prognosis, albeit not statistically significant, in adult AML leukaemia datasets (Supplementary Fig. 1b). Importantly, in this dataset high expression of *TGS1* significantly associates with *P53* and *SRSF2* mutations, typically found in more aggressive AML subtypes^29,30^ and is associated with a less differentiated phenotype of leukaemic blasts (Supplementary Fig. 1c and 1d). Taken together, our data show TGS1 as a relevant regulator in AML, with potential implications for understanding disease biology, particularly in patients with poorer prognosis.

To generate inducible *TGS1* loss of function cellular models, we derived human AML cell lines expressing two different, doxycycline (DOX) inducible, shRNAs targeting *TGS1*, (sh1TGS1, sh2TGS1), and a non-targeting control shRNA (shCTRL). In all AML cell lines, we observed a strong decrease of both TGS1 isoforms upon doxycycline treatment (Fig. 1e and Supplementary Fig. 1e, f). The short TGS1 isoform does not originate from a specific mRNA but is a product of post-translational proteolysis of full length TGS1. Both isoforms are catalytically competent and only differ in their subcellular localization. While the short isoform is found in both the cytoplasm and the nucleus, the long isoform is exclusively cytoplasmic^31^. Cell fractionation experiments followed by immunoblotting confirmed subcellular localization of TGS1 isoforms and show efficient depletion of both nuclear and cytoplasmic TGS1 levels in DOX-treated sh1TGS1 OCI-AML3 cells (Supplementary Fig. 1g). Next, we measured the growth rates of TGS1-depleted AML cell lines and observed a significant decrease in cell proliferation (Fig. 1f). Finally, cell cycle analysis confirmed that TGS1 knock-down cells accumulate in G0/G1, together with decreased levels of cells in S and G2/M phases compared to shCTRL cells (Fig. 1g and Supplementary Fig. 1h). Despite this, sh1TGS1 and sh2TGS1 OCI-AML3 cells did not show signs of increased apoptosis in comparison to negative (shCTRL cells) or positive controls (cells treated with Venetoclax), as demonstrated by Annexin V staining (Fig. 1h and Supplementary Fig 1i).

To assess the global effects of TGS1 depletion on gene expression of leukaemia cells, we performed bulk RNA-seq on shCTRL and sh1TGS1 OCI-AML3 cells. Differential gene expression analysis identified 189 upregulated and 209 downregulated genes in sh1TGS1 cells (Supplementary Data 1a and Supplementary Fig. 1j). Gene ontology analysis of differentially expressed genes provided further insights into the effect of TGS1 depletion in leukaemia cells (Supplementary Data 1b, c). Specifically, transcripts overexpressed upon TGS1 knock-down were enriched in genes involved in leukocyte differentiation (GO:0002521), myeloid leukocyte differentiation (GO:0002573) and negative regulation of cell proliferation (GO:0008285) (Supplementary Fig. 1k). In contrast, downregulated transcripts were enriched for proinflammatory pathways such as neutrophil degranulation (GO:0043312) and myeloid leukocyte mediated immunity (GO:0002444). Importantly, it has been previously reported that high expression levels of genes involved in these proinflammatory pathways promote cell proliferation and survival of transformed leukaemic blasts^32^.

Taken together, our results indicate that TGS1 depletion specifically disrupts the proliferation of both mouse and human AML cell lines by inducing cell cycle arrest and differentiation.

### Loss of TGS1 impairs leukaemia growth *in vivo*

We next assessed the effect of inducible TGS1 depletion on the leukemogenic potential of OCI-AML3 cells *in vivo*. shCTRL and sh1TGS1 OCI-AML3 cells were injected subcutaneously into NSG mice, and tumour formation was monitored for 15 days (Fig. 2a). Following doxycycline administration in drinking water, TGS1-depleted cells exhibited a significant delay in subcutaneous tumour growth (Fig. 2b) and a marked reduction in tumour size at the experimental endpoint (Fig. 2c).

**Figure 2.**
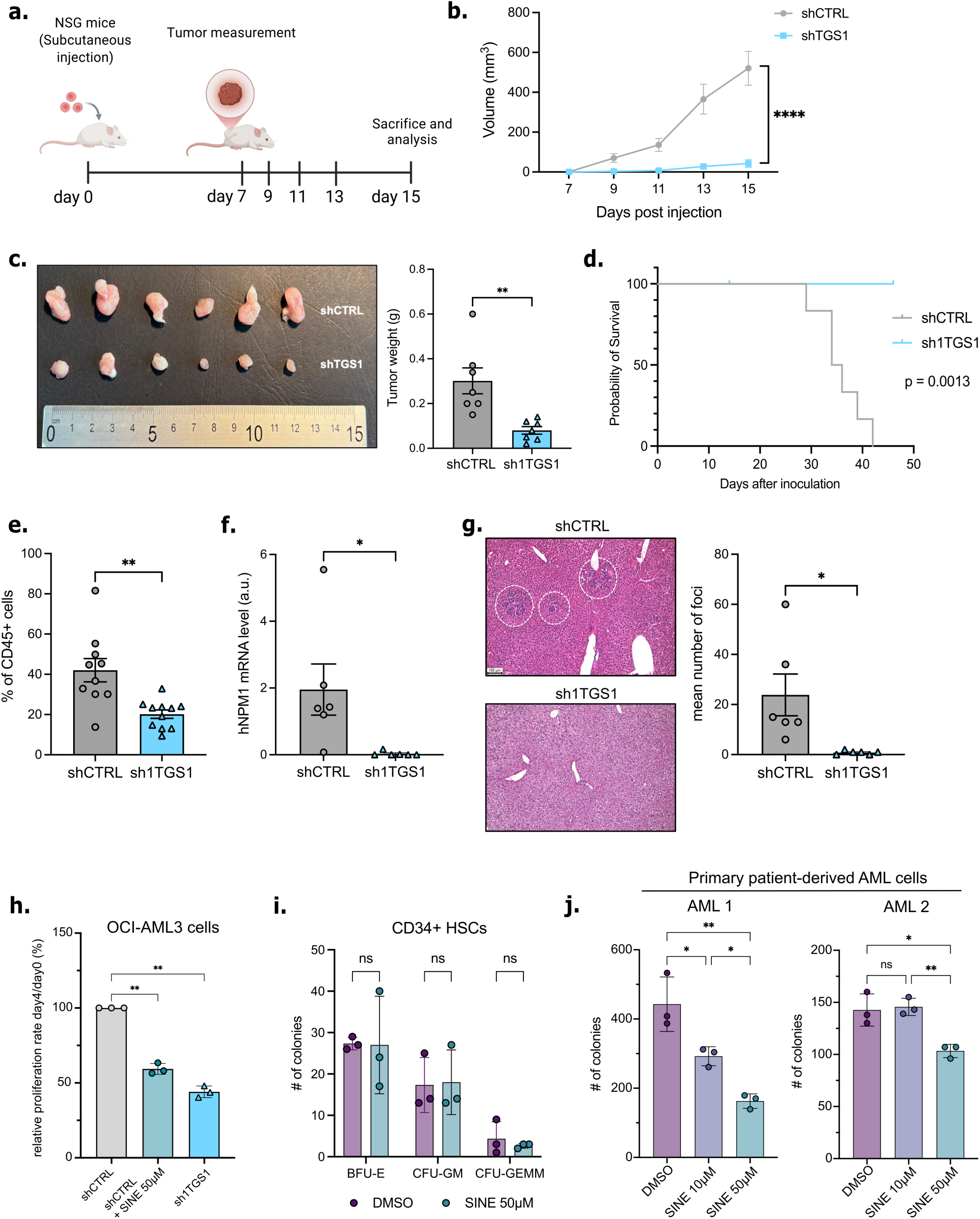
Loss of TGS1 impairs leukaemia cells’ growth in vivo and in vitroa. Schematic representation of the in vivo experimental design; tumour growth was monitored at the indicated time points. **b.** Tumour volume of mice injected with OCI-AML3 cells transduced with shCTRL or sh1TGS1, measured at the indicated time points. **c.** Representative images (left) of tumours collected 15 days after injection of shCTRL-or sh1TGS1-transduced OCI-AML3 cells and relative weight (right) of collected tumours. **d.** Kaplan–Meier survival curves of mice injected with shCTRL- or shTGS1-transduced OCI-AML3 cells (P = 0.013, Logrank Mantel-Cox test). **e.** Flow cytometric analysis of human CD45+ cells in the bone marrow of mice injected with shCTRL- or sh1TGS1-transduced OCI-AML3 cells. **f.** RT–qPCR analysis of human *NPM1* expression in the spleen of mice injected with shCTRL- or sh1TGS1-transduced OCI-AML3 cells. **g.** Hematoxylin and eosin (H&E) staining of liver sections showing myeloid infiltration and metastatic lesions in mice injected with shCTRL- or sh1TGS1-transduced OCI-AML3 cells. **h.** Proliferation assay of shCTRL and sh1TGS1 OCI-AML3 cells, measured between day 0 and day 4, for DOX treated cells, these time points correspond to 4 days and 8 days of treatment, respectively; shCTRL cells were also treated with SINE (50 µM) or DMSO, with fresh medium containing the respective treatments. **i.** Distribution of colonies across distinct myeloid lineages, including burst-forming unit-erythroid (BFU-E), multipotent progenitors (CFU-GEMM), and granulocyte-macrophage progenitors (CFU-GM), derived from human CD34+ HSCs treated with SINE (50 µM) or DMSO. Colony-forming units were manually enumerated on day 11 based on distinct colony morphologies. **j.** Total number of colony-forming units derived from AML patients’ blasts were quantified after 7 (AML 1) or 14 (AML2) days of culture with SINE (10 or 50) or DMSO treatment. Values are presented as mean ± SD of *n* = 3 independent experiments. *P<0.05, **P<0.01 and ****P<0.0001 unpaired two-tailed t-test was used in **b**, **c**, **e**, **f** and **g**, one-way ANOVA test with Dunnett’s (**h**) or Tukey’s (**j**) multiple comparison or two-way ANOVA test with Šidák’s multiple comparisons (**i**).

To further evaluate their engraftment capacity, shCTRL and sh1TGS1 OCI-AML3 cells were injected intravenously into NSG mice, which were then monitored for clinical signs of AML, including reduced hind limb mobility.

Survival was assessed in mice injected as described above and monitored for clinical deterioration. Mice receiving shCTRL cells began to die early, starting at 28 days post-injection (Fig. 2d), and exhibited classical signs of bone marrow infiltration, including a hunched posture and reduced posterior limb movement. In contrast, mice injected with sh1TGS1 cells showed no signs of leukaemia engraftment throughout the observation period (Fig. 2d).

Moreover, shCTRL OCI-AML3 cells demonstrated significantly higher engraftment efficiency in the bone marrow, as determined by flow cytometric detection of human CD45-positive cells (Fig. 2e), and in the spleen, as confirmed by RT-PCR analysis of the human *NPM1* gene (Fig. 2f). We also examined the ability of OCI-AML3 cells to infiltrate and form metastases in the liver. Mice injected with shCTRL cells displayed clear evidence of myeloid infiltration, whereas no liver metastases were detected in mice receiving TGS1-depleted cells (Fig. 2g). Collectively, these results demonstrate that TGS1 depletion profoundly impairs OCI-AML3 cell growth, reduces tumour engraftment, and effectively prevents leukaemia onset *in vivo*.

We extended our finding in primary CD34+ blasts derived from the peripheral blood of either healthy patients or AML patients at diagnosis (Supplementary Data 2). Since primary cells cannot be easily transduced or transfected, we used sinefungin to inhibit TGS1 catalytic activity. This SAM mimic was previously reported to have a selective and potent inhibitory effect on TGS1 at μM concentrations ^33,34^. Firstly, we validated the inhibitory effect of sinefungin on the proliferation of OCI-AML3 cells, showing effects similar to TGS1 conditional depletion (Fig. 2h). Next, we performed colony formation assay on CD34+ cells derived from healthy patients. In this context sinefungin did not affect vitality or colony forming potential of primary healthy blasts, as we didn’t observe any significant difference in the formation of erythroid (BFU-E), Granulocyte/macrophage (CFU-GM) or general myeloid stem cells (CFU-GEMM) (Fig. 2i). In contrast, sinefungin treatment induced a specific inhibition of colony formation potential for leukaemic blasts derived from two AML patients’ peripheral blood (Fig. 2j). A third patient’s primary blasts did not form colonies *in vitro* but sinefungin treatment affected vitality of leukaemic blast in a dose-dependent manner (Supplementary Fig. 2a). Taken together these results indicate that TGS1 catalytic inhibition predominantly affects the survival and colony formation potential of patient-derived leukaemic blasts.

### TGS1 directly methylates mRNAs encoding mitochondrial proteins

Previously known TGS1 targets include small nuclear non-coding RNAs and 11 out of 25 mRNAs encoding selenoproteins^7,10^. To better understand the molecular mechanisms downstream of TGS1 depletion in leukaemia cells, we performed RNA-immunoprecipitation (RIP) experiments followed by sequencing using a commercially available specific anti-2,2,7-trimethylguanosine antibody (m^2,2,7^G-RIP-seq). To avoid competition between highly abundant snRNA and other potential TGS1 targets, we performed the m^2,2,7^G-RIP-seq experiments separately on small and long (>200 nucleotides) RNA fractions following differential purification. Furthermore, since RIP experiments are prone to high background levels, we performed our experiments in both DOX-treated shCTRL and sh1TGS1 OCI-AML3 cell lines.

PCA analysis of m^2,2,7^G-RIP-Seq on short RNAs showed that RIP samples cluster separately from total (Input) RNA controls (Supplementary Fig. 3a) and that a specific enrichment was obtained (Supplementary Fig. 3b). Importantly, the majority of known TGS1 targets shows specific enrichment in our dataset, including *U1* and *U2* snRNAs, TERC and several different snoRNAs (Supplementary Data 3). Notwithstanding the above, we were surprised not to detect decreased enrichment, upon TGS1 depletion, of any modified short RNA, including all known TGS1 targets (Supplementary Fig. 3b). Our results were validated by m^2,2,7^G-RIP-RT-qPCR for several TGS1 small RNA targets (Supplementary Fig. 3c), and, since at the same induction time points, we already observed a quite remarkable phenotype, we speculated that these TGS1-dependent effects were unlikely to be mediated by the impairment of snRNA/snoRNA methylation and function.

Consistent with this view, PCA analysis performed on long RNA m^2,2,7^G RIP data revealed an evident clustering of biological replicates for both input and immunoprecipitated RNAs (Fig. 3a) and a decreased enrichment of immunoprecipitated RNAs upon TGS1 depletion (Fig. 3b). This not only indicates that m^2,2,7^G RIP is highly specific but also that TGS1 depletion significantly affects m^2,2,7^G levels on multiple long RNAs.

**Figure 3.**
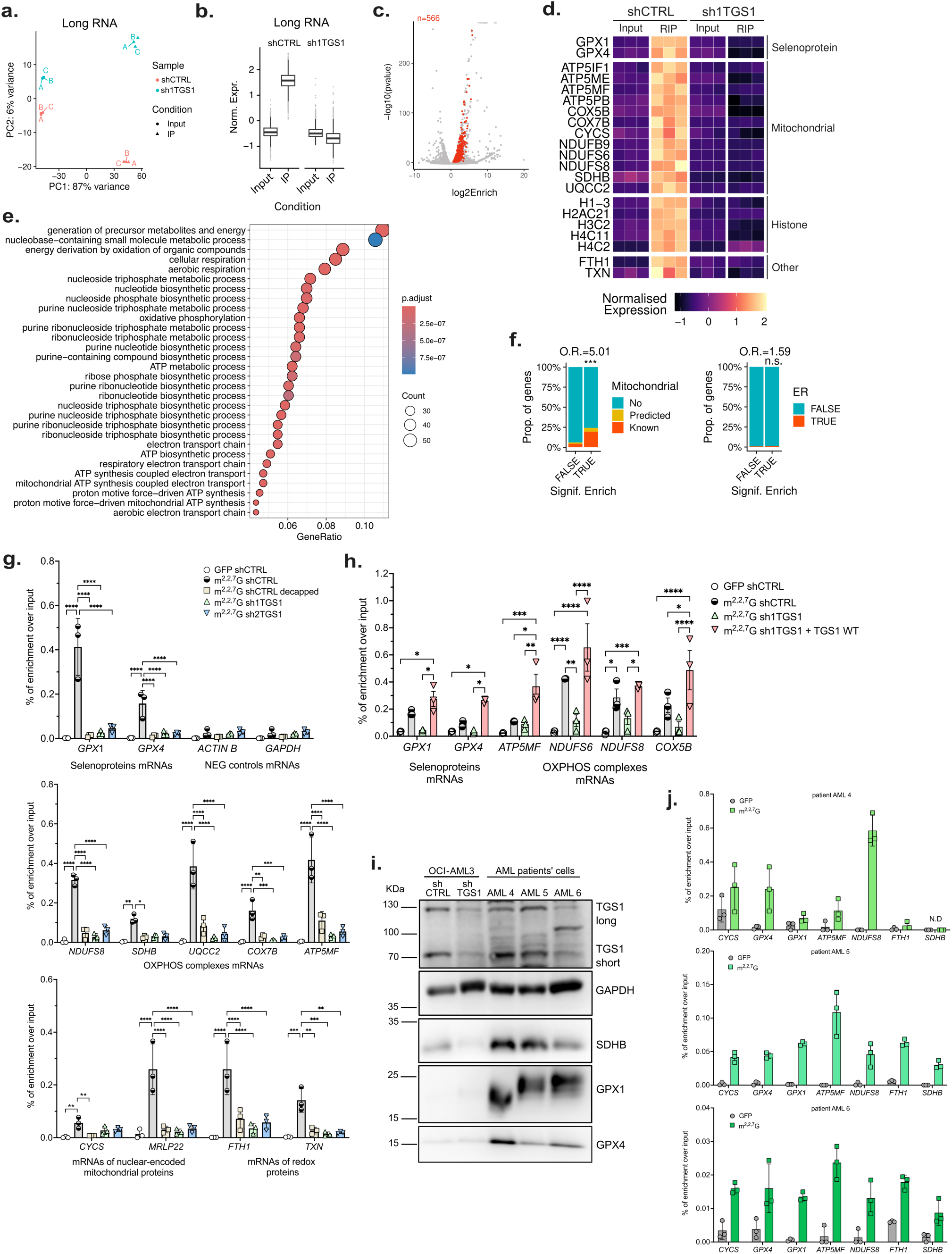
Newly identified TGS1 mRNA-targets detected in AML cell lines and patient samples. **a.** PCA analysis of the long RNAs m^2,2,7^G RNA-IP-Seq of both input and immunoprecipitations (IP) datasets of 3 independent biological replicates of shCTRL and sh1TGS1 OCI-AML3 cells after 6 days of DOX treatment. **b.** Normalized enrichment of m^2,2,7^G-modified long RNAs in shCTRL and sh1TGS1 OCI-AML3 cells after 6 days of DOX treatment. **c.** Volcano plot showing the Log2 enrichment over input of long RNA in shCTRL and sh1TGS1 OCI-AML3 cells after 6 days of DOX treatment. TGS1-dependent RNAs are highlighted in red. **d.** Heatmap showing the normalised expression of selected TGS1-target mRNAs in input and RIP samples from shCTRL or sh1TGS1 OCI-AML3 cells. **e.** Gene ontology analysis of TGS1-dependent m^2,2,7^G-modified mRNAs. **f.** Histograms showing the sub-cellular compartment localization of proteins encoded by TGS1-target mRNAs (TRUE) within mitochondria (left) or the Endoplasmic Reticulum (ER, right) compared with all other mRNAs (FALSE). **P<0.001, Fisher’s exact test, Prop. = proportion, O.R. = odds ratio. **g.** Bar plots showing m^2,2,7^G-RIP-RT-qPCR performed on long RNAs purified from shCTRL, sh1TGS1 and sh2TGS1 OCI-AML3 after 6 days of DOX treatment. Upper panel: selenoproteins and negative controls (not m^2,2,7^G-modified) mRNAs. Middle panel: mRNAs encoding OXPHOS complex subunits. Lower panel: nuclear mRNAs encoding mitochondrial proteins and mRNAs of proteins involved in redox processes. **h.** m^2,2,7^G-RIP-RT-qPCR was performed on long RNA purified from shCTRL, sh1TGS1 and re-expressing WT exogenous TGS1 (shRNA-resistant) sh1TGS1 OCI-AML3 cells after 6 days of DOX treatment. **i.** Immunoblot of total protein extracts from three AML patient samples and shCTRL and sh1TGS1 OCI-AML3 cells after 6 days of DOX treatment with the indicated antibodies. GAPDH was used as a loading control. **j.** m^2,2,7^G-RIP-RT-qPCR validation on long RNA purified from three AML patients (AML 4 top panel, AML 5 middle panel, AML 6 bottom panel). Values in **j** are means ± SD of three technical replicates. Values in **g** and **h** are means ± SD of *n*= 3 independent experiments; *P<0.05, **P<0.01, ***P<0.001, ****P<0.0001, two-way ANOVA test with Tukey’s multiple comparisons.

Transcriptomic analysis of both total input and immunoprecipitated RNA identified a set of TGS1-dependent m^2,2,7^G-modified RNAs with high-confidence. This set includes all genes displaying both: 1) a statistically significant enrichment in IP over input samples in shCTRL cells, and 2) a loss of IP enrichment upon TGS1 depletion (Methods).

This highly stringent analysis led to the identification of 566 previously unknown mRNAs carrying TGS1-dependent m^2,2,7^G cap hypermethylation (Fig. 3c and Supplementary Data 4a). Thus, TGS1-mediated cap hypermethylation of mRNAs extends far beyond the few previously reported selenoprotein mRNAs^10^. The enrichment levels of a selected subset of TGS1 targets are shown in Figure 3d. We next searched for common features at the 5’ end of TGS1 target mRNAs using HOMER, but this failed to identify a high confidence consensus sequence. Thereafter, to search for common functional features, we performed a gene ontology analysis of our newly discovered TGS1 targets. The analysis revealed strong enrichment of genes involved in mitochondrial metabolic pathways (Fig. 3e and Supplementary Data 4b), including mainly gene ontology terms involving cellular respiration, oxidative phosphorylation and ATP synthesis. Next, to confirm our results, we analysed the subcellular localization of proteins encoded by TGS1 mRNA targets combining three different annotation datasets. Our analysis showed a strong and significant enrichment in nuclear-encoded mitochondrial proteins amongst TGS1 targets (Fig. 3f). Precisely, mitochondrial proteins accounted for one third of TGS1 targets, the remainder being mainly endoplasmic reticulum (ER)-localised proteins and histones (Supplementary Data 4b).

To perform a more granular and structural analysis of TGS1-target mRNAs, we employed a recently developed approach to identify enrichment of sequence features combined with CLIP-seq dataset analysis. Since the m^2,2,7^G modification occurs at the 5′ cap, we focused our analysis on the 5′UTR sequences of TGS1-target mRNAs and performed a K-mer Enrichment Analysis (KEA) against a control set matched for 5′UTR length and mitochondrial representation (Methods). This analysis identified 49 7-mers with heterogeneous sequence composition significantly enriched in the 5′UTRs of TGS1 targets (Supplementary Fig. 3d and Supplementary Data 5). Importantly, applying the same analysis to the 3′UTR sequences revealed no enriched 7-mers, indicating that TGS1 target discrimination is primarily encoded by 5′UTR-localized sequence determinants (Supplementary Fig. 3d). Furthermore, as shown in Supplementary Fig. 3e, the enrichment detected by KEA was observed in both mitochondrial and non-mitochondrial TGS1 targets. Thus, the identified sequences represent a general feature of the TGS1 interactome rather than being driven simply by the subset of mitochondrial targets. To investigate the functional relevance of these 7-mers, we performed a K-mers RBP binding propensity analysis (K-RBP; Methods), which predicts Protein–k-mers associations based on eCLIP data^35,36^. The analysis of the TGS1-5’UTR 7-mers, grouped by nucleotide composition, revealed a strong and significant association with 17 known RNA-binding proteins (Supplementary Fig. 3f; Cliff’s delta ≥ 0.47 and Bonferroni-corrected p ≤ 0.05; analysis details in Methods). Notably, among them we found RBPs involved in the transport and maturation of mRNA encoding for mitochondrial protein, like LARP4^37^, U2AF^38^ and FASTKD2^39^. Collectively, our data support a model in which TGS1 selectivity is given by a combination of sequence determinants which are likely bound by a specific RBPs signature that favour TGS1 recruitment.

Next, we validated our sequencing dataset through m^2,2,7^G-IP-RT-qPCR experiments, including two previously known TGS1 targets encoding for selenoproteins, namely *GPX1* and *GPX4*, two negative controls and 20 of the newly identified targets (Fig. 3g and Supplementary Fig. 3g). Specifically, we included sveral mRNAs encoding for subunits of OXPHOS complexes *ATP5MF*, *NDUFS6*, *UQCC2*, *COX7B*, other mitochondrial proteins *CYCS*, *MRLP22* and proteins involved in redox homeostasis *FTH1* and *TXN*. Additionally, we also validated several histone encoding mRNAs (Supplementary Fig. 3g), such as *H1-3* and *H3C2*. Importantly, we further substantiated our findings by confirming the same TGS1 targets in sh2TGS1 cells, as well as in an additional AML cell line (Fig. 3g and Supplementary Fig. 4a). To highlight the specificity of our assay for 5’-cap-modification, we performed m^2,2,7^G-RIP-RT-qPCR on enzymatically de-capped long RNA and observed a complete loss of enrichment (Fig. 3g and Supplementary Fig. 3g). Finally, we expressed a shRNA resistant codon-optimized *TGS1* mRNA in both shCTRL and sh1TGS1 OCI-AML3 cell lines generating an overexpression model and a TGS1 rescue, respectively. m^2,2,7^G RNA immunoprecipitation performed in these cellular models showed that indeed TGS1 overexpression in CTRL cells could increase the methylation levels on TGS1 mRNA targets (Supplementary Fig. 4b) and that re-expression of the methyltransferase could efficiently rescue the loss of m^2,2,7^G on mRNA targets (Fig. 3h). Taken together, these experiments consistently validated the m^2,2,7^G-RIP-Seq dataset obtained in shCTRL and sh1TGS1 cells. Next, to determine whether TGS1 activity is conserved in primary human AML cells, we collected three samples derived from either bone marrow or peripheral blood of 3 AML patients at diagnosis (Supplementary Data 2). TGS1 expression in these cells was confirmed by immunoblotting, with particularly high levels seen in 2 out of 3 samples (Fig. 3i). Interestingly, the third patient showed lower TGS1 expression levels as well as a potential uncharacterised TGS1 isoform at approximately 100 KDa (Fig. 3i). We performed m^2,2,7^G-RIP-RT-qPCR on long RNAs isolated from the three AML samples plus an additional sample obtained from an AML-PDX model maintained *in vivo*^40^. Our analysis showed, for several of the newly identified TGS1 targets, consistent enrichment of m^2,2,7^G immunoprecipitations compared to a control anti-GFP antibody (Fig. 3j and Supplementary Fig. 3c). These data support the results obtained in primary AML patients’ blast (Fig. 2j and Supplementary Fig. 2a) and strengthen the possibility that the function of TGS1 is conserved between human AML cell lines and primary AML patients’ cells.

### Loss of TGS1 downregulates the expression of m^2,2,7^G-capped transcripts at protein level

Since the vast majority of TGS1 mRNA targets are not differentially expressed upon TGS1 downregulation, we concluded that m^2,2,7^G loss has likely no significant effect on either mRNA maturation or stability. Additionally, our RNA-seq Gene Ontology analysis did not identify any major alteration in splicing mechanisms between TGS1-depleted and control OCI-AML3 cells. In order to further investigate the phenotype of TGS1-depleted cells we performed a proteomic analysis of shCTRL and sh1TGS1 cells. Firstly, we performed a puromycin incorporation assay to assess the overall translation efficiency of shCTRL and sh1TGS1 cells reporting no significant difference (Fig. 4a). Next, we subjected shCTRL and sh1TGS1 OCI-AML3 cells to total proteomic analysis through mass spectrometry. Differential expression analysis revealed 973 regulated proteins upon TGS1-knockdown (Supplementary Fig. 5a), of which 625 upregulated and 348 downregulated (Fig. 4b and Supplementary Data 6a). While overexpressed proteins showed a similar GO enrichment to the upregulated transcripts previously identified (Fig. 3e), GO enrichment of downregulated proteins strikingly resembled the GO of the meRIP-identified m^2,2,7^G mRNAs (Fig. 4c, Supplementary Fig. 5c, d and Supplementary Data 6b, c, d). This analysis suggests that TGS1 targets are specifically downregulated at protein levels upon TGS1 depletion. The differential expression of TGS1-targets in the proteomic dataset confirmed that proteins encoded by both mitochondrial and non-mitochondrial TGS1-targets are indeed significantly downregulated in sh1TGS1 cells (Fig. 4d). Moreover, a significant correlation between downregulated proteins and m^2,2,7^G-mRNA modification was confirmed by additional orthogonal analysis (Fig. 4e). Overall, these proteomic data indicated that protein levels of TGS1-target mRNAs are specifically affected upon TGS1 downregulation, therefore suggesting that cap trimethylation might promote the translation of a subset of cellular mRNAs.

**Figure 4.**
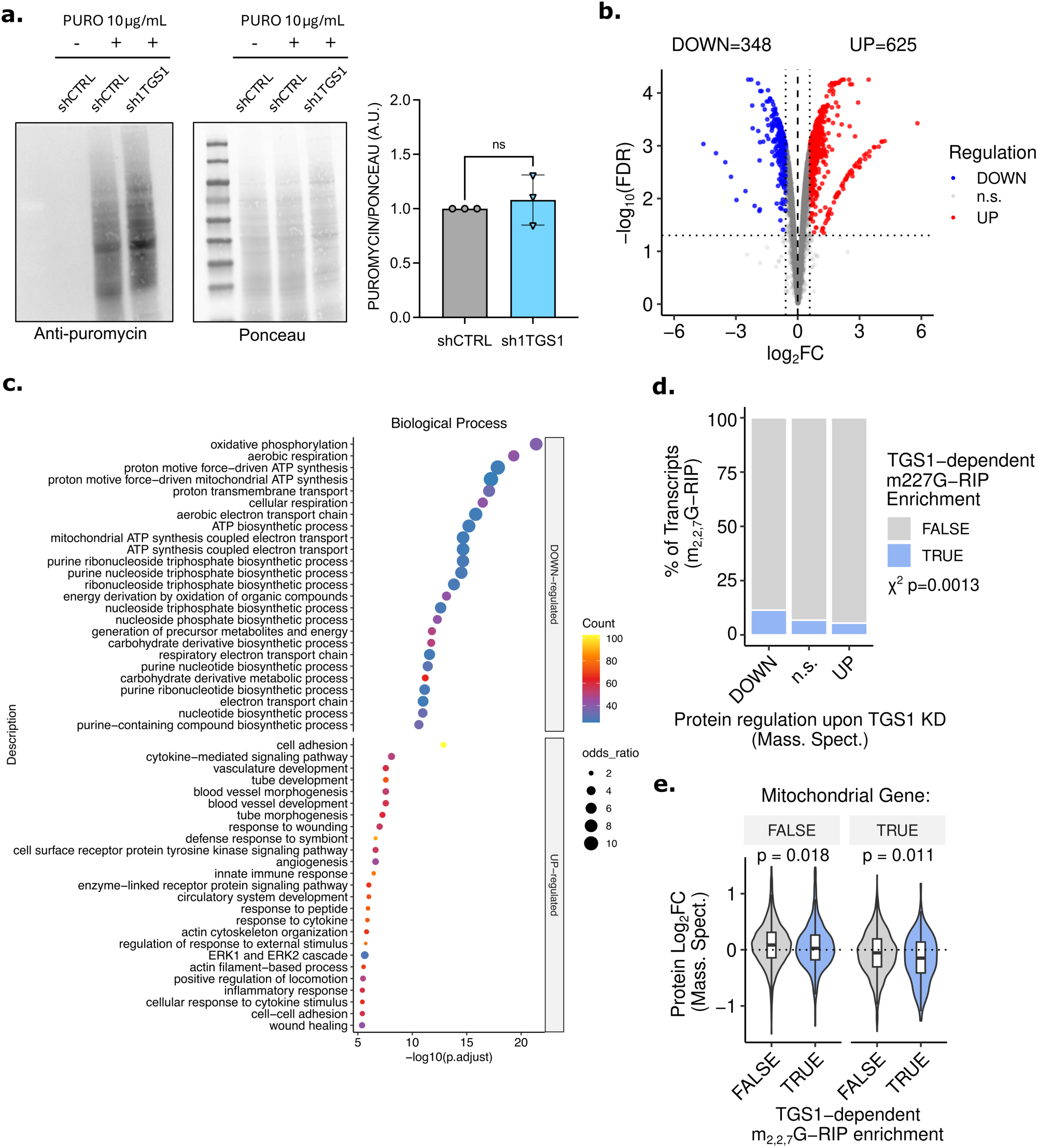
TGS1 knockdown selectively reduces the protein levels of TGS1 targets. **a.** The puromycin incorporation assay was performed in shCTRL or sh1TGS1 OCI-AML3 cells, followed by immunoblot analysis using an anti-puromycin antibody. shCTRL cells not incubated with puromycin were used as a negative control. Ponceau S staining was used as a loading control. **b.** Volcano plot of the log₂ fold change of differentially expressed proteins in OCI-AML3 cells following TGS1 knockdown. Upregulated proteins are highlighted in red, downregulated proteins in blue. Total proteomic analysis was performed by mass spectrometry on three independent biological replicates of doxycycline-induced shCTRL and sh1TGS1 OCI-AML3 cells. **c.** Gene Ontology (GO) analysis of Biological Processes terms associated with proteins downregulated or upregulated in sh1TGS1 compared with shCTRL OCI-AML3 cells (top 25 most significant terms are reported). **d.** The histogram reports the percentage of TGS1 targets amongst upregulated, not significant and downregulated proteins upon TGS1 depletion in OCI-AML3 cells. **e.** The violin plot reports differential expression of TGS1 non mitochondrial and mitochondrial targets compared to non TGS1 targets upon TGS1 depletion.

### Cap trimethylation promotes translation of mitochondrial OXPHOS subunits

To confirm and validate the proteomic analysis, we selected several proteins encoded by m^2,2,7^G-modified transcripts for further investigation, focusing mainly on mitochondrial-localised proteins.

Immunoblot analyses revealed significant downregulated expression of proteins encoded by several of the newly identified TGS1 targets (Fig. 5a, b), as well as selenoproteins, in both TGS1-KD OCI-AML3 (Supplementary Fig. 6a, b) and HL60 cells compared to shCTRL cells (Supplementary Fig. 6c, d). Specifically, SDHB, a subunit of the mitochondrial complex II was strongly decreased, together with CYCS (cytochrome *c*), TRX1 (thioredoxin 1) and ATP5I, (a structural component of the ATP synthase complex V). In contrast, protein level of an unmodified mRNA, SDHA, was not affected by TGS1-silencing (Fig. 5a). Dealing with mitochondrially localized targets, these differences were more apparent following cell fractionation experiments, which indeed showed a strong decrease in expression levels within the mitochondrial fractions of TGS1-depleted cells (Fig. 5c, d and Supplementary Fig. 6e, f). Importantly, downregulation of mitochondrial proteins did not appear to stem from a decrease in total mitochondrial mass in sh1TGS1 cells, as indicated by equivalent levels of the mitochondrial markers VDAC and TOM20 in control and TGS1-depleted cells (Fig. 5c, d and Supplementary Fig. 6e, f) and by MitoTracker-dye staining (Supplementary Fig. 6g). To further confirm these results, we performed immunoblot analysis upon TGS1 overexpression in shCTRL cells and upon rescue of TGS1 expression in sh1TGS1 cells, as previously described. Immunoblotting analysis revealed that TGS1 overexpression can increase the levels of nuclear-encoded mitochondrial proteins and that the re-expression of TGS1 can rescue the downregulation observed upon TGS1-knockdown (Fig. 5e). Next, we evaluated whether the TGS1 catalytic inhibition can phenocopy TGS1 depletion by treating OCI-AML3 cells with the methyltransferase inhibitor sinefungin^33^. Firstly, we confirmed that sinefungin treatment can indeed affect m^2,2,7^G levels on TGS1 targets, similarly to TGS1 depletion (Fig. 5f). Moreover, immunoblot analysis revealed that sinefungin treatment downregulates the protein levels of TGS1-targets, without affecting the expression of TGS1 (Fig. 5g). These results clearly demonstrated that the effects of TGS1-depletion on the expression of nuclear-encoded mitochondrial proteins is dependent on the deposition of m^2,2,7^G on their mRNAs through TGS1 catalytic activity.

**Figure 5.**
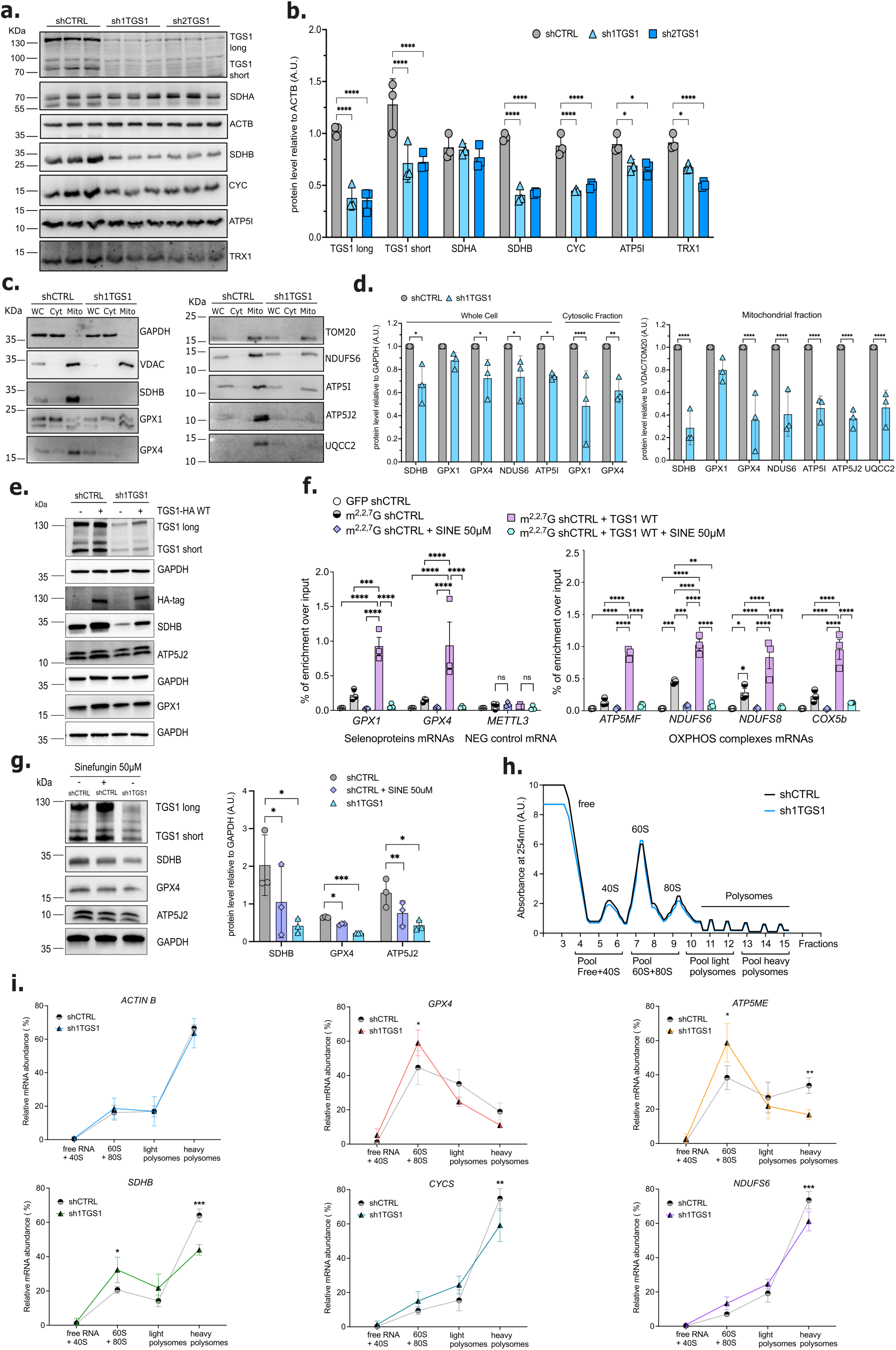
TGS1 silencing affects translation efficiency of modified targets. **a.** Immunoblot of total protein extracts from cells: shCTRL, sh1TGS1 or sh2TGS1 OCI-AML3 cells following 6 days of DOX treatment, using the indicated antibodies. A representative blot of three independent experiments is shown. **b.** Densitometric analysis of the immunoblotting shown in panel **a** and normalized to ACTB. **c.** Immunoblot of whole cell extracts (WC), cytoplasmic (Cyt) and mitochondrial (Mito) fractions purified from shCTRL and sh1TGS1 OCI-AML3 cells after 6 days of DOX treatment, using the indicated antibodies. A representative blot of three independent experiments is shown. **d.** Densitometric analysis of the immunoblotting shown in panel **c**. Each fraction was normalized using an internal control (upper panel: GAPDH for the WC and Cyt fraction; lower panel: VDAC or TOM20 for the Mito fraction). **e.** Immunoblot analysis of OCI-AML3 shCTRL and sh1TGS1 cells with exogenous expression of HA-tagged wild-type (WT) TGS1 after 6 days of DOX treatment, using the indicated antibodies. **f.** RT–qPCR bar plots showing enrichment of m^2,2,7^G-RIP over input performed on long RNA purified from OCI-AML3 cells, OCI-AML3 cells treated with SINE (50 µM), OCI-AML3 cells overexpressing WT TGS1, and WT TGS1-overexpressing OCI-AML3 cells treated with SINE. *METTL3* mRNA was used as a negative control. **g.** Immunoblot analysis of OCI-AML3 shCTRL and sh1TGS1 cells, untreated or treated with SINE (50 µM), using the indicated antibodies. Quantification of immunoblot signal intensities is shown on the right. **h.** Cell extracts from shCTRL and sh1TGS1 OCI-AML3 cells following 6 days of DOX treatment resolved in a 10–50% sucrose gradient. Absorbance at 254 nm was continuously measured. Peaks corresponding to free RNA, 40S and 60S subunits, 80S and polysomes are indicated together with the pooling strategy used for analysis shown in **i**. **i.** The indicated mRNAs in each pooled polysome fraction were quantified by RT-qPCR and plotted as a percentage of the total. Values in **b**, **d**, **f** and **i** are mean ± SD of *n* = 3 independent experiments, *P<0.05, **P<0.01, ***P<0.001 and ****P<0.0001, two-way ANOVA test with Tukey’s (**b**, **d** and **f**) or Šidák’s (**i**) multiple comparisons. Values in **g** are mean ± SEM of *n* = 3 independent experiments *P<0.05, **P<0.01, ***P<0.001, multiple paired t-test analysis.

To test whether TGS1-target mRNAs were differentially translated upon TGS1 depletion, we performed polysome fractionation experiments on control and TGS1-depleted OCI-AML3 cells. The fractionation profiles of shCTRL and sh1TGS1 cells did not show significant differences in overall levels of polysomes, nor in the relative abundance of ribosomal subunits (Fig. 5h). We then collected individual fractions and pooled them into 4 groups following the scheme outlined in Figure 5h. RT-qPCR analysis of TGS1 target mRNAs within pooled fractions highlighted a shift of these transcripts from the high polysomes to single ribosome fractions, indicating a decrease in their translation efficiency in sh1TGS1 cells (Fig. 5i and Supplementary Fig. 6h). In contrast, unmodified control mRNA *ACTIN Β* showed no discernible differences in its distribution between low and high polysome fractions upon TGS1 depletion. These data suggest that the lack of m^2,2,7^G modification significantly impairs translation efficiency and, ultimately, protein levels of mitochondrial oxidative phosphorylation components, potentially affecting cellular metabolism.

Inefficient translation of subunits belonging to the mitochondrial inner membrane complexes involved in the TCA cycle and oxidative phosphorylation could potentially affect the aerobic metabolism in human leukaemia cells, which are highly dependent on mitochondrial respiration to maintain energetic requirement for both their proliferation rates and their undifferentiated phenotype. We therefore assessed mitochondrial physiology in TGS1-depleted leukaemia cell lines.

### TGS1 depletion impairs mitochondrial respiration and causes oxidative stress in AML cells

Given that cap trimethylation supports the translation of mitochondrial OXPHOS subunits, we next assessed whether TGS1 depletion alters mitochondrial function by measuring the oxygen consumption rate (OCR) in sh1TGS1 and sh2TGS1 OCI-AML3 cells compared to shCTRL. Firstly, we performed a 6-hour time course using a Resipher oxygen flux analyser measuring a significantly decreased OCR over time in both TGS1 depleted cell lines (Fig. 6a). This result was independently validated using complementary approaches to evaluate mitochondrial oxygen consumption (Fig. 6b and Supplementary Fig. 7a). TGS1 depletion consistently resulted in a reduction of basal mitochondrial respiration across shRNA constructs (Fig. 6b and Supplementary Fig. 7a). Importantly, these differences were not dependent upon an intrinsic metabolic heterogeneity of our derived cell lines since no differences were detected before induction of the control and the TGS1-targeting shRNAs (Supplementary Fig. 7b). Our results confirmed that cellular respiration positively correlates with both TGS1 expression and catalytic activity in OCI-AML3 cells (Fig. 6c).

**Figure 6.**
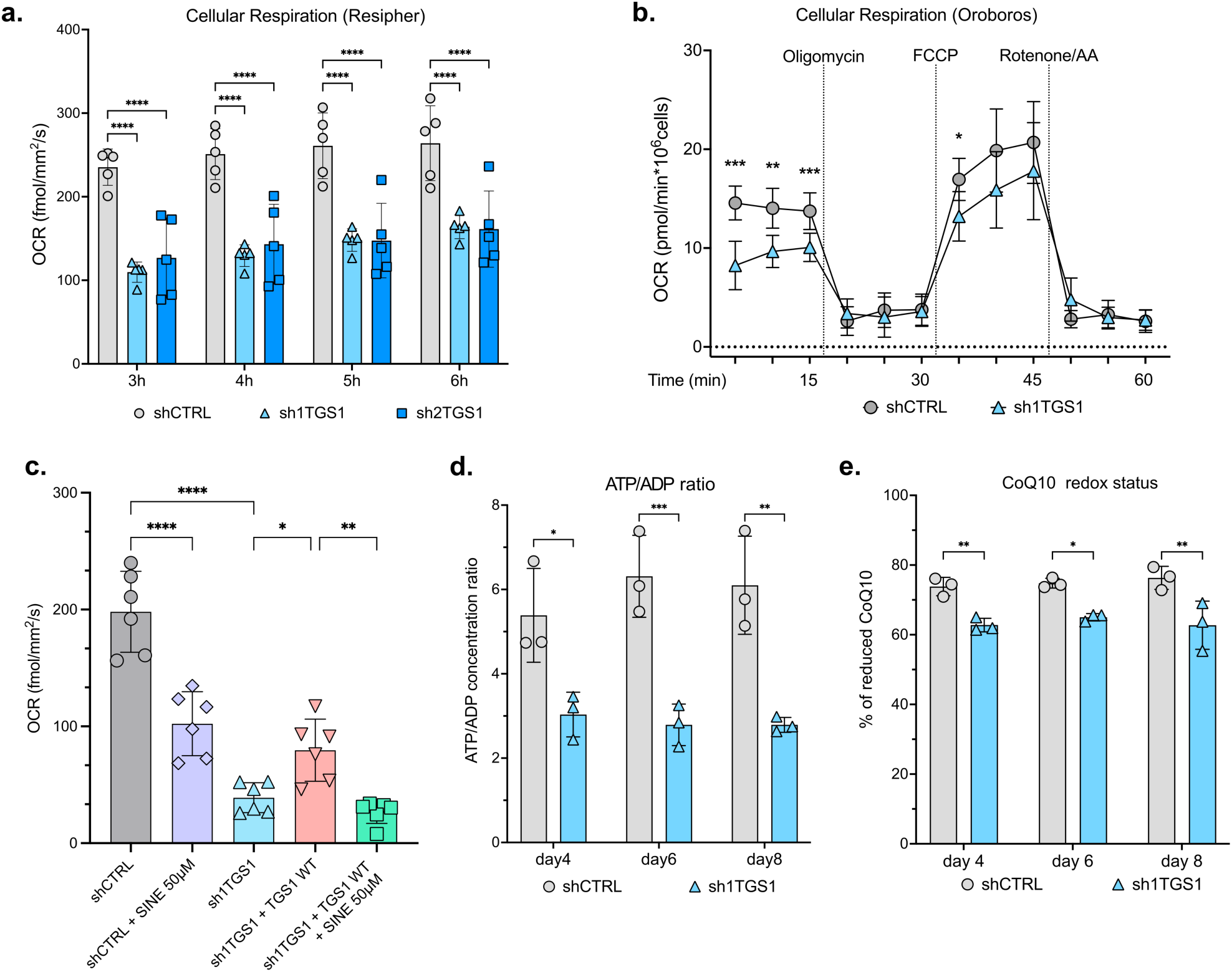
TGS1 depletion impairs cellular respiration in AML cellsa. Basal respiration of shCTRL, sh1TGS1 or sh2TGS1 OCI-AML3 cells following 6 days of DOX treatment was measured using the Resipher instrument at the indicated times (hours). **b.** High-resolution respirometry analyses of shCTRL or sh1TGS1 OCI-AML3 cells following 6 days of DOX treatment, detected by Oroboros. Basal respiration of cells was measured before oligomycin (ATP synthase inhibitor) treatment; maximal respiration of cells was measured in the presence of the uncoupler FCCP. The background was set after treatment with rotenone and antimycin A (AA). **c.** Basal respiration of shCTRL, shCTRL + Sinefungin 50µM, sh1TGS1, sh1TGS1 + TGS1 WT or sh1TGS1 + TGS1 WT + Sinefungin 50µM OCI-AML3 cells following 6 days of DOX treatment was measured using the Resipher instrument. **d.** Redox status of Coenzyme Q10 (CoQ_10_) measured by LC-MC in shCTRL or sh1TGS1 OCI-AML3 cells at the indicated days of DOX treatment (reduced CoQ_10_ over the sum of reduced and oxidised CoQ_10_). **e.** Bioenergetic ratio of shCTRL or sh1TGS1 OCI-AML3 cells at indicated days of DOX treatment estimated by measuring the ratio of ATP/ADP by LC-MS. Values in **a**, **b**, **c**, **d** and **e** are mean ± SD of *n* = 3 independent experiments. *P<0.05, **P<0.01, ***P<0.001 and ****P<0.0001, two-way ANOVA test with Šidák’s (**b**, **d** and **e**) or Tukey’s (**a** and **c**) multiple comparisons.

Since the majority of ATP generated in leukaemia cells is derived from aerobic respiration, we also assessed the changes in ATP to ADP ratio. Additionally, we applied an orthogonal approach and measured the reduction status of the CoQ_10_ pool. These analyses showed that ATP/ADP ratio was decreased (Fig. 6d) and the CoQ_10_ pool was significantly more oxidised (Fig. 6e) in OCI-AML3 cells upon TGS1 depletion, suggesting limited electron flow downstream in the respiratory chain. Together, our data highlight that TGS1 depletion has a significant impact on OXPHOS and ATP/ADP ratio in AML cells.

It is well established that slow rate of oxidative phosphorylation can lead to elevated levels of cellular reactive oxygen species (ROS)^41^, potentially impairing cellular function. We therefore directly measured cellular oxidative stress using the CellROX-dye staining and observed an increased percentage of dye-positive cells upon TGS1 depletion (Fig. 7a). We extended these findings by measuring the redox status of peroxiredoxin 3 (PRX3), a thiol-based peroxidase responsible for degrading mitochondrial hydrogen peroxide (H_2_O_2_)^42^. PRX3 undergoes reversible dimerization between its monomers via its reactive dithiol/disulfide active site that can be interrogated by immunoblotting. Our results show a shift of PRX3 towards the disulfide dimerized form upon TGS1 depletion (Fig. 7b). We also observed elevated levels of protein S-glutathionylation (Fig. 7b), a ROS-dependent post-translational modification of protein cysteine residues by glutathione disulfide (GSSG)^43,44^. This modification is suggestive of conditions characterised by oxidative stress and GSH oxidation to GSSG. Taken together, our data show that TGS1 depletion leads to decreased oxidative phosphorylation and increased oxidative stress in AML cells.

**Figure 7.**
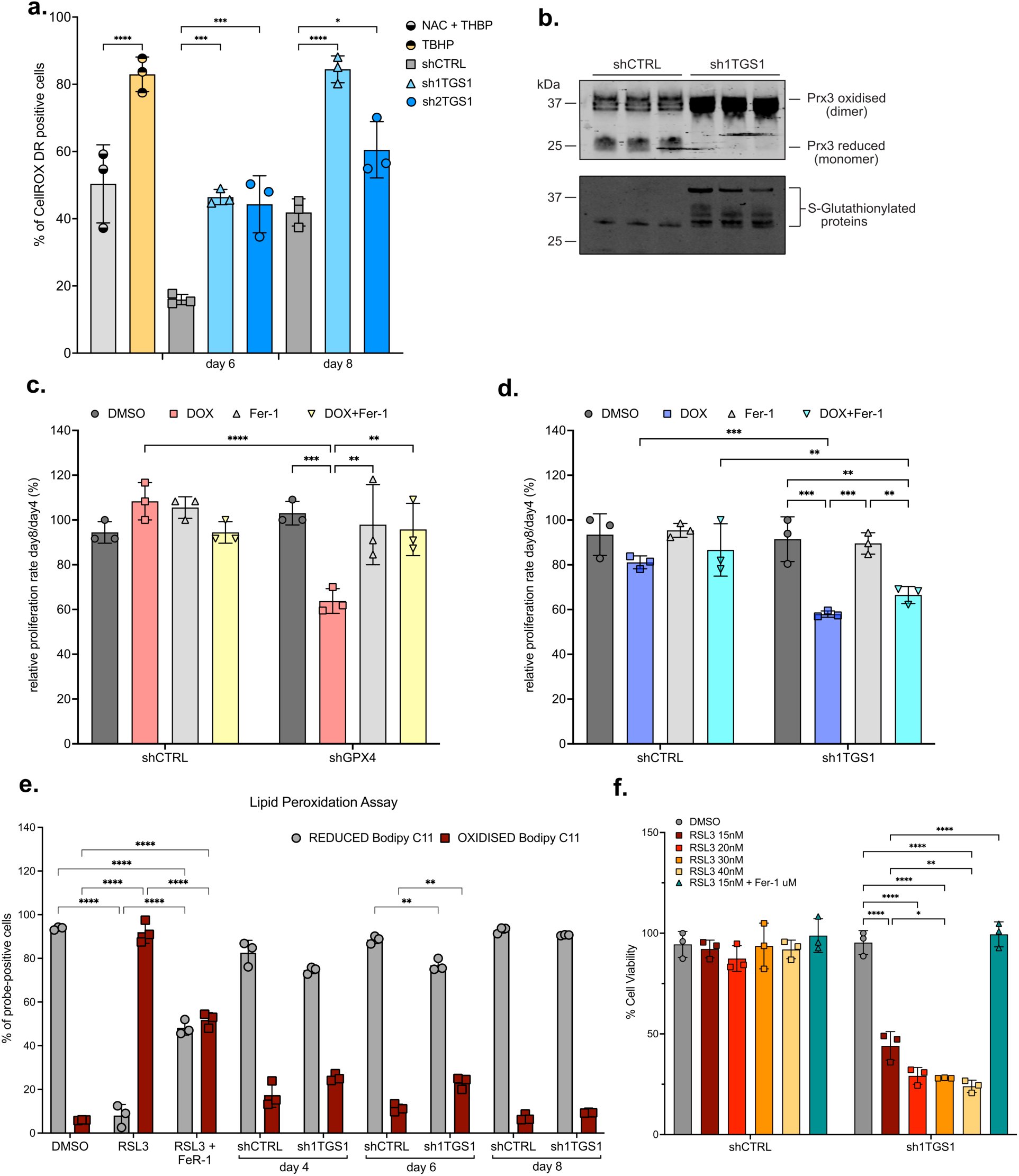
TGS1 knock-down induces accumulation of ROS and sensitise AML cells to low-doses of RSL3, a ferroptosis inducer. **a.** Flow cytometry measurement of reactive oxidative species (ROS) in shCTRL, or sh1TGS1 or sh2TGS1 OCI-AML3 cells, at the indicated days of DOX treatment, using CellROX™ Deep Red probe. N-Acetylcysteine (NAC) treatment of cells, followed by treatment with Tert-butyl hydroperoxide (TBHP), was used as a negative control. TBHP-treatment alone was used as a positive control. **b.** Levels of reduced and oxidised Peroxiredoxin 3 (Prx3) were analysed by immunoblotting of shCTRL and sh1TGS1 OCI-AML3 cells following 6 days of DOX treatment (upper panel), in 3 independent experiments. A semi-quantitative analysis of global protein S-glutathionylation, assessed via immunoblotting of the same samples, is shown in the lower panel. **c.** and **d.** Proliferation assay of shCTRL or shGPX4 (**c.**) or sh1TGS1 (**d.**) OCI-AML3 cells measured between day 4 and day 8 after DOX treatment, using a CellTiter-Blue® Assay kit. During the four days of the assay, cells were also treated with 5 μM Ferrostatin 1 (Fer-1) alone or together with DOX. **e.** Flow cytometry detection of the reduced and oxidised forms of the BODIPY™ 581/591 C11 probe in shCTRL or sh1TGS1 OCI-AML3 cells, at the indicated days of DOX treatment. Treatment for 2 hours with 0.5 μM RSL3 alone or in combination with 5 μM Fer-1 were used as positive and negative controls of lipid-peroxidation, respectively. **f.** Cell viability of shCTRL or sh1TGS1 OCI-AML3 cells following 5 days of DOX treatment and additional 24 hours treatment with the indicated concentrations of RSL3 (nM) alone or in combination with 5 μM Fer-1. Cell viability following exposure to DMSO (vehicle control) was used as the 100% viability reference for either shCTRL or sh1TGS1 cells, separately. Values in **a**, **c, d**, **e** and **f** are mean ± SD of *n* = 3 independent experiments. *P<0.05, **P<0.01, ***P<0.001 and ****P<0.0001, two-way ANOVA test with Šidák’s (**a** and **c**) or Tukey’s (**d**, **e** and **f**) multiple comparisons.

Recently, increased oxidative stress has been shown to be a trigger of ferroptosis^45^, with the role of selenoprotein GPX4 emerging as a major repressor of ferroptosis by preventing lipid peroxidation^46,47^. Furthermore, a number of newly identified TGS1 target mRNAs are also involved in regulating ferroptotic death, including *TXN*^48^, *SLC3A2*^49^ and *PDRX5*^50^. Since TGS1 inactivation drives oxidative stress, we speculated that TGS1 downregulation might also induce ferroptosis in AML cells.

To assess this, we first generated GPX4-depleted OCI-AML3 cells (Supplementary Fig. 8a), which showed decreased proliferation levels, comparable with those due to TGS1 depletion (Fig. 7c). The treatment with RSL3, a potent GPX4 inhibitor used to induce ferroptosis in cancer cells^47^, also decreased cell viability (Supplementary Fig. 8b), and this was associated with increased lipid peroxidation measured by C11-BODIPY staining (Supplementary Fig. 8c). However, while treatment with the antioxidant Ferrostatin-1 (Fer-1)^26^ was capable of rescuing cell proliferation in GPX4-depleted cells (Fig. 7c), it did not rescue the phenotype due to TGS1 knockdown in OCI-AML3 cells (Fig. 7 and Supplementary Fig. 8d). Additionally, TGS1 depletion on its own did not substantially increase levels of oxidised lipids, as instead induced by the treatment with RSL3 (Fig. 7e and Supplementary Fig. 8e). Thus, our results clearly demonstrate that TGS1 depletion alone does not trigger ferroptosis in AML cells, at least under our experimental conditions.

Next, we evaluated the ability of TGS1 depletion to sensitise leukaemia cells to ferroptosis induction by RSL3. We therefore treated OCI-AML3 cells, both shCTRL and TGS1 depleted, with a range of suboptimal doses of RSL3 and measured cell viability. Our results showed that, while there were no differences in overall sensitivity to high doses of RSL3 (Supplementary Fig. 8f), shTGS1 cells were significantly more sensitive to lower doses of RSL3 compared to shCTRL cells, with high levels of cell death already apparent at 15nM (Fig. 7f and Supplementary Fig. 8g). Importantly, sh1TGS1 cells treated with RSL3 underwent ferroptotic cell death since the effects of RSL3 were reversed by antioxidant Ferrostatin-1 treatment (Fig. 7f and Supplementary Fig. 8g). These data suggest that, while TGS1-depletion alone is insufficient to induce ferroptosis, it sensitizes leukaemia cells to ferroptosis inducing agents. Thus, TGS1 inhibition may represent a valid therapeutic strategy to induce oxidative stress in AML, either alone or in combination with ferroptosis inducers, to synergistically impair the hyperproliferative phenotype of leukaemia cells.

### TGS1 depletion affects local translation of mitochondrial-targeted proteins

The translation of nuclear-encoded mitochondrial proteins occurs in close proximity of the mitochondrial outer membrane, where newly synthesized proteins are rapidly imported inside mitochondria through a specific recognition peptide^22^. We hypothesized that TGS1-mediated cap hypermethylation might regulate the localization of nuclear transcribed mRNAs encoding mitochondrial proteins and that the observed decrease in translation could be due to the mis-localization of TGS1 target mRNAs. Recently, the development of the RNA-APEX-seq technique allowed the specific proximity labelling of mRNAs localized in different cellular compartments^51^. The technique was used to identify a subset of mRNAs localised at the outer mitochondrial membrane (OMM) (Fig. 8a). We therefore interrogated this set of OMM-localised mRNAs for overlap with our newly identified TGS1 mRNA targets. Despite the original APEX-seq dataset identifying OMM transcripts was generated in HEK-293T cells, we found a statistically significant overlap between the OMM-APEX dataset and TGS1-targets’ dataset from OCI-AML3 cells. Our analysis shows that indeed 175 TGS1 target mRNAs (31% of the total 566) also belonged to the OMM-APEX subset (Fig. 8a, P=3.912e-24). Importantly, these 175 overlapping mRNAs consisted mostly of nuclear transcribed mRNAs encoding mitochondrial proteins. To investigate the role of TGS1 in the localization of its target mRNAs through APEX2-system, we generated an inducible TGS1-KD HEK-293T cell line (Supplementary Fig. 9a). Firstly, we confirmed the protein downregulation of several TGS1 targets identified in OCI-AML3 cells (Fig. 8b). Next, we transfected both shCTRL and sh1TGS1 HEK-293T cells with two plasmids encoding APEX2 proteins, localized either in the cytoplasm or within the OMM (Supplementary Fig. 9b). We confirmed the correct localization of the APEX2 proteins by immunostaining (Supplementary Fig. 9c) and showed that TGS1 downregulation did not affect their expression (Supplementary Fig. 9d). After shRNA induction with doxycycline, we transiently treated HEK-293T cells with H_2_O_2_ to trigger peroxidase-based mRNA labelling and captured biotin-labelled mRNAs with streptavidin-beads. Finally, we measured the enrichment of several TGS1 targets at the OMM upon TGS1 depletion by RT-qPCR. In the original dataset, OMM-APEX enrichment was observed upon either cycloheximide (CHX) or puromycin (PUR) treatment^51^. While cycloheximide stalls ribosomes but does not disrupt ribosome-mRNA interaction, puromycin can disassemble the translation complex. Interestingly, while cycloheximide treatment showed OMM enrichment for mRNAs encoding for mitochondrial proteins in general, puromycin treatment only preserved OMM localization of mRNAs encoding for OXPHOS subunits^48^. To test whether TGS1 depletion might affect ribosome-independent OMM localization of OXPHOS mRNAs, we performed both CYTO-APEX-RT-qPCR and OMM-APEX-RT-qPCR upon puromycin treatment. Unexpectedly, we observed increased localization of TGS1 target mRNAs at the OMM upon TGS1 depletion following puromycin treatment (Fig. 8c). Importantly, localization of non-target negative control *FTO* didn’t show the same trend. We therefore concluded that m^2,2,7^G modification is not necessary for ribosome-independent recruitment of mRNAs to the OMM, at least as measured via the APEX2 technique.

**Figure 8.**
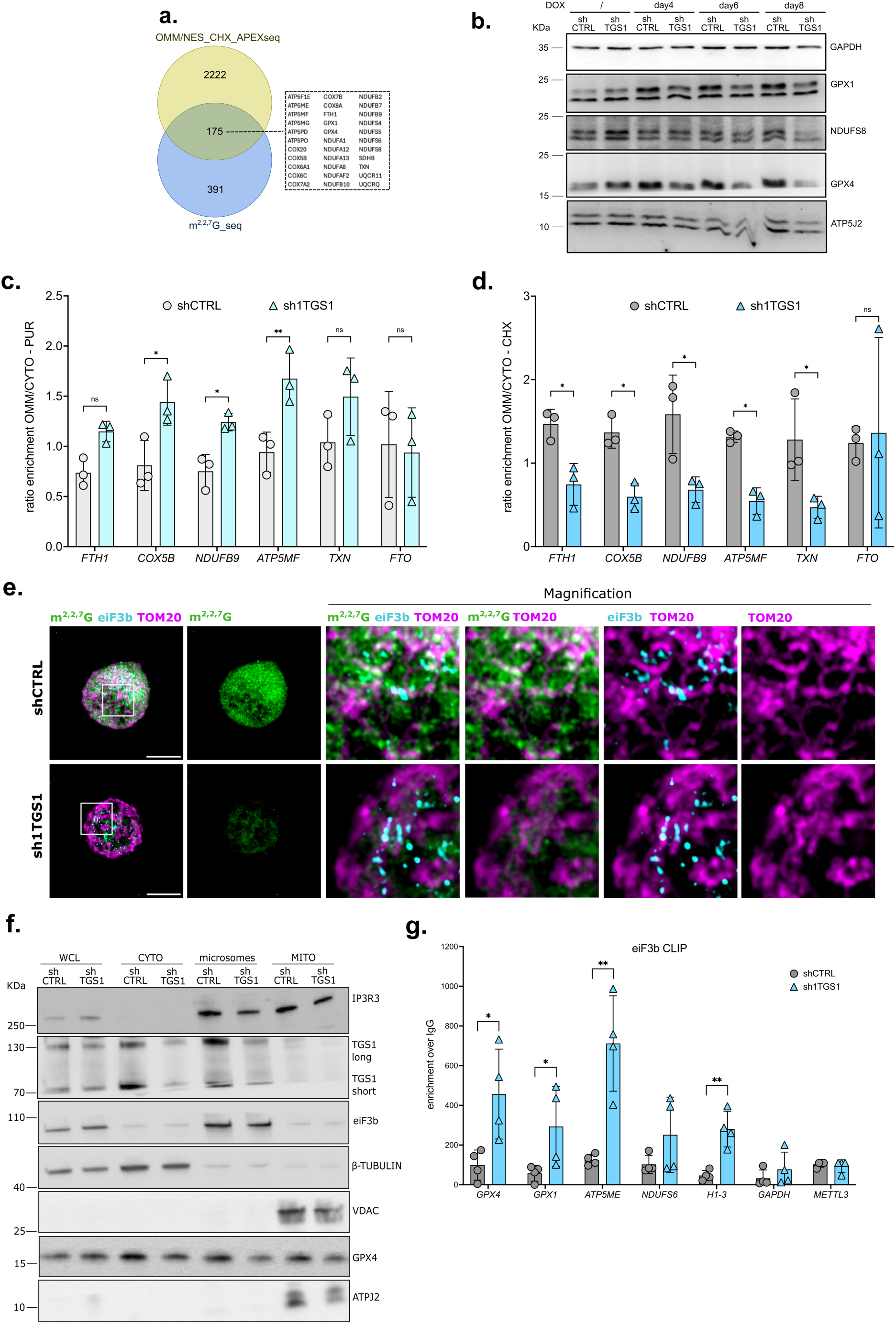
TGS1 regulates the OMM-localised translation of modified targetsa. Venn diagram of the overlap between TGS1 targets and outer mitochondrial membrane (OMM)-localised mRNAs identified in^42^ after cycloheximide treatment (P=3.912e-24, Fisher’s Exact Test). **b.** Representative immunoblot of total protein extracts from shCTRL or sh1TGS1 HEK-293T cells after the indicated days of DOX treatment. GAPDH were used as loading control. The images show a representative immunoblot of three independent experiments. **c.** and **d.** RT-qPCR analysis of streptavidin-captured biotinylated-RNA in shCTRL and shTGS1 HEK-293T cells expressing OMM-APEX2 or APEX2-CYTO treated with puromycin (**c.**) or cycloheximide (**d.**) The bar plots represent enrichment of the indicated targets in OMM-APEX cells over enrichment in APEX2-CYTO cells. Unmodified *FTO* mRNA was used as a negative control. **e.** Representative confocal images of immunofluorescence staining for eIF3b (light blue), m^2,2,7^G (green) and the mitochondrial marker TOM20 (purple) in shCTRL and sh1TGS1 HEK-293T cells. Scale bar = 10 μm (left panels) and indicated magnification (middle, right panels). **f.** Representative immunoblot of whole cell extracts (WCL), cytoplasmic (CYTO), microsomal (microsomes) and mitochondrial (MITO) fractions purified from shCTRL and sh1TGS1 OCI-AML3 cells following 6 days of DOX treatment. A representative immunoblot of three independent experiments is shown. **g.** CLIP-RT-qPCR on total RNA from DOX-induced (6 days) shCTRL or sh1TGS1 OCI-AML3 cells using a specific antibody against eIF3b. The bar plots represent the average enrichment over negative control IgG for the indicated TGS1 targets. Values in **c** and **d** are mean ± SD of *n* = 3 independent experiments. *P<0.05, **P<0.01 and ns = non-significant, two-way ANOVA test with Šidák’s multiple comparisons. Values in **g** mean ± SD of *n* = 4 independent experiments. P*<0.05, **P<0.01, one-tailed unpaired t-test.

Next, we performed APEX-RT-qPCR after treatment of cells with cycloheximide, to test the effect of TGS1 depletion on ribosome-dependent localization at the OMM. Under these conditions, we observed the opposite effect, a decreased OMM localization for TGS1 target mRNAs upon downregulation of the enzyme (Fig. 8d). Taken together, our results show that the recruitment of OXPHOS mRNAs targets at the OMM does not depend on TGS1. In contrast, TGS1 depletion may specifically affect the translation dynamics of OMM-localised nuclear encoded mitochondrial proteins.

It was previously reported that m^2,2,7^G-capped mRNAs are specifically bound by the translational initiation factor eIF3^52^. This multiprotein complex is necessary for translation initiation but is released from ribosome-mRNA complexes at later elongation stages. Firstly, we evaluated eIF3 localization through confocal immunofluorescence imaging using an antibody against one of the eIF3 subunits, eIF3b. Strikingly, eIF3b accumulated within cytoplasmic foci localized in close proximity to the OMM, stained using an anti-TOM20 antibody (Fig. 8e). Together, these data suggest that eIF3 localization marks the sites of OMM-associated translation of mitochondrial proteins. Importantly, our data showed that eIF3b localisation was not affected by TGS1 depletion, despite a substantial depletion of m^2,2,7^G modification within the cytoplasm, observed in doxycycline-induced sh1TGS1 cells (Fig. 8e). Accordingly, cell fractionation experiments confirmed EIF3b and TGS1 localization within the microsomal fraction and that TGS1 knockdown does not alter eIF3b levels (Fig. 8f).

Altogether, this suggests that TGS1 exerts its function at the ER-mitochondria interface to regulate translation, rather than affecting mRNA localization. Our data also show that eIF3b localization is not affected by TGS1 depletion, supporting the hypothesis that eIF3b localization is independent from m^2,2,7^G-cap hypermethylation. Despite previous reports indicating that the eIF3 complex binds 5’-cap-hypermethylated mRNAs, it was never shown whether its binding was actually dependent on m^2,2,7^G. To test this, we performed CLIP-RT-qPCR using an antibody against the eIF3 subunit eIF3b, in both shCTRL and sh1TGS1 OCI-AML3 cells. Our data show that, similar to our APEX-RT-qPCR analysis, eIF3b binding to TGS1 targets is significantly increased upon TGS1 depletion (namely GPX1, *GPX4*, *ATP5ME, NDUFS6* and *H1-3*), with no differences in eIF3b binding of control unmodified mRNAs *GAPDH* and *METTL3* (Fig. 8g). Collectively, our data further support the hypothesis that TGS1 and m^2,2,7^G affect translation dynamics of nuclear mRNAs encoding mitochondrial proteins. Specifically, they may facilitate the transition from initiation to elongation, and the release of eIF3 from the translational elongation complex.

Considering TGS1 and eIF3 complex microsomal co-fractionation, we tested the possibility that they may directly interact on the outer mitochondrial membrane. Proximity ligation assay performed with antibodies against TGS1 and eIF3b, eIF3d and eIF3e, showed no interaction and a relative weak signal for eIF3e and eIF3b, respectively, while a strong interaction was found for eIF3d in shCTRL OCI-AML3 and a significant decreased signal in sh1TGS1 cells (Fig. 9a and Supplementary Fig.9e). These findings open the possibility that TGS1 specificity for its mRNA targets could be mediated by its binding to eIF3d. Importantly, eIF3d was reported in previous studies as the main subunit of the eIF3 complex capable of interacting directly with mRNA 5’-Cap^53^.

**Figure 9.**
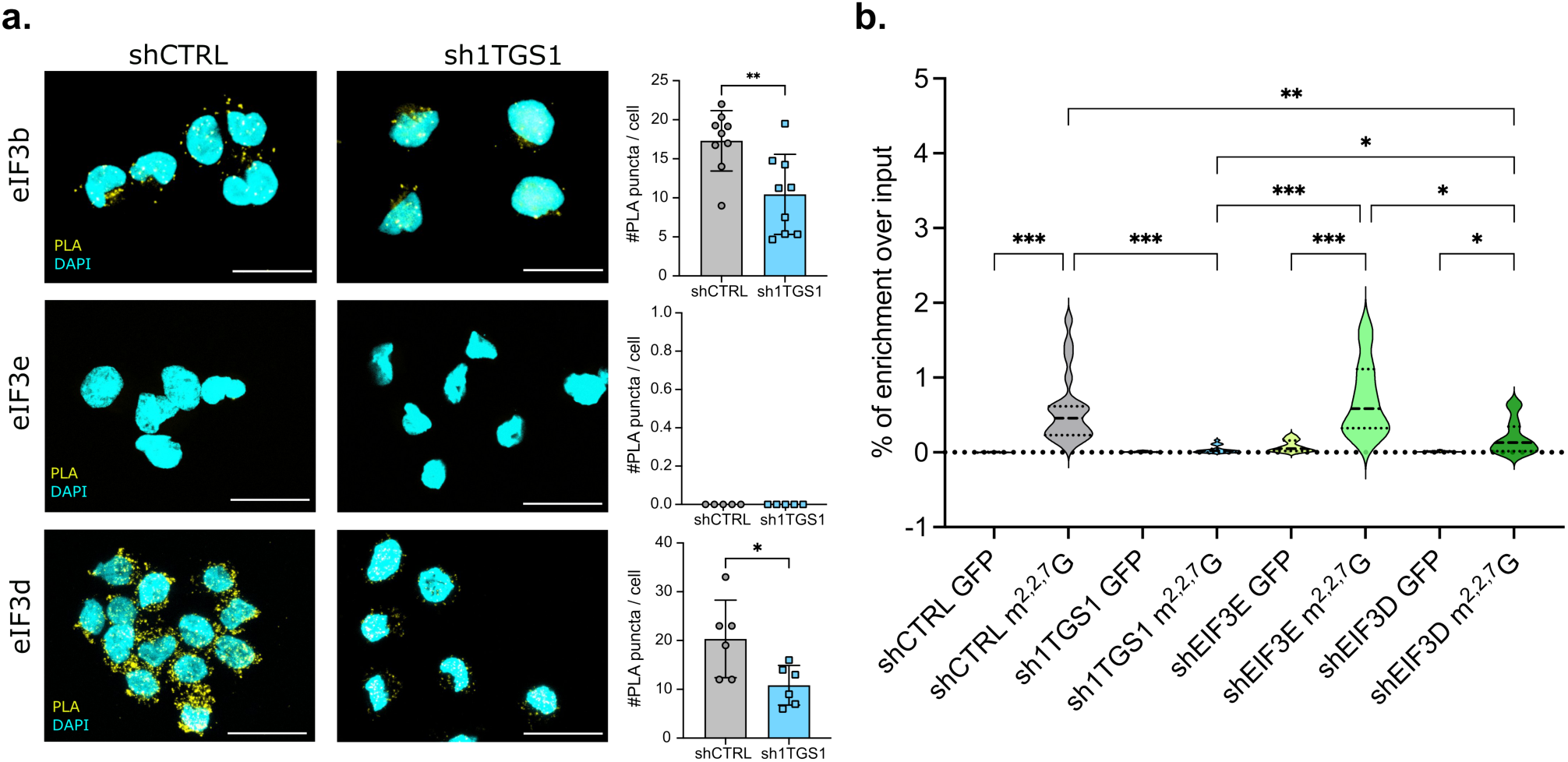
Interaction with eIF3 complex is required for TGS1 catalytic activity on mRNA targets. **a.** Representative Proximity Ligation Assay (PLA) images showing the interaction between TGS1, eIF3b, eIF3e and eIF3d. Yellow puncta indicate positive PLA signals, and nuclei are stained with DAPI (light blue). Scale bars represent 20 µM. Data are presented as mean ± SD from three independent experiments (n = 3). Statistical significance was determined using multiple unpaired t-tests; P*<0.05 and **P < 0.01. **b.** Violin plot showing the m^2,2,7^G-RIP-RT-qPCR performed on long RNA purified from shCTRL, sh1TGS1, shEIF3E and shEIF3D OCI-AML3 cells after 6 days of doxycycline treatment. An antibody against GFP was used as a technical negative control. Seven TGS1-targets were analysed by RT-qPCR, in *n* = 3 independent experiments. The dash thick and the dotted lines indicate the median and the quartiles of the data. Statistical significance was determined using one-way ANOVA test with Tukey’s multiple comparisons; P*<0.05, **P < 0.01, ***P < 0.001.

Next, we tested the hypothesis that the eIF3d complex itself might be required for the targeting of TGS1 on its specific substrates. We therefore generated EIF3B, EIF3D and EIF3E inducible-knockdown OCI-AML3 cells. While the knockdown of the core subunit eIF3b quickly led to cell death and we could not test this model, we obtained consistent depletion of eIF3e and eIF3d (Supplementary Fig. 9f). Next, we subjected these cell lines to m^2,2,7^G-RIP-qPCR, measuring the modification levels of 7 previously validated TGS1 targets. Whereas eIF3e depletion did not affect m^2,2,7^G levels, the knockdown of eIF3d strongly decreased the modification at levels comparable with TGS1 inactivation (Fig. 9b and Supplementary Fig. 9g). Taken together, our data identify eIF3d as a TGS1 interactor and, moreover, a potential recruiter of TGS1 on its target mRNAs encoding for mitochondrial protein. We thus propose a model in which the direct interaction with the eIF3 complex works as a docking site for TGS1 catalytic activity on these specific target mRNAs.

## Discussion

In this study, we have identified a new biological function of TGS1 in AML and uncovered a previously unreported molecular mechanism through which this methyltransferase exerts its function.

We have shown that *TGS1* is highly expressed in AML and this expression correlates with poor prognosis in refractory AML patients. Specifically, TGS1 is required to maintain high translational levels of key components of the TCA and oxidative phosphorylation metabolic pathways, in turn promoting high levels of mitochondrial respiration and preventing the accumulation of ROS. Conversely, TGS1 downregulation induces oxidative stress and a decreased oxygen consumption rate, leading to cell cycle arrest and differentiation of AML cells. These data are in keeping with the known dependency of AML cell lines on oxidative phosphorylation when compared to their untransformed counterparts^19,20^.

The relevance of metabolism-driven differentiation of leukaemia stem cells has been demonstrated in multiple studies^54^. Several small molecule inhibitors directly targeting mitochondrial metabolism, such as Tigecycline and IACS-010759, effectively arrest AML growth both *in vitro* and in AML animal models^20^. Despite this, recent phase I clinical trials of the Complex I inhibitor IACS-010759 for the treatment of both solid tumours and leukaemia were discontinued due to high toxicity adverse events^55^, indicating that direct targeting of mitochondrial OXPHOS complexes might not be a viable option for cancer therapeutics. Importantly, we demonstrated that decreased rates of OXPHOS and low ATP availability induced by TGS1 depletion can affect cellular fitness, while increased oxidative stress can activate differentiation programmes actively inducing cell cycle arrest. Importantly, these metabolic and phenotypic effects were recapitulated by catalytic inactivation of TGS1 using sinefungin, indicating that TGS1 enzymatic activity is required to sustain mitochondrial respiration and metabolic fitness in AML cell lines. Although further characterization is required, targeting TGS1 may represent an alternative strategy to effectively impair AML cell metabolism. Importantly, we showed that inhibition of TGS1 using sinefungin does not show marked toxicity against normal CD34+ haematopoietic precursors while compromising the clonogenic potential of primary leukaemic blasts *in vitro.* Whether TGS1 inhibition can elicit anti-leukaemic effects in human patients with an acceptable safety profile will need to be addressed in future studies.

High levels of ROS are associated with leukaemic transformation and can be involved in the activation of pro-oncogenic pathways^56^. On the other hand, it has been reported that further increase in ROS levels can quickly become detrimental to several different cancer types, including leukaemia^57^. While TGS1 downregulation alone disrupts AML cell lines growth without inducing a specific cell death mechanism, it can sensitise AML cell lines to RSL3, a clinically relevant ferroptosis inducer. Importantly, several factors involved in ferroptosis are TGS1-direct targets. In the absence of direct evidence, it can be hypothesized that impairment of TGS1 activity might increase cellular susceptibility to oxidative stress, potentially lowering the threshold for ferroptotic cell death. If this were the case, TGS1 inhibition could be envisioned as a possible adjuvant strategy to enhance the activity of pro-ferroptotic agents; however, this concept remains speculative and will require dedicated experimental validation.

In support of these considerations, and with the aim of translating our *in vitro* observations, our data indicate that silencing TGS1 in AML cell lines is associated with reduced tumour growth and decreased bone marrow replacement in recipient mice compared with control cells. Notably, these effects are accompanied by lower mortality in animals inoculated with TGS1-depleted cell lines, suggesting that TGS1 targeting may limit AML progression *in vivo* while being tolerated.

In our experimental settings, TGS1-depletion didn’t affect the m^2,2,7^G levels on the cap of snRNAs and snoRNAs. This might be due to our cellular models, where the short-term, inducible TGS1 depletion may not be sufficient to decrease m^2,2,7^G on nuclear non-coding RNA. Alternatively, our data open the possibility that a redundant catalytic enzyme might be responsible for m^2,2,7^G methylation of nuclear ncRNAs in AML cells. Further studies will be required to address this open question.

Focusing on long RNA, we identified more than 500 previously unknown mRNAs direct targets of TGS1 in AML cells. In particular, we discovered that more than 100 nuclear mRNAs encoding for mitochondrial proteins are m^2,2,7^G-modified and that TGS1, through its catalytic activity, is essential for their functional translation. Our investigation has uncovered TGS1 activity on mRNAs as a relevant mechanism in maintaining the proliferation of AML cells. Mechanistically, loss of TGS1 activity and m^2,2,7^G-5’-cap modification affects translational efficiency of proteins involved in OXPHOS. Translation of mitochondrial proteins specifically occurs in close proximity of the mitochondrial outer membrane and newly synthesised proteins are translocated inside the organelle in a tightly regulated manner. It is therefore likely that these translational mechanisms use specific epitranscriptomic modifications for their regulation. In support of this hypothesis, we showed that, although the recruitment of TGS1-target mRNAs at the proximity of the mitochondrial outer membrane is not affected by the loss of m^2,2,7^G, the dynamics of their translation is indeed impaired.

A recent publication demonstrated that the recruitment of transcripts encoding OXPHOS subunits at the OMM requires the RNA-binding protein AKAP1^58^. The authors suggest that AKAP1 may promote OXPHOS by increasing localized translation of ETC subunits. Our results introduce m^2,2,7^G-modification of OXPHOS mRNAs as a crucial player in this localised and complex translational regulation of nuclear encoded mitochondrial proteins. Importantly, AKAP1 protein is not modulated upon TGS1 silencing (Supplementary Data 6a) and recruitment of TGS1-targets at the OMM is not affected in TGS1-KD cells (Fig. 8c). Overall, our data support a model in which the mRNAs encoding for mitochondrial factors are recruited on the OMM through their interaction with AKAP1 and eIF3 while the m^2,2,7^G modification affects the translation dynamics of these mRNAs.

Mechanistically, while AKAP1 and eIF3 may be responsible for recruiting OXPHOS mRNAs near the outer-mitochondrial membrane, TGS1 and the m^2,2,7^G-5’-cap may be required to destabilize eIF3 binding to mRNA and promote efficient translational elongation. Another possibility is that a yet unidentified m^2,2,7^G reader mediates its function at mitochondria-associated translation sites. Further studies, aimed at fully dissecting the molecular mechanism, will be required to address these questions.

Importantly, it was reported that the eIF3 translational initiation complex binds m^2,2,7^G modified mRNAs but the recognition of mRNAs by eIF3 is mediated by specific 5’UTR structural conformations rather than the modification itself^52^. Interestingly, it was recently demonstrated that eIF3 is a global translational regulator of cellular metabolism, where differential binding of specific eIF3 subunits can promote translation of glycolytic or mitochondrial metabolic factors^59^. Our data, reporting a direct interaction between TGS1 and the eIF3 complex, highlight TGS1’s role as a key player in this peculiar, organelle-specific translation mechanism. In our proposed model TGS1 is recruited on its target through this interaction to depose m^2,2,7^G, in turn regulating the translational dynamic of its targets.

Recently, it was reported that the formation of specific RNA secondary structures, namely G-quadruplex, could mediate the localization and translation of mRNAs encoding for mitochondrial proteins^60^. Taken together these studies suggest that OMM-localized translation may be specifically regulated through the complex interplay of RNA binding proteins, epitranscriptomic factors and RNA secondary structures.

In summary, our data provide a comprehensive characterization of the role of TGS1 in AML cells. We have defined its biological role and identified the molecular mechanisms underlying its functions. Notably, these observations raise the possibility that TGS1 targeting could also be explored in combination with established AML therapies, such as cytarabine or Venetoclax, to potentially enhance anti-leukaemic effect. Future studies will be needed to determine whether modulation of TGS1 can influence treatment response and contribute to improved therapeutic strategies in AML.

## Material and Methods

### Cell lines

RN2c-Cas9, THP-1, and HEL cells were cultured in RPMI 1640 medium (Thermo Fisher Scientific, 31870025) supplemented with 10% FBS (Sigma-Aldrich, F9665) and 1% glutaMAX (Thermo Fisher Scientific, 35050061). HL60 cells were cultured in RPMI 1640 medium supplemented with 20% FBS and 1% glutaMAX. OCI-AML3 cells were cultured in MEMɑ (nucleosides GlutaMAX Supplement, Thermo Fisher Scientific, 22571-038) supplemented with 20% FBS. HEK-293T cells were cultured in DMEM (High Glucose, Thermo Fisher Scientific, 41965062) supplemented with 10% FBS and 1% glutaMAX. HEL and RN2c-Cas9 cells were grown in Ultra Low attachment flasks or plates (Corning, CLS3814, CLS347). All cell lines were incubated at 37°C, 5% CO2 and 100% humidity. All human cancer cell lines were obtained from the Sanger Institute Cancer Cell Collection, except OCI-AML3 which were received as a kind gift from Prof Brian Huntly’s lab (Cambridge, Stem Cell Institute) and routinely tested to be negative for mycoplasma contamination. The human cell lines used are not listed in the cross-contaminated or misidentified cell lines database curated by the International Cell Line Authentication Committee (ICLAC).

### Lentiviral vector preparation and cell transduction

For virus production, HEK-293T cells were transfected with the appropriate lentiviral vector (PLKO.1 for shRNA, LRG, LentiCRISPR-v2 for CRISPR-Cas9 and pLX304 for APEX2 constructs) together with the packaging plasmids PAX2 and VSVg at a 1:1.5:0.5 ratio by using Lipofectamine 2000 reagent (Thermo Fisher Scientific, 11668019), according to the manufacturer’s instructions. Supernatant was harvested 48h after transfection and filtered through a 0.45μm sterile filter (Sartorius Minisart, 16555K) before 2 × 10^6^ cells and the viral supernatant were mixed in a final volume of 2 ml volume of medium supplemented with 8 μg/ml (human) or 4 μg/ml (mouse) polybrene (Sigma-Aldrich, TR1003G), followed by spinfection (60 min, 900g, 32 °C) and incubation overnight at 37 °C. The medium was refreshed the following day, and the transduced cells were cultured further. A complete list of plasmids can be found in Supplementary Data 7a.

### Competition assays

RN2c-Cas9 cells were infected with LRG lentiviral vectors expressing GFP and a single gRNA targeting the catalytic domain of TGS1 or control region/genes. OCI-AML3, HEL, THP1 and HL60 cells were infected with LentiCRISPR-v2 lentiviral vectors expressing Cas9, GFP and a single gRNA targeting the catalytic domain of TGS1 or control regions/genes. A complete list of sgRNAs oligos (Custom Design, Sigma-Aldrich) can be found in Supplementary Data 4b. The percentage of GFP-positive cells (GFP+) was measured at day 3 after infection as a baseline by flow cytometry (LSR Fortessa cytometer, BD Bioscience). The percentage of GFP+ cells was measured again at day 11 after infection. The ratio of GFP+ cells at day 3 / GFP+ cells at day 11 was used as a measure of cells’ depletion. Two sgRNAs were used for TGS1, one sgRNA for the Rosa26 locus as a negative control, one sgRNA for RPA3 and one sgRNA for POLII as a positive control for mouse and human cells, respectively. The sgRNA sequences are listed in Supplementary Data 7b. Flow cytometry data were analysed using Flowjo software. Examples of gating strategy are reported in the Source Data.

### AML patients’ survival analysis

Transcriptomic data for the pediatric TARGET-AML project were obtained from the TCGA atlas using the TCGA biolinks package^61^ while transcriptomic data for the adult acute Myeloid leukaemia were obtained from the OHSU dataset^62^.

Patients were stratified according to *TGS1* expression, and survival trends of the patients belonging to the upper and lower 25% percentiles of TGS1 expression were compared using survminer (github.com/kassambara/survminer).

### Generation of conditional knockdown cells

OCI-AML3, HEL, HL-60, THP1 and HEK-293T cells were infected with pLKO-TET-on-Puro lentiviral vectors expressing shRNAs against the coding sequence of the specific targets or a scrambled control (Supplementary Data 7b). Medium was refreshed the day following transduction and cells were then treated with 2 μg/mL of puromycin (InvivoGen, ant-pr-1) for 6 days. In all experiments shRNA expression was induced through treatment with 100 ng/mL doxycycline hydrochloride (Sigma-Aldrich, D9891-1G) for the indicated times.

### Immunoblotting

Immunoblotting was performed as previously described^63^. For total cell protein extraction and western blot analysis, cells were lysed in a buffer containing 50 mM Tris-HCl pH 7.5, 150 mM NaCl, 1% Triton X-100, 0.5% deoxycholic acid, 0.1% sodium dodecyl sulphate, complete protease inhibitor cocktail (Roche, 11836153001) and cleared by centrifugation. For the isolation of nuclei from the cytoplasmic fraction, cells were incubated in hypotonic buffer (10 mM Tris-HCl, pH 7.8, 100 mM NaCl, 10 mM EDTA, 1.5 mM MgCl_2_) and the cytoplasmic membranes disrupted by adding 0.5% NP40. Pelleted nuclei were successively lysed in IPH buffer (50 mM Tris-HCl pH 8.0, 150 mM NaCl, 10 mM EDTA, 0.5% NP-40) for 10 min on ice and cleared by centrifugation. The protein concentration was determined by BCA Protein Assay (Thermo Fisher Scientific, 23225) or Bradford assay (Bio-Rad, 5000001). Resolved proteins were transferred to a 0.45 μm nitrocellulose membrane (GE Healthcare, GE10600002). Membranes were blocked with 5% non-fat milk in TBST (50 mM Tris-HCl pH 7.6, 150 mM NaCl, 0.05% Tween-20) for 1h at RT, and probed with primary antibodies overnight (Supplementary Data 7c). Membranes were then washed three times with TBST (10 min each) and probed with an HRP-conjugated secondary anti-rabbit/mouse/sheep antibody, as appropriate, for 1 h (Supplementary Data 7c). After three more washes in TBST, the signal was detected by ECL (GE Healthcare, RPN2134) and developed using an LAS-4000 Image Analyzer (Fujifilm/Raytek) or using a Chemidoc MP instrument (BioRad). Protein bands were quantified using ImageJ.

Cell lysates for assessment of protein S-glutathionylation and for estimation of the redox status of PRX3 protein were obtained from 1 x 10^6^ OCI-AML3 cells lysed in 500 mL RIPA buffer containing 1% of complete protease and phosphatase inhibitor cocktail (Roche 11836153001 and 4906845001) and 40 mM iodoacetamide, on ice for 15 min. After lysis, DNA was disrupted by resuspending the lysate using a 26G needle and 1 mL syringe. Subsequently, lysates were subjected to centrifugation (10 min at 17 000 x *g*, 4 °C) and protein concentration determined and adjusted to 1 mg / mL using the BCA Protein Assay. Lysate (40 μg of proteins) were mixed either with reducing or non-reducing 4X Laemmli sample buffer (Biorad, 1610747), denatured for 10 min at 45 °C and resolved at 170 V using reducing or non-reducing 4 - 20 % Novex Tris-Glycine gels (ThermoFisher) and NuPAGE MES SDS buffer (ThermoFisher). After electrophoresis, gels were electro-blotted onto a nitrocellulose membrane (22 mm) using a semi dry-transfer system (Biorad). Membranes were blocked using 5% non-fat milk in PBST for 1h at RT and probed overnight with corresponding primary antibodies (Supplementary Data 7c) in 1 % non-fat milk in TBST. Subsequently, membranes were washed three times with PBST and incubated with corresponding IR dye coupled IgG antibodies (Supplementary Data 7c) in blocking buffer (LiCOR) for 1h at RT in dark. After incubation, membranes were washed three times with PBST and signal was detected and quantified using Odyssey CLx imager (LiCOR) and Image Studio Light software (LiCOR).

### Proliferation assays

Cells (2 x 10^5^ / mL) were seeded into three separate 6 well plates and cultured for 4 days in media ± doxycycline (100 ng/mL) for 4 days. Cell proliferation was measured either by counting trypan blue stained cells using a Countess Cell Counting Chamber Slides (Invitrogen, C10228) and a Countess Automated Cell Counter (Invitrogen) or by CellTiter-Blue® Cell Viability Assay kit (Promega, G8081), according to the manufacturer’s instructions. CellTiter-Blue® fluorescence analysis was performed using the SpectraMax iD3 plate reader (Molecular Devices). Proliferation rate was analysed as the ratio between the initial number of seeded cells and the number of cells after incubation, normalised to untreated control cells set at 100% of proliferating cells.

### Cell cycle analysis

Cells (1 x 10^6^) were harvested on the indicated days following the start of doxycycline treatment and pellets were washed once with PBS. Cells were then permeabilized with ice-cold 70% ethanol, which was added dropwise under constant mixing with a vortex, for a minimum 30 minutes on ice. On the day of the assay, cell pellets were washed twice with PBS 1X and resuspended first in 50 μl of 100 μg/mL RNase solution (Roche, 11119915001) and then with 400 μl of 50 μg/mL Propidium Iodide solution (Sigma-Aldrich, P4170) in the dark for 10 minutes before transfer to a flow cytometry tube and assessment of the percentage of cells in each phase of the cell cycle by flow cytometry (LSR Fortessa cytometer and BD FACSCelesta, BD Bioscience). At least 30,000 events were collected per sample. Flow cytometry data were analysed using Flowjo software or BD FACSCelesta analysis software. Examples of gating strategy are reported in the Source Data.

### Apoptosis assay

Cells (2 x 10^6^) were harvested following 8 of doxycycline treatment and, for the positive control, after 24 h of 10μM Venetoclax treatment (MedChem Express, HY-15531-5MG). Cells were washed in PBS 1X and Annexin V/PI staining was performed using the eBioscience Annexin V Apoptosis Detection Kit APC, according to the manufacturer’s instructions (Thermo Fisher Scientific, 88-8007-74). Stained samples were kept for a maximum of 1 h on ice in the dark before flow cytometric analysis (LSR Fortessa cytometer and BD FACSCelesta, BD Bioscience). At least 30,000 events were collected per sample. Flow cytometry data were analysed using Flowjo software or BD FACSCelesta analysis software. Examples of gating strategy are reported in the Source Data.

### RT-qPCR

Total RNA from OCI-AML3, HEL, HL-60 or HEK-293T cells was purified using either the Direct-zol RNA MiniPrep Kit (Zymo Research, R2050) or the RNeasy mini kit (Qiagen, 74104), according to the manufacturer’s instructions. For gene expression evaluation, 500 ng of total RNA were retro-transcribed using the High-Capacity cDNA RT Kit (Applied Biosystems, 4368814). Reverse transcription (RT) reactions were diluted 1:10 and used as the template DNA for RT-qPCR using the PowerUp SYBR Green Master Mix (Thermo Fisher Scientific, A25780) on either the QuantStudio™ 6 Flex Real-Time PCR System or the ABI 7900 real-time PCR system. Double delta Ct (ΔΔCt) analysis was conducted with normalization to a selected control gene (ΔCt) and then relative expression levels were compared to the control sample. All RT-qPCR reactions were performed in three technical replicates. A complete list of primers can be found in Supplementary Data 7b.

### Mouse Experimental Protocols

#### Subcutaneous Injection Protocol

For subcutaneous administration, NSG mice were gently restrained, and injections were performed in the dorsal flank region using sterile technique. A total of 1 × 10⁶ cells were injected in a suspension consisting of 80% MEM and 20% Matrigel. Following injection, animals were monitored until full recovery and subsequently assessed daily for general health status, body weight, and signs of distress. Tumour growth was monitored using calipers, and tumour volume was recorded throughout the study period. After 24 days, mice were euthanized, and tumours were excised and weighed for further analysis. All procedures were approved by the Institutional Animal Care and Use Committee and were conducted in compliance with national and international regulations for animal welfare.

### Intravenous (i.v.) Injection Protocol

For intravenous administration, NSG mice were restrained, and injections were performed via the lateral tail vein under aseptic conditions. The compound was freshly prepared and administered in a volume adjusted according to body weight. A total of 0.5 × 10⁶ cells were injected in 10 µl of sterile PBS. Animals were monitored post-injection for adverse reactions and overall condition. All experiments were conducted in accordance with approved ethical guidelines and institutional protocols.

After 24 days, mice were euthanized, and flow cytometry (FACS) analysis was performed to assess the presence of human CD45-positive cells in the bone marrow. Spleens were analysed by RT-PCR to evaluate human NPM expression (Taqman Hs02339479_g1). Liver tissues were collected and analyzed for the presence of metastases. Liver sections were stained with hematoxylin and eosin (H&E) and evaluated for metastatic lesions using a DM6 microscope (Leica Microsystems).

### Doxycycline (DOX) Treatment via Drinking Water

Doxycycline (DOX) treatment was administered via drinking water. The DOX solution (0.6 mg/mL) supplemented with 5% sucrose was freshly prepared, protected from light, and water bottles were replaced every 2 days to ensure drug stability and consistent dosing. Water consumption was monitored throughout the treatment period to verify intake. Mice were observed daily for clinical signs of toxicity, changes in body weight, and alterations in general behaviour.

### Survival Curve Protocol

To evaluate overall survival and body weight changes, a total of 0.5 × 10⁶ cells were injected via the tail vein in 100 µL of sterile PBS. Mice were monitored daily from the start of treatment until reaching predefined humane endpoints or natural death. Survival time was recorded for each animal and used to generate Kaplan–Meier survival curves. Humane endpoints included significant body weight loss, impaired mobility, severe lethargy, or other clinical signs of distress, as defined by the approved protocol. Differences in survival between groups were analyzed using the log-rank (Mantel–Cox) test.

Experiments were carried out in accordance with the ethical guidelines of the European Communities Council Directive (2010/63/EU). Mice were housed in a certified animal facility under controlled environmental conditions: temperature 20–24 °C, relative humidity 40–60%, and a 12:12 h light/dark cycle. Animals were maintained in individually ventilated cages with 10–15 air changes per hour, with CO₂ levels kept below 3,000 ppm. Food and water were provided *ad libitum*. All animals were housed under specific pathogen–free (SPF) conditions. Animals were used in accordance with Animal Welfare Protocol No. VP_T8 CC652.172 EXT.

### Primary AML leukaemia samples

Primary cells from three AML patients (AML 1, AML 2, AML 3, AML4, AML5, AML6) were collected from peripheral blood or bone marrow, and isolated through a Ficoll gradient. All samples had a blast percentage >75% prior to Ficoll gradient purification. All patients gave informed consent for research use under ethical approval, for patients enrolled in the MetaboCAR study, under the approval of the Comitato Etico Territoriale Lombardia 3 (CET L3) (AML 1, AML 2, AML 3), or by the University of Perugia research ethics authority (AML 4, AML 5, AML 6; prot. #88471). Genetic characterisation of AML patient cells is reported in Supplementary Data 2. Cells from a *NPM1c/FLT3ITD/ITD* AML patient derived xenograft model (PDX2) were previously described in^33^.

### Colony-Forming Assay

CD34+ Hematopoietic Stem Cells (HSCs; Human Bone Marrow CD34+ Cells, Stemcell Technologies, 70002, LOT # 2508420015) cells were prepared as a single-cell suspension. Viable cells (3 x 10^4^) were plated in 35 mm Petri dishes and cultured for 11 days in MethoCult™ H4435 Enriched (Stemcell Technologies, 04435) prepared in Iscove’s Modified Dulbecco’s Medium (IMDM, Thermo Fisher Scientific, 12440053). Cultures were treated with Sinefungin (SINE, MedChemExpress, HY-101938-5MG) at 50 μM final concentration or with an equivalent DMSO dilution. Cultures were monitored throughout the assay period, and colony formation was evaluated by bright-field microscopy on day 11. Colonies were identified according to predefined morphological criteria (STEMCELL Technologies Technical Manual for Human CFU Assays), and the number of colonies formed in each condition was quantified.

After Ficoll gradient separation, AML blasts were counted and plated in 35 mm Petri dishes at 3,000 cells per dish in 1 mL of MethoCult™ H4434 (Stemcell Technologies, 04434), in the presence of sinefungin (at 10 µM or 50 µM final concentration) or the respective vehicle control (DMSO). Owing to the natural heterogeneity of AML blasts in colony formation, colonies were counted at 7 days (AML 1) or 14 days (AML 2 and AML 3) after treatment. All experiments were performed in *n* = 3 independent biological replicates and technical replicates for each condition.

### m^2,2,7^G-RIP

Total RNA from shCTRL, shTGS1, shEIF3D and shEIF3E OCI-AML3 cells, from shCTRL and shTGS1 HEL cells (30 x 10^6^), following 6 days of doxycycline treatment, and from AML 4, AML 5 and AML 6 cells (10 or 20 x 10^6^), was isolated using the Direct-zol RNA MiniPrep Kit (Zymo Research, R2050). Total RNA (20 or 30 μg) was then used to selectively purify long (> 200bp) and short (< 200bp) RNAs using the RNA Clean & Concentrator-25 kit, according to the manufacturer’s instructions (Zymo Research, R1018). For meRIP-RT-qPCR validation experiments in OCI-AML3 cells, 20 μg of the long RNA fraction was subjected to enzymatic de-capping using Cap-Clip™ Acid Pyrophosphatase, according to the manufacturer’s instructions (Cambio, C-CC15011H), and purified using the RNA Clean & Concentrator-25 kit. Long RNAs (5 μg) or short RNAs (1 μg) were incubated with 4 μg of anti-m^2,2,7^G antibody or anti-GFP antibody (Supplementary Data 7c) in a final volume of 1.2 mL of RIP buffer (10 mM Tris-HCl, pH 7.4, 150 mM NaCl, 0.5% NP40) supplemented with RNaseOUT (Thermo Fisher Scientific, 10777019), for 2 hours at 4°C. For the meRIP-RT-qPCR experiments performed on shEIF3E and shEIF3D OCI-AML3 and AML-primary cells’ RNA, 5 pg of m_3_^2.2.7^GP_3_G Trimethylated Cap Analog (Jena Bioscience, NU-853-1) were added into the RIP samples. Of the total amount of RNA used for the RIP, 5% was saved as the total input fraction (for both long and short RNAs). A protein G dynabead (Invitrogen, 1004D) slurry was washed in RIP buffer and pre-coated with 5 μg/μL BSA solution in RIP buffer for 2 h at 4°C. Subsequently, 80 μL of pre-coated bead slurry was added to the immunoprecipitation mix and further incubated at 4 °C for 2 hours. Dynabead-antibody-RNA complexes were washed 3 times with RIP buffer and immunoprecipitated RNA was eluted by competition using freshly made Elution Buffer (RIP buffer supplemented with 6.7 mM m_3_^2.2.7^GP_3_G Trimethylated Cap Analog), by shaking at 1100 rpm for 30 min at 37°C. Total input and immunoprecipitated RNA fractions were purified using the RNA Clean & Concentrator-5 kit, according to manufacturer’s instructions (Zymo Research, R1016), and eluted in 12 μL of nuclease-free water. Purified RNA was stored at - 80 °C until RNAseq-libraries were prepared for sequencing. For the meRIP-RT-qPCR validation experiments, a fixed volume of input and immunoprecipitated RNA were retro-transcribed, RT reactions were diluted 1:10 and used as the template DNA for RT-qPCR, as described above. Immunoprecipitated RNA enrichment over input was calculated as ΔCt and normalised to the % of total input. A complete list of primers can be found in Supplementary Data 7b.

### RNA library preparation and sequencing

Next generation sequencing RNA libraries were prepared using the SMART-Seq® Stranded Kit (TaKaRa Bio, 634443), according to manufacturer’s instructions. Briefly, RNA quantity and quality were determined using Qubit RNA HS Assay Kit and High Sensitivity RNA screen tapes (Agilent, 5067-5579) on the 4200 TapeStation System (Agilent). RNA libraries were generated from 500 pg of total input RNA or 7 μL of immunoprecipitated RNA, following the ultra-low protocol of the SMART-Seq® Stranded Kit. Libraries were barcoded using Indexing Primer Set HT for Illumina v2 (TaKaRa Bio, 634443) and finally, equi-molar multiplexed libraries were sequenced on the Illumina NovaSeq platform with paired-end 150 bp reads, at 5M (short RNA meRIP) or 15M (long RNA meRIP) raw reads per sample at Novogene (Cambridge Science Park, Cambridge, UK).

### RNA-seq data analysis

Transcript tags were quantified using Salmon^64^ in quasi-mapping mode using the Human GENCODE release 33 (ENCODE 99) annotation and summarized at the gene level with the tximport package^65^. We performed differential expression and enrichment analyses using DESeq2^65^.

Briefly, for m^2,2,7^G-RNA immunoprecipitation data, a generalized linear model was applied to fit count distributions, accounting for both sample treatment (Input and m^2,2,7^G-RIP) and source cell line (control and TGS1 knock-down). The drop in TGS1-dependent enrichment upon knock-down of the enzyme was evaluated using a negative binomial Wald test, with statistical significance defined as a Benjamini and Hochberg-adjusted p-value below 0.01 of the interaction term between sample treatment and cell line, under the alternative hypothesis of a decrease in enrichment upon TGS1 knock-down. Among the significant hits from this analysis, we selected only those transcripts that also exhibited a statistically significant RIP/Input enrichment > 0 in control cells. This criterion further strengthened the robustness of the analysis and ensured the exclusion of mildly enriched or unenriched transcripts showing even lower enrichment in the knockdown condition, as the interaction model alone would not take into account the initial amount of RIP enrichment in control cells.

In order to conduct gene expression analysis upon TGS1 depletion, we assessed gene expression fold changes and statistical significance of input RNA-seq data comparing shCTRL and shTGS1 cells, using these data as a proxy of total RNA levels.

### Gene Ontology analysis

Gene Ontology analysis was performed with the package clusterProfiler^66^ using the full list of expressed genes as a background universal set.

### Sequence feature analysis of TGS1 target transcripts

To perform RNA motif analysis, we extracted the first 100bp of the 5’UTR sequences from the full set of TGS1-modified mRNAs and analysed them using the findMotifs tool of the HOMER suite^67^.

Expressed genes (mean normalized RNA-seq count > 5) were annotated using GRCh38 Ensembl release 99, and mitochondrial protein-coding genes were identified using a MitoCarta-derived gene list. Each TGS1 mRNA target was matched to three expressed non-target mRNA controls by 5′UTR length and mitochondrial status, without reusing controls. Transcripts lacking valid 5′UTR annotations or sequences were excluded.

Complete 5′UTR sequences were analysed using K-mer Enrichment Analysis (KEA; D’Angelo Angelo, Ktools, GitHub, 2026 https://github.com/AngeloDAngelo/Ktools), identifying 49 enriched 7-mers supported by both enrichment procedures (empirical P < 0.01, nominal P < 0.05 and log₂ fold change ≥ 2; Supplementary Data 5). Motifs were grouped by nucleotide composition. Motif frequencies and log₂ odds ratios were calculated across meta-transcript bins spanning the 5′UTR, CDS and 3′UTR. The percentage of transcripts containing each motif family in the 5′UTR was also calculated separately for mitochondrial and non-mitochondrial targets and controls.

Candidate RNA-binding protein (RBP) associations were assessed using K-mer RBP binding propensity analysis (K-RBP; D’Angelo Angelo, Ktools, GitHub, 2026 https://github.com/AngeloDAngelo/Ktools), with an eCLIP-derived k-mer score matrix based on Kuret et al. (2022) and Van Nostrand et al. (2020)^35,36^. Each motif family was decomposed into constituent 5-mers and compared with the reference background (Mann– Whitney U test, Bonferroni-adjusted P ≤ 0.05 and z-score difference ≥ 0.47). Up to ten qualifying RBP-cell-line associations per family were displayed, ranked by z-score difference, with larger words indicating stronger predicted associations.

### Analysis of RNA localisation in Mitochondria/Endoplasmic Reticulum (ER)

The over-representation of mitochondrial and ER-localized transcripts were assessed by checking the IMPI^68^ evidence annotation status and presence of a "KDEL" amino acid motif, respectively, within the set of expressed putative TGS1-modified and unmodified transcripts. The statistical significance of the association was probed using a Fisher’s exact test.

### Protein LC-MS/MS and Proteomics data analysis

Cells (200 x 10^6^) were washed once with ice-cold 1× PBS and resuspended in lysis buffer (MIB; 220 mM mannitol, 70 mM sucrose, 10 mM HEPES, 1 mM EDTA, 1% Triton, supplemented with protease inhibitor cocktail, pH 7.4). Cells were mechanically disrupted on ice using a 10 mL Dounce homogenizer (Kimble Chase). Cell lysis efficiency, reaching approximately 80%, was monitored by transmission light microscopy.

Protein concentration was determined using the Bradford assay. For proteomic analysis, 20 µg of protein per sample were adjusted to a final volume of 60 µL and processed by in-solution tryptic digestion. Sample processing, including digestion and LC–MS/MS analysis, was performed by the EMBL Proteomics Core Facility (Heidelberg, Germany).

#### 1. Proteomic sample processing

Protein samples were subjected to the SP3 protocol^69^ conducted on the KingFisher Apex™ platform (Thermo Fisher). For digestion, trypsin was used in a 1:20 ratio (protease:protein) in 50 mM Triethylammonium bicarbonate (TEAB) supplemented with 5 mM Tris(2-carboxyethyl)phosphine hydrochloride (TCEP) and 20 mM 2-chloroacetamide (CAA). Digestion was carried out for 5 hours at 37°C. Residual magnetic beads were removed and the samples were dried by vacuum centrifugation.

#### 2. LC-MS/MS measurement

An UltiMate 3000 RSLCnano LC system (Thermo Fisher Scientific) equipped with a trapping cartridge (µ-Precolumn C18 PepMap™ 100, 300 µm i.d. × 5 mm, 5 µm particle size, 100 Å pore size; Thermo Fisher Scientific) and an analytical column (nanoEase™ M/Z HSS T3, 75 µm i.d. × 250 mm, 1.8 µm particle size, 100 Å pore size; Waters) was used. Samples were trapped at a constant flow rate of 30 µL/min using 0.05% trifluoroacetic acid (TFA) in water for 6 minutes. After switching in-line with the analytical column, which was pre-equilibrated with solvent A (3% dimethyl sulfoxide [DMSO], 0.1% formic acid in water), the peptides were eluted at a constant flow rate of 0.3 µL/min using a gradient of increasing solvent B concentration (3% DMSO, 0.1% formic acid in acetonitrile). The gradient was as follows: 2% to 8% in 6 minutes (min), 8% to 25% in 99 min, 25% to 40% in 5 min, 40-85% in 0.1 min, maintained at 85% B for 3.9 min and re-equilibrated to 2% B for 6 min. The outlet of the analytical column was directly coupled to an Orbitrap Exploris™ 480 mass spectrometer (Thermo Fisher Scientific) equipped with a Nanospray Flex™ ion source, operating in positive ion mode. Peptides were introduced into the instrument using a Pico-Tip emitter (360 µm OD × 20 µm ID; 10 µm tip, CoAnn Technologies) with a spray voltage of 2.5 kV. The capillary temperature was maintained at 275 °C. Full MS (MS1) scans were acquired in centroid mode over an m/z range of 420– 680, with a resolution of 60,000 at m/z 200 in the Orbitrap. The automatic gain control (AGC) target was set to ‘custom’ with a normalized AGC target of 300%. Data-independent acquisition (DIA) was performed across a precursor mass range of 430–670 m/z, using 4 m/z isolation windows. MS/MS scans were acquired in centroid mode over a scan range of 200–1800 m/z, with a resolution of 30,000. The AGC target for DIA was set to ‘custom’, with a normalized value of 3000%. A normalized higher-energy collisional dissociation (HCD) energy of 28% was applied for fragmentation.

#### 3. Data analysis

Raw files were analyzed using DIA-NN 1.8.1^70^ with directDIA module using an in-silico DIA-NN predicted spectral library (Trypsin as protease; C carbamidomethylation and N-terminal M excision; Protein N-terminal acetylation and Oxidation on Methionine enabled; 2 missed cleavages; precursor m/z range: 430-670; precursor charge range 2 to 4). The spectral library was generated from fasta files downloaded from Uniprot (Human: UniProt 2022 release, 20,311 sequences). The DIA-NN search used the following parameters: Precursor FDR (%) = 1; Mass accuracy, MS1 accuracy, and Scan window = 0; Use isotopologues; match between runs enabled; No shared spectra; Protein inference: Genes, Neural network classifier: Single-pass mode; Quantification strategy: Robust LC (high precision); Cross-run normalization: RT-dependent; Library generation: Smart profiling; Speed and RAM usage: optimal results.

DIA-NN output (report.tsv ) was processed to generate filtered results at the protein level for further downstream analysis. To ensure high data quality, the following filtering criteria were applied to the DIA-NN results: Q.Value <= 0.01, PG.Q.Value <= 0.01, and Lib.PG.Q.Value <= 0.01. Functions from the diann-rpackage (https://github.com/vdemichev/diann-rpackage/blob/master/R/diann-R.R) were employed to load, aggregate, and quantify proteins using DIA-NN’s MaxLFQ algorithm^70^. For proteins lacking gene annotations, the ’Protein.ID’ was used to fill in missing ’Gene’ entries, ensuring comprehensive gene-level representation. Also the ’PG.MaxLFQ’ quantity was used in these cases to fill the missing ’Genes.MaxLFQ.Unique’ entries. The filtered table, ’P4141_filtered_report.ProteinGroups_maxLFQ_matrix.tsv’, was generated. This matrix includes protein-level data, enriched with protein-level summaries, such as the number of Razor Peptides, Unique Peptides, Total Peptides, and the Total Intensity, which were calculated on the combined dataset and integrated into the gene-level output.

For the proteomics data analysis, the filtered DIA-NN output (maxLFQ quantification on gene level files) was processed using the R programming environment (ISBN 3-900051-07-0) to generate filtered results at the protein level for further downstream analysis. To ensure high data quality, the following filtering criteria were applied to the DIA-NN results: Q.Value <= 0.01, PG.Q.Value <= 0.01, and Lib.PG.Q.Value <= 0.01.

Functions from the diann-rpackage (https://github.com/vdemichev/diann-rpackage) were employed to load, aggregate, and quantify proteins using DIA-NN’s MaxLFQ algorithm^70^. For proteins lacking gene annotations, the ’Protein.ID’ was used to fill in missing ’Gene’ entries, ensuring comprehensive gene-level representation. Also the ’PG.MaxLFQ’ quantity was used in these cases to fill the missing ’Genes.MaxLFQ.Unique’ entries.

Initial data processing included filtering out contaminants and reverse proteins. Only proteins quantified with at least 2 razor peptides (with Razor.Peptides >= 2) were considered for further analysis. Additionally, only proteins identified and quantified in at least 2 out of 3 replicates for each condition were retained to ensure robustness. 7460 proteins passed the quality control filters.

Log2 transformed raw DIA intensities values (’Intensity’ columns) were normalized using the ’normalizeQuantiles’ function of the limma package. Missing values were imputed with the ’knn’ method using the ’impute’ function from the Msnbase package^71^. This method estimates missing data points based on similarity to neighbouring data points, ensuring that incomplete data did not distort the analysis.

Differential expression analysis was performed using the moderated t-test provided by the limma package^72^. The model accounted for replicate information by including it as a factor in the design matrix passed to the ’lmFit’ function. Imputed values were assigned a weight of 0.01 in the model, while quantified values were given a weight of 1, ensuring that the statistical analysis reflected the uncertainty in imputed data. Proteins with an FDR<0.05 were annotated as UP- or DOWN-regulated when their fold change was greater than 1.5 or lower than 0.67, respectively. These were used for downstream analyses (i.e. intersection with m2,2,7G-RIP-seq results and Gene Ontology enrichment).

Gene ontology (GO) enrichment analysis was performed using the ’compareCluster’ function of the ’clusterProfiler’ package^66^, which assesses over-representation of GO terms in the dataset relative to the background gene set. Enrichment was conducted for the following GO categories: Cellular Component (CC), Molecular Function (MF), and Biological Process (BP). The analysis was performed using ’org.Hs.eg.db’ as the reference database. The odds ratio (’odds_ratio’) for each GO term was calculated by comparing the proportion of genes associated with that term in the dataset (’GeneRatio’) to the proportion in the background set (’BgRatio’).

### Subcellular fractionation

Cells (200 x 10^6^) were washed once with cold PBS 1X before resuspension in mitochondrial isolation buffer (MIB) (220 mM mannitol, 70 mM sucrose, 10 mM HEPES, 1 mM EDTA, protease cocktail inhibitor, at pH 7.4). Cells were mechanically disaggregated using a 10 mL Dounce homogenizer (Kimble chase), on ice and cell breakage efficiency (around 80% of breakage) was checked using a transmission microscope. Whole cell fractions were saved, followed by centrifugations of the remaining extracts for 10 min at 800*g*, until no more nuclei and debris were observed. To obtain mitochondria, microsomal and cytosolic fractions, the supernatant was then subjected to centrifugation for 10 min at 2,500*g* to obtain the heavy mitochondria fraction. Pelleted mitochondria were then washed with MIB buffer and collected by centrifugation at 8,000*g* for 5 min, twice. The supernatant containing cytosol and microsomes, was subjected to ultracentrifugation at 100,000*g* for 60 min (Beckman Coulter Ultracentrifuge) to separate microsomes (pellet) and cytosolic (supernatant) fractions. All centrifugation steps were carried out at 4 °C. For western blotting analysis mitochondrial and microsomal pellets were resuspended in MIB containing 1% Triton and were heated at 50°C for 5 minutes.

From sh2TGS1 OCI-AML3 cells, mitochondria were isolated using the Mitochondria Isolation Kit for Cultured Cells (Thermo Fisher, #89874) in accordance with the manufacturer’s protocol.

### Puromycin incorporation assay

OCI-AML3 cells were incubated for 10 min at 37°C with 10 µg/mL of puromycin (Thermo Fisher Scientific, #A1113803). Thereafter, medium was replaced with fresh medium and after 50min of incubation at 37°C cells were harvested and processed for immunoblot analysis.

### Mitochondrial mass assessment by flow cytometry

Cells (1 x 10^6^) were collected by centrifugation for 5 min at 300*g* and resuspended in 1 mL of medium containing 15 nM MitoTracker™ Deep Red FM probe (Thermo Fisher Scientific, M22426), using a freshly made 15 μM stock solution of the probe. Following incubation for 30 min at 37 °C, stained cells were washed twice in warm PBS 1X, resuspended in 500 μL of PBS 1X containing 5 μl of 1 mg/mL 7-AAD staining solution (Thermo Fisher Scientific, A1310) and incubated for at least 10 min on ice before flow cytometry analysis (LSR Fortessa cytometer, BD Bioscience). MitoTracker Mean Fluorescence Intensity (M.F.I.) was recorded for at least 30,000 live cells (7-AAD negative cells) per sample. Flow cytometry data were analysed using Flowjo software. Examples of gating strategy are reported in the Source Data.

### Polysome fractionation

OCI-AML3 cells (15 x 10^6^) following 6 days of incubation with doxycycline (100 ng/mLl), were resuspended in 15mL of medium supplemented with 0.1 mg/ml of cycloheximide (Sigma-Aldrich, C7698) and incubated for 6 min at 37 °C. Cells were then washed twice in cold PBS 1X supplemented with 0.1 mg/mL of cycloheximide. Cell pellets were lysed in Polysome Extraction Buffer (20 mM Tris-HCl pH7.5, 100mM KCl, 5mM MgCl_2_, 0.5% NP-40), supplemented with protease inhibitor cocktail, RNaseOUT and 0.1 mg/mL of cycloheximide, for 10 min on ice with occasional, gentle tube-inversion. Lysates were subjected to centrifugation for 10 min at 12000*g* and the supernatant collected in a new tube. Finally, the supernatant was loaded into a freshly made 10-50% sucrose gradient (Fisher Scientific, S/8600/60) and subjected to ultracentrifugation for 90 min at 4 °C at 38,000 rpm, using the lowest deceleration force (Optima XE-90 Beckman Ultracentrifuge). Polysome fractions were collected using a Foxy® R1 Fraction Collector (Teledyne Isco) while measuring absorbance at 254 nm. The protocol was performed as reported in^73^ and experiments were run in four independent replicates. Fractions corresponding to free RNA+40S, 60S+80S, light polysomes and heavy polysomes were analysed separately, as reported in Figure 5h. RNA from each fraction was extracted using the Direct-zol RNA MiniPrep Kit (Zymo Research, R2050). A fixed volume of RNA was retro-transcribed as described above and relative RNA abundance in each fraction was quantified by RT–qPCR as a percentage of total RNA. A complete list of primers can be found in Supplementary Data 7b.

### Respirometry

The oxygen consumption measurements were performed in shCTRL and sh1TGS1 using a High-Resolution Respirometer Oroboros Instrument^74^ and intact OCI-AML3 cells. Approximately 2 × 10^6^ cells resuspended in 2 mL of MiR05 buffer were used for each measurement. Basal respiration was recorded until the steady state was reached. The non-phosphorylating respiration (LEAK) was measured by adding 2.5 μM oligomycin (Sellckchem, S1478) to the chambers to inhibit the ATP synthase, and the respiration rates were left to reach steady state. The uncoupled state or maximal capacity of the electron transfer system (ETS capacity) was achieved by titrating FCCP (Sellckchem, 8276) in 0.2 μM steps until the respiratory rates did not increase any further. Lastly, 1 μM antimycin A (Cell Signaling, 33357S) and 1 μM rotenone (Tocris 3616/50) were added to inhibit complex III and complex I, respectively. Oxygen consumption rate (OCR) measurements of shCTRL and sh2TGS1 cells were performed using a Seahorse XFe96 analyzer following the manufacturer’s instructions. The growth medium was replaced with assay medium (phenol red- and sodium bicarbonate-free DMEM [Corning, Cat. No. 90–013-PB]) supplemented with 1 mM pyruvate, 2 mM L-glutamine, and 10 mM glucose, adjusted to pH 7.4, and incubated in a non-CO₂ incubator. During the measurement, 1 μM oligomycin (Sigma, Cat. No. 495455), 2 μM FCCP (Sigma, Cat. No. C2920), and 1 μM rotenone/antimycin A (Sigma, Cat. Nos. R8875 and A8674) were sequentially injected into each well according to standard protocols. Three measurements of OCR were obtained following injection of each drug and drug concentrations optimized on cell lines prior to experiments. OCR rates were exported from Wave software (Agilent), normalized by cell count, and subsequently analyzed using GraphPad Prism v11.

Alternatively, OCR measurements of OCI-AML3 cells were performed using a Resipher instrument (Lucid Scientific), a multi-well dynamic oxygen consumption reader. Twenty-four hours before measurements were taken, cells were seeded at 5 x 10^5^ cells/mL to normalise for different proliferation rates. On the day of OCR measurements, 2 x 10^5^ cells/well were seeded in a Flat-Bottom 96 multi-well plate (Nunc™, 168055), in 100 uL of fresh medium containing 100 ng/mL doxycycline or not, in three technical replicates. Then, the Resipher oxygen sensing lid was attached to the plate and let to record for at least 12 hours (overnight). Since the Resipher measures OCR by analysing the medium oxygen concentration gradients within each well, to obtain accurate consumption analyses, these gradients need time to form, requiring a brief equilibration period after every plate movement. The equilibration time for our cells was set at 3h, and the ultra-sensitive oxygen sensors recorded the dissolved oxygen concentration gradient formed by metabolically active cells every 30 minutes for 8 hours. Data were analysed with Lucid Lab software (https://lab.lucidsci.com).

### LC-MS Analyses

#### Nucleotide quantification

OCI-AML3 cells (2 x 10^6^ cell / sample) were washed 2 times in ice cold saline (0.9% w/v NaCl) and collected by centrifugation (3 min at 300 x g). After centrifugation, the cell pellets were immediately lysed in nucleotide extraction buffer (80 % MeOH / 20 % H_2_O (v/v), containing 5 ⌠M of C_13_-labeled internal standards) using a Precellys lysing system (2 x 15 sec). After lysis, the supernatants were filtered using a PTFE filter (0.45 mm) and immediately analysed by LC-MS/MS. Analysis was performed using a SIL-300AC LC system coupled to a LCMS-8060 triple quadrupole mass spectrometer (Shimadzu, UK). The sample (5 ⌠l) was injected into a 50 ⌠L sample loop and separated by a HILIC column at 40 ^0^C using an Atlantis Premier BEH Z-HILIC (1.7 μm, 2.1 x 150 mm). The mobile phase was 15mM Ammonium Bicarbonate with 0.1 % Ammonium in DI H_2_O (A) and 100 % MeOH (B). The LC System’s Rinsing settings were: 50: 50 MeOH : MQ H_2_O, Rinse Type Internal & External, Rinse Port Liquid : R0, Rinse Mode : Before and after aspiration, Rise Dip Time 1 sec. Rinse Pump settings were: Rinse Method: Rinse port only, Rinse time 1sec. Internal Rinse On. The Starting gradient was 75 % B at 0.2 m / min. The Programming gradient was 0 to 0.5 min % B 75 keeping an isocratic gradient, 0.5 to 1.5 min % B 75 to 20, 1.5 to 4 min % B 20 keep isocratic gradient, 3.99 to 4 min % B 20 to 75, 4 to 10 min % B 75 keeping isocratic gradient. The mass spectrometer was operated in positive electrospray ionization with multiple reaction monitoring (MRM). MS Setting used for quantification were DL Temperature: 230 °C, Nebulizing Gas Flow 3 L / min, Heat Block Temperature 400 °C, Heating Gas Flow 10 L / min, Drying Gas Flow 10 L / min, Interface Temperature 300 °C. Desolation Temperature 526 °C. MS Setting used for quantification were: Adenosine 5′-(tetrahydrogen triphosphate) 507.9 → 136.05 with Dwell 100 msec Q1 -24V, CE -34, Q3 -29V as a major Ion 507.9 → 409.95 with Dwell 100 msec Q1 -24V, CE -18, Q3 -19V and 507.9 → 427.95 with Dwell 100 msec Q1 -24V, CE -21, Q3 -20V as a reference Ion; Adenosine 5′-(trihydrogen diphosphate) 427.8 → 136.05 with Dwell 100 msec Q1 -28V, CE -27, Q3 -28V as a major Ion 427.8 → 348.05 with Dwell 100 msec Q1 -20V, CE -19, Q3 -16V and 427.8 → 119.02 with Dwell 100 msec Q1 -12V, CE -55, Q3 -26V as a reference Ion; 5′-Adenylic acid 347.95 → 136.05 with Dwell 100 msec Q1 -17V, CE -19, Q3 -13V as a major Ion 347.95 → 151 with Dwell 100 msec Q1 -24V, CE -28, Q3 -27V as a reference Ion; C13 Adenosine 5′-(tetrahydrogen triphosphate) 516.5 → 141.1 with Dwell 100 msec Q1 -26V, CE -34, Q3 -27V as a major Ion 516.5 → 140.05 with Dwell 100 msec Q1 -26V, CE -33, Q3 -29V, 516.5 → 419.05 with Dwell 100 msec Q1 -26V, CE -18, Q3 -20V and 516.5 → 437.05 with Dwell 100 msec Q1 -26V, CE -21, Q3 -30V as a reference Ion.

#### CoQ redox status

Quantification of redox status of UQ_10_ was performed as described previously^75^. OCI-AML3 cells (1 x 10^6^ cell / sample) were washed twice in ice-cold PBS and collected by centrifugation (3 min at 300 x g). Cell pellets were resuspended in 200 mL of ice-cold PBS and mixed with a solution containing 300 mL of acidified MeOH (0.1 % w/v HCl) and 250 mL hexane and vigorously mixed using a vortex for 2 min. After mixing, the hexane layer was removed by centrifugation (5 min at 17 000 x *g*), dried under a stream of N_2_, resuspended in 200 mL of MeOH containing 2 mM ammonium formate, overlaid with Argon and immediately analysed by LC-MS/MS. Analysis was performed using an I-Class Acquity LC system coupled to a Xevo TQ-S triple quadrupole mass spectrometer (Waters, USA). Sample (2 ⌠L) was injected into a 15 ⌠L sample loop and separated by RP-HPLC at 45 °C using an Acquity C18 column (2.1 x 50 mm, 1.7 ⌠m; Waters). The isocratic mobile phase was 2 mM ammonium formate in MeOH used at 0.8 mL / min over 5 min. The mass spectrometer was operated in positive ion mode with multiple reaction monitoring. The following settings were used for electrospray ionisation in positive ion mode: capillary voltage - 1.7 kV; cone voltage - 30V; ion source temperature 100 °C; collision energy – 22 V. Nitrogen and argon were used as curtain and collision gases respectively. Transitions used for quantification were: UQ_10_, 880.804 > 197.204; UQ_10_H_2,_ 882.904 > 197.204. Samples were quantified using MassLynx 4.1 software to determine the peak area for UQ_10_ and UQ_10_H_2_.

### Reactive Oxygen Species (ROS) Detection by flow cytometry

ROS measurement was performed using the CellROX™ Deep Red Flow Cytometry Assay Kit (Thermo Fisher Scientific, C10491), according to the manufacturer’s instructions. Positive and negative controls were prepared as follows: briefly, a total of 2 x 10^6^ cells (seeded at 0.5 x 10^6^ cells/mL) were incubated in medium supplemented with 200 nM Tert-butyl hydroperoxide (TBHP) for 30 min at 37 °C, as an assay positive control. In a different well, a total of 2 x 10^6^ cells (seeded at 0.5 x 10^6^ cells/mL) were incubated in medium supplemented with 250 μM N-Acetylcysteine (NAC) for 1 hour at 37 °C before treatment with TBHP for 30 min 37 °C, as an assay negative control. Finally, control samples and experimental samples were incubated for 30 min at 37 °C with 550 nM CellROX DR probe, diluted in medium from a freshly made 250 μM probe-stock solution in DMSO (Thermo Fisher Scientific, D12345). During the final 15 min of staining, 1 μM SYTOX™ Blue Dead Cell staining-probe was added to cells (1 nM final concentration). The samples were kept on ice and immediately analysed by flow cytometry (LSR Fortessa cytometer, BD Bioscience). CellROX DR fluorescence was recorded in at least 30,000 live cells (SYTOX blue negative cells) per sample. Flow cytometry data were analysed using Flowjo software. Examples of gating strategy are reported in the Source Data.

### Sinefungin, RSL3 and Ferrostatin-1 Treatment and cell viability assay

Sinefungin (SINE, MedChemExpress, HY-101938-5MG) and Ferrostatin-1 (Fer-1, Selleckchem, S7243) were used at 10 or 50 (SINE) and 5 (Fer-1) μM final concentration, in the cell media. Cells were treated with an equivalent DMSO dilution. For proliferation assays, cells (2 x 10^5^ / mL) either untreated or following doxycycline treatment (100 ng/mL) for four days were seeded in DMSO or 5 μM Fer-1 and cultured for a further 4 days. Fresh media for all treatments was replaced every 2 days. For sinefungin treatment, cells (2 x 10^5^ / mL) were cultured in the presence of 50 μM sinefungin for four days. Fresh medium containing the respective treatments was replaced every two days. Cell proliferation was measured by CellTiter-Blue® Cell Viability Assay kit (Promega, G8081), according to the manufacturer’s instructions. Proliferation was analysed as the ratio between the initial number of seeded cells and the number of cells after incubation and normalised to DMSO control-treated cells representing 100% of the proliferating cells.

To detect RSL3-sensitivity, OCI-AML3 cells were cultured in doxycycline-containing medium (100 ng/mL) and decreasing concentrations of RSL3 (Selleckchem, S8155): 300 nM, 200 nM, 100 nM, 50 nM for the EC50 calculation; 40 nM, 30 nM, 20 nM and 15 nM for sensitisation experiments. Following 5 days of doxycycline treatment, cells (2 x 10^5^ / mL) were seeded in a 96 multi-well plate and incubated for 24 h with RSL3 at the indicated doses, DMSO or RSL3 in combination with 5 μM Fer-1. Relative Fluorescence units (RFU) for each well were normalized to the median RFU from the DMSO control wells taken as 100% cell viability. CellTiter-Blue® Fluorescence analysis was performed using the SpectraMax iD3 plate reader (Molecular Devices). Three technical replicates per drug condition were performed as well as three independent biological replicates. IC50 values of RSL3 (half maximal inhibitory concentration causing 50% inhibition of cell proliferation) was calculated using GraphPad Prism v11 (RRID:SCR_002798), and the curves show the mean ± SD of three replicates per condition measured.

### Lipid Peroxidation Assay by flow cytometry

The lipid peroxidation assay was performed using the Lipid Peroxidation Sensor BODIPY™ 581/591 C11 (Thermo Fisher Scientific, D3861), according to manufacturer’s instructions. Positive and negative controls were prepared as follows: briefly, a total of 2 x 10^6^ cells (seeded at 0.5 x 10^6^ cells/mL) were incubated in media supplemented with 0.5 μM RSL3 for 2 h at 37 °C, as an assay positive control; in a different well, a total of 2 x 10^6^ cells (seeded at 0.5 x 10^6^ cells/mL) were incubated in media supplemented with 0.5 μM RSL3 and 5 μM FeR-1 for 2 h at 37 °C, as an assay negative control. DMSO incubation for 2 h at 37 °C was also performed, as a vehicle condition. Alternatively, following 5 days of doxycycline treatment, OCI-AML3 cells were treated for 24 hours with 300 nM RSL3 or DMSO. Finally, control samples and experimental samples were incubated (10^6^ cells/mL) for 30 min at 37 °C with 2.5 μM BODIPY™ 581/591 C11 staining solution, freshly made in media on the day of the assay. After staining, cells were washed three times with PBS, kept on ice and immediately analysed by flow cytometry (LSR Fortessa cytometer and BD FACSCelesta, BD Bioscience). Reduced and oxidised BODIPY™ 581/591 C11 fluorescence was recorded for at least 15,000 events per sample. Flow cytometry data were analysed using Flowjo software or BD FACSCelesta analysis software. Examples of gating strategy are reported in the Source Data.

### Generation of HEK-293T cells stably expressing APEX2 constructs

ShCTRL and sh1TGS1 HEK-293T cells were infected at ∼50% confluency with lentiviral vectors expressing APEX2-OMM and APEX2-NES (CYTO, cytoplasmic form) constructs^42^, followed by selection in 8 mg/mL blasticidin supplemented growth medium for 7 days before further analysis. APEX2 construct expression was verified by immunoblotting, as described above, and their correct localisation at the Outer Mitochondrial Membrane (OMM) or in the cytoplasm (CYTO) was verified by immunofluorescence, as described in the “Immunofluorescence staining and confocal imaging” section of the Methods (below).

### APEX labelling in living cells

Apex labelling in shCTRL and sh1TGS1 HEK-293T cells expressing APEX2-OMM and APEX2-NES was performed as previously described^42^, with small changes. Briefly, 24 h after seeding HEK-293T cells (corresponding to day 6 of doxycycline treatment), APEX labelling was started by adding fresh medium containing 1 mM biotin-tyramide (Iris Biotech GMBH, LS-3500) and incubated for 30 min at 37°C at 5% CO_2_. For the cells treated with cycloheximide (CHX), 15 min after biotin-incubation, CHX was added to the medium to a final concentration of 0.1 mg/mL and the cells were further incubated for another 15 min at 37°C at 5% CO_2_. For the cells treated with puromycin (PUR), APEX labelling was started by adding a fresh medium containing 1 mM biotin-tyramide and 200 μM puromycin and incubated for 30 min at 37°C at 5% CO_2_. After incubation, cells were washed three times with wash buffer (0.5 mM MgCl_2_, 1 mM CaCl_2_ in DPBS), then 0.5 mM H_2_O_2_ (Fisher Scientific, HYD005) in wash buffer was added and the plates were gently agitated for 1 min. The reaction was immediately quenched by replacing the H_2_O_2_ solution with an equal volume of 5 mM Trolox (Sigma-Aldrich, 238813-5G), 10 mM sodium ascorbate (Sigma-aldrich, A7631-25G) and 10 mM sodium azide in DPBS (Thermo Fisher Scientific, D8537). Cells were finally washed with DPBS containing 5 mM Trolox and 10 mM sodium ascorbate three times before proceeding to RNA extraction. The unlabelled controls were processed identically, except that the H_2_O_2_ addition step was omitted. Total RNA was extracted using the RNeasy mini kit (Qiagen, 74104), following the manufacturer’s protocol, including adding β-mercaptoethanol to the lysis buffer, and replacing the RW1 buffer with RWT buffer (Qiagen, 1067933) for washing.

### Enrichment of biotinylated RNA

To enrich the APEX-biotinylated RNA we used the protocol described in42, with small changes. Pierce streptavidin magnetic beads (Thermo Fisher Scientific, 88816) were washed 3x in B&W buffer (5 mM Tris-HCl, pH = 7.5, 0.5 mM EDTA, 1 M NaCl, 0.1% TWEEN 20), followed by 2x in Solution A (0.1 M NaOH and 0.05 M NaCl), and 1 time in Solution B (0.1 M NaCl). The beads were then suspended in 150 μL 0.1 M NaCl and incubated with 25 μg of RNA, diluted in water and RNaseOUT to a final volume of 150 μL, at RT for 2 h on a shaker (500 rpm). The beads were then placed on a magnet, the supernatant discarded, and three washes in B&W buffer were performed before resuspending the beads in 56 μl water. A 3X proteinase digestion buffer was prepared as follows: 1.1 mL buffer contained 330 μl 10X PBS pH = 7.4, 330 uL 20% N-Lauryl sarcosine sodium solution (Sigma Aldrich, L7414-10ML), 66 μL 0.5M EDTA, 16.5 μL 1M dithiothreitol (DTT, Thermo Fisher Scientific, 707265ML) and 357.5 μL water. 33 uL of this 3X digestion buffer was added to the beads along with 10 μl Proteinase K (20 mg/mL, Ambion, AM2548) and 1 μL RNaseOUT. The beads were then incubated at 42°C for 1 h, followed by 55°C for 1 h on a shaker (1100 rpm). The RNA was then purified using the RNA clean and concentrator −5 kit (Zymo Research). A fixed volume of RNA was retro-transcribed and relative RNA abundance was measured by RT-qPCR as described above. For each mRNA the enrichment of RNA recovered in the labelled samples over RNA recovered in the un-labelled sample (ΔCt = CtLABELLED-CtUNLABELLED) was calculated for both APEX2 construct, OMM and CYTO. For each mRNA, the ratio between the OMM enrichment over the CYTO enrichment (ratio enrichment = ΔCtOMM/ΔCtCYTO) in both treatment conditions, CHX and PUR, was calculated. A complete list of primers can be found in Supplementary Data 7b.

### Immunofluorescence staining and confocal imaging

HEK-293T were seeded on sterile coverslips, pre-treated with poly-D-Lysine (Thermo Fisher Scientific, A3890401). For eIF3b imaging, cells were treated with 0.1 mg/mL of cycloheximide for 15 min. Cells were then fixed in 5% paraformaldehyde (PFA) in PBS (pH 7.4) for 15 min at 37°C, with three subsequent washes in PBS. Autofluorescence was quenched with 50 mM ammonium chloride for 10 min at room temperature (RT) and three additional washes in PBS. Cells were then permeabilized with Triton X-100/0.1% PBS for 10 min, followed by three washes in PBS. Blocking was carried out in 10% FBS/PBS for 20 min. Primary antibodies were prepared in 5% FBS/PBS and incubated on a shaker for 2 hours overnight at 4°C (Supplementary Data 7c). Cells were washed three times in 5% FBS/PBS and incubated with the corresponding secondary antibody (dilution 1: 1000, Supplementary Data 7c) prepared in 5% FBS/PBS. Coverslips were washed three times in PBS and then in deionized water (DIW), dried, and mounted onto slides using fluorescence mounting medium (Dako).

Images were acquired on a Nikon Eclipse TiE inverted microscope coupled with an Andor Dragonfly 500 confocal spinning disk system using a 100× objective. Images were captured with 15 stacks each of 0.2 μm using the Zyla 4.2 PLUS sCMOS camera (Andor), which was coupled to Fusion software (Andor). Images were acquired under the same conditions with the same laser intensity (laser lines 488, 568, and 647 nm) and exposure time. Raw seven-stack images were compiled into “max projection,” converted into composites, and processed using the Fiji software.

### Formaldehyde-cross linked RNA Immunoprecipitation

Immunoprecipitation of endogenous eIF3 complexes was performed in OCI-AML3 cells, according to a protocol adapted from^43^. Cells were washed with PBS and the pellets were resuspended in 1x volume of 0.2% formaldehyde for 5 min on a rocker to stabilize RNP complexes. Cross-linking reactions were quenched by the addition of 0.15 M glycine, pH 7 for 5 min on a rocker. Cells were subsequently lysed in RNP buffer (10 mM HEPES-NaOH pH 7.9, 100 mM KCl, 5 mM MgCl2, 0.5% NP-40, 1 mM DDT, complete protease inhibitor cocktail). Next, 1 mg of total protein extract was incubated under rotation for 2h at 4° with 2 μg of either an antibody against eIF3b or an anti-GFP control antibody (Supplementary Data 7c), and with 100 μL of Dynabeads Protein G (Invitrogen, 10004D) in a total 1 mL of IP Buffer (50 mM Tris-HCl pH 8, 150 mM NaCl, 1% Triton X-100). Of the total input, 10% was saved before immuno-precipitation. After incubation, complexes were washed 3 times in wash buffer (10mM Tris-HCl pH 7.4, 1mM EDTA pH 8, 1mM EGTA, 150mM NaCl, 1% Triton) and then RNA was eluted at 56° on a shaker (1100 rpm) for 30 minutes in 200 μL of Elution Buffer (0.1 M NaHCO_3_, 1%SDS and 1 uL proteinase-K 20mg/mL per sample). Eluted RNA was purified using the RNA Clean & Concentrator-5 Kit (Zymo Research, R1014). A fixed volume of RNA was retro-transcribed as described above and CLIP-enriched RNA was quantified by RT–qPCR. The % of RNAs in the IP was determined by the ΔCq method as enrichment over anti-GFP controls. A complete list of primers can be found in Supplementary Data 7b.

### Proximity Ligation Assay (PLA)

To detect protein-protein interactions, a Proximity Ligation Assay (PLA) was performed using the Duolink in Situ PLA kit (Sigma-Aldrich), according to the manufacturer’s instructions. Briefly, cells were seeded on sterile glass coverslips in 24-well plates and allowed to adhere overnight. Cells were then fixed with 4% PFA in PBS for 15 minutes at room temperature, followed by permeabilization with 0.1% Triton X-100 in PBS for 10 minutes. After washing with PBS, cells were blocked with Duolink Blocking Solution for 1 hour at 37°C. After blocking, cells were incubated with primary antibodies against the proteins of interest (e.g., eIF3b, eIF3e, eIF3d and TGS1) diluted in Duolink Antibody Diluent. The primary antibodies were used at optimized concentrations (e.g., 1:200). Incubation was performed overnight at 4°C. Following primary antibody incubation, cells were washed with PBS and incubated with species-specific PLA probes (anti-rabbit PLUS and anti-mouse MINUS) diluted in Duolink Antibody Diluent for 1 hour at 37°C. Cells were washed and then incubated with ligation solution containing ligase for 30 minutes at 37°C to allow for the formation of circular DNA strands. This was followed by incubation with the amplification solution containing polymerase for 100 minutes at 37°C to amplify the DNA circles, which were detected as fluorescent dots. After amplification, coverslips were washed, mounted with Duolink Mounting Medium containing DAPI, and sealed with nail polish. Fluorescent signals were visualized and captured using a Leica TCS SP8 confocal microscope (Leica Microsystems). PLA signal quantification was performed with ImageJ software on a total of 9, 5 and 6 images for eIF3b, eIF3e and eIF3d, respectively, obtained from at least three independent experiments. A complete list of antibodies can be found in Supplementary Data 7c.

### Exogenous TGS1 expression in OCI-AML3 cells

A codon-optimized, shRNA-resistant TGS1 open reading frame with a C-terminal HA tag was synthesized (ThermoFisher Scientific; Supplementary Data 7b) and cloned into the pHIV-ZsGreen lentiviral vector (Addgene plasmid #18121). Lentiviral transduction of control and TGS1-knockdown OCI-AML3 cells was carried out as described above. Cells transduced with the pHIV-ZsGreen constructs were subsequently isolated by flow-cytometric sorting based on GFP fluorescence.

### Statistical analysis

All data in bar- and line-plots are presented as means ± SD of at least three independent replicates, unless stated otherwise. Significance was determined using nested a one-way analysis of variance (ANOVA), one-way ANOVA, two-way ANOVA, unpaired *t* test or a Wilcoxon test (when distributions were assessed not to be normal and homoscedastic) at a confidence interval of 95%, unless otherwise specified. No statistical methods were used to predetermine sample sizes. All statistical analyses were performed using GraphPad Prism v11 software. \**P* < 0.05, \*\**P* < 0.01, \*\*\**P* < 0.001, and \*\*\*\**P* < 0.0001 were considered significant, and *P* > 0.05 was considered non-significant (ns).

## Data availability

Sequencing data that support the findings of this study have been deposited in the NCBI Sequence Read Archive under BioProject ID PRJNA1229521. Data are temporarily available for reviewers at the following link: https://dataview.ncbi.nlm.nih.gov/object/PRJNA1229521?reviewer=sugar47u6liddlmreh9oe2b9ot

Mass Spectrometry proteomic raw data are available via ProteomeXchange with identifier PXD073282.

Computational codes for K-mer signatures of RNA populations and individual transcripts, with RBP association analysis are available at the following link: https://github.com/AngeloDAngelo/Ktools

Source Data are provided with this manuscript.

## Acknowledgements

This research was supported by Cancer Research UK (grant reference RG86786) (V.M, A.M.B., S.L., IB), by the Joseph Mitchell Fund (I.B.) and by AIRC (AIRC Start-Up 26505) (M.B., D.I., I.B.) and (AIRC Star-UP 22895) (L.B.). Work in Micheal P.M.’s lab was supported by the Medical Research Council UK (MC_UU_00028/4) and by a Wellcome Trust Investigator award (220257/Z/20/Z). J.Lj.M. was additionally supported by the Biotechnology and Biological Sciences Research Council (BLAST Pumping Award, Grant No: BB/W01825X/1). S.D.T was supported by the project National Institute for Cancer Research (Programme EXCELES, ID Project No. LX22NPO5102) - Funded by the European Union - Next Generation EU. The research was further supported by Intramural IIT funding (S.G, L.P.). D.V. was supported by an Italian Cancer Research Foundation (FIRC) post-doctoral fellowship (Project No. 31186). G.C. was supported by Giovani Ricercatori-GR-2021-12374957 and is currently supported by AIRC under MFAG 2025 (ID: 31911). Valeria P. was supported by AIRC (IG#24851). P.E.P. was supported by AIRC (IG 2025 #30515). A.M.S. was supported by Individual AIRC START-UP grant (#29211).

This research was supported by the Cambridge NIHR BRC Cell Phenotyping Hub (Department of Medicine, University of Cambridge, UK). We wish to thank the staff members for their advice and support in Flow Cytometry. We thank the RNA Initiative@IIT, Dr. Andy J Bannister and Dr. Gonzalo Millan for thought-provoking discussions.

## Authors Contributions

I.B. and V.M. devised, designed and supervised the project. V.M. and G.C. performed most of the experimental studies, with the fundamental assistance of M.B., J.L.M., Vincent P., C.S.Y., D.I., S.M., L.M., D.V., A.M.B. and S.L. for specific techniques. A.D. and A.S. performed computational analysis of RNA-motifs discovery. L.P. performed computational analysis of data from sequencing and proteomic experiments, and related additional analysis. Maria P.M, L.B. and A.M.S. provided AML patients’ samples and supervised the related experiments. D.A., Valeria P., P.E.P, S.G., A.M., S.D.T, Micheal P.M. supervised specific experimental studies. I.B. and V.M wrote the manuscript with significant contribution from all the authors.

## Competing Interests

The authors declare no competing interests

## Notes

### Competing Interest Statement

The authors have declared no competing interest.

### Summary of Updates

In this new version, we further dissected the molecular mechanism of TGS1 function in leukaemia cells. To clarify the specificity of TGS1 targets, we applied a newly developed bioinformatic approach that identified a set of RNA sequence k-mers strongly enriched in the 5 UTRs of TGS1 mRNA targets. Additionally, we investigated the interaction between TGS1 and the eIF3 translation initiation complex. We identified a specific interaction between TGS1 and eIF3d and showed that its depletion strongly decreases m2,2,7G methylation on TGS1 targets, providing a mechanistic explanation of TGS1 selectivity for mRNAs encoding mitochondrial proteins. We tested the effect of sinefungin, a known inhibitor of TGS1, on primary leukaemia blasts derived from freshly diagnosed patients and primary CD34+ healthy blasts. Our data support a specific effect of the inhibitor on the colony-forming potential and viability of leukaemia blasts.

