## Supplementary Figures for "TGS1 mediates mRNA 5’-cap trimethylation to promote oxidative phosphorylation in acute myeloid leukaemia"

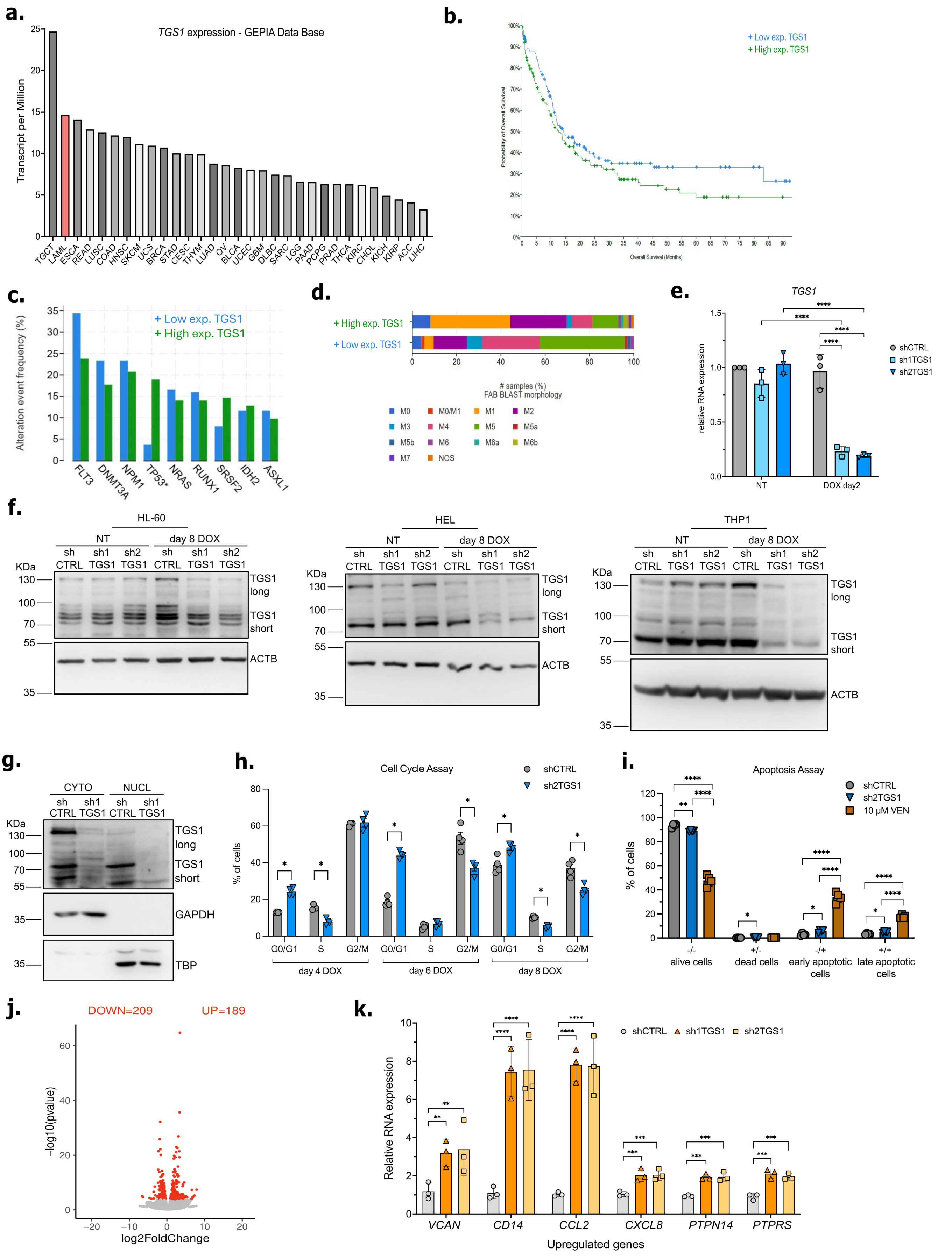

**Supplementary Figure 1. Generation of inducible TGS1-knock-down cell lines and RNA-seq analysis**

**a.** Average *TGS1* expression levels across cancer types from the TCGA public dataset, reported as transcripts per million (TPM). Expression in the Acute Myeloid Leukemia (LAML) dataset is highlighted in red. **b.** Kaplan–Meier survival curves of AML patients with low versus high *TGS1* expression, derived from the combined BEAT AML and TARGET AML datasets on cBioPortal (https://www.cbioportal.org/). **c.** Frequency of genomic alterations in the most commonly mutated genes in the same cohort of **b.**, comparing patients with low versus high *TGS1* expression. **d.** Distribution of FAB blast morphologies in patients with low versus high *TGS1* expression in the same cohort of **b. e.** RT-qPCR measure of *TGS1* mRNA expression in untreated (NT), shCTRL, sh1TGS1 and sh2TGS1OCI-AML3 cells or following 2 days of DOX treatment **f.** Immunoblot of TGS1 protein level in the indicated AML cell lines untreated (NT) and following 8 days of DOX treatment. A representative immunoblot of three independent experiments is shown. **g.** Immunoblot of cytoplasmic (CYTO) and nuclear (NUCL) localisation of the long and short isoforms of TGS1 in shCTRL and shTGS1 OCI-AML3 cells following 4 days of DOX treatment. GAPDH or TBP were used as loading control of CYTO and NUCL fractions, respectively. A representative immunoblot of two independent experiments is shown **h.** Flow cytometric assessment of cell-cycle distribution by propidium iodide (PI) staining in shCTRL and sh2TGS1 OCI-AML3 cells following the indicated DOX treatment. **i.** Flow cytometric detection of PI/Annexin V stained shCTRL or sh2TGS1 OCI-AML3 cells following 8 days of DOX treatment. **j.** Volcano plot of the log2 fold change of differentially expressed transcripts following TGS1 knockdown in OCI-AML3 cells. Downregulated (left) and Upregulated (right) genes are highlighted in red. **k.** RT-qPCR measure of upregulated genes in **j.** Values in **e and j** are mean ± SD of $n = 3$ independent experiments. Values in **h and i** are mean ± SD of $n = 4$ independent experiments. $^{**}P < 0.01$, $^{***}P < 0.001$, $^{****}P < 0.0001$, two-way ANOVA test with Tukey's (**e.**) or Dunnet's (**j.**) multiple comparisons, or multiple unpaired t-tests in **h.** and **i.**

**a.**

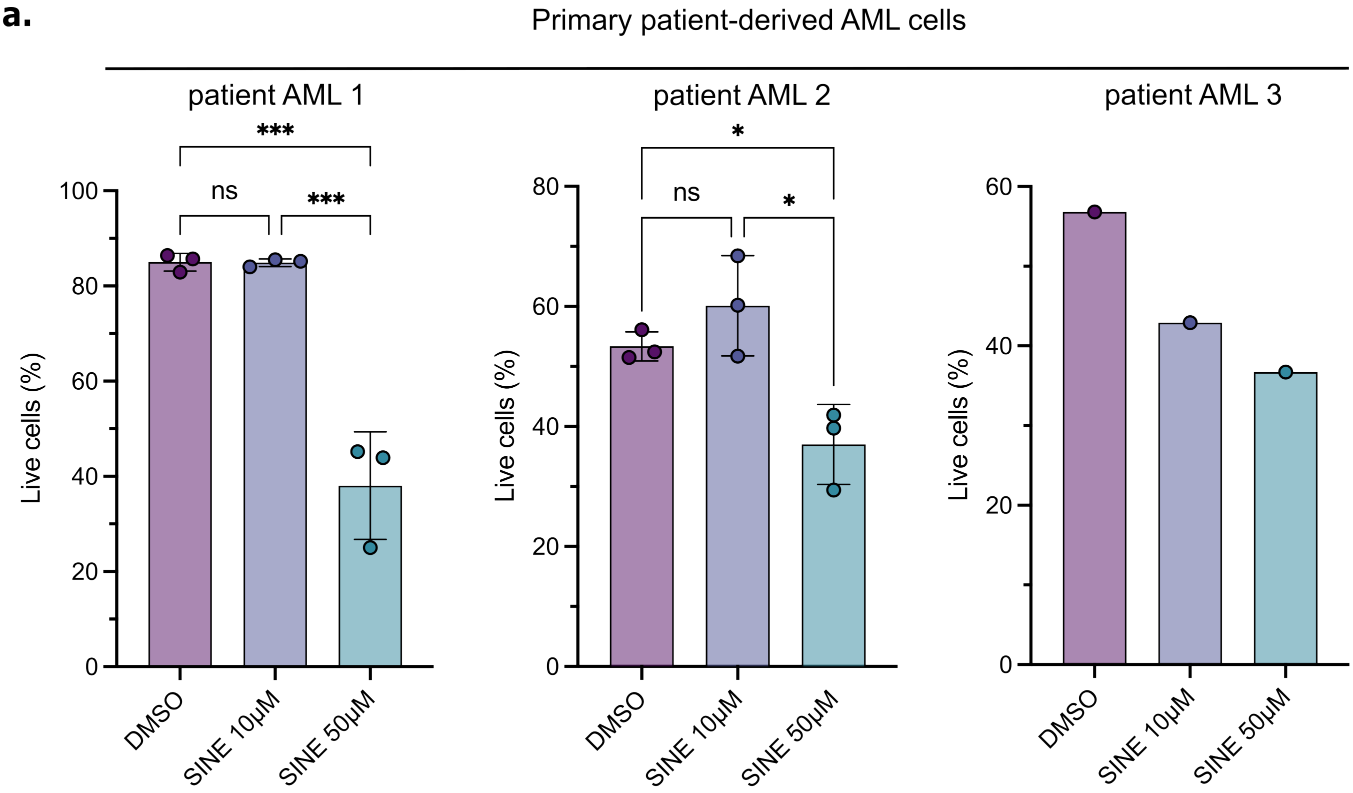


**Supplementary Figure 2. Inhibition of TGS1 catalytic activity by sinefungin reduces viability of primary blast derived from AML patients**

**a.** Percentage of live cells derived from AML patients' blasts were quantified after 7 (AML 1) or 14 (AML2 and AML3) days of culture following treatment with SINE (10 or 50 μM) or DMSO treatment. Values represent means ± SD of *n* = 3 independent experiments. *P<0.05 and ***P<0.001, one-way ANOVA test with Tukey's multiple comparisons.

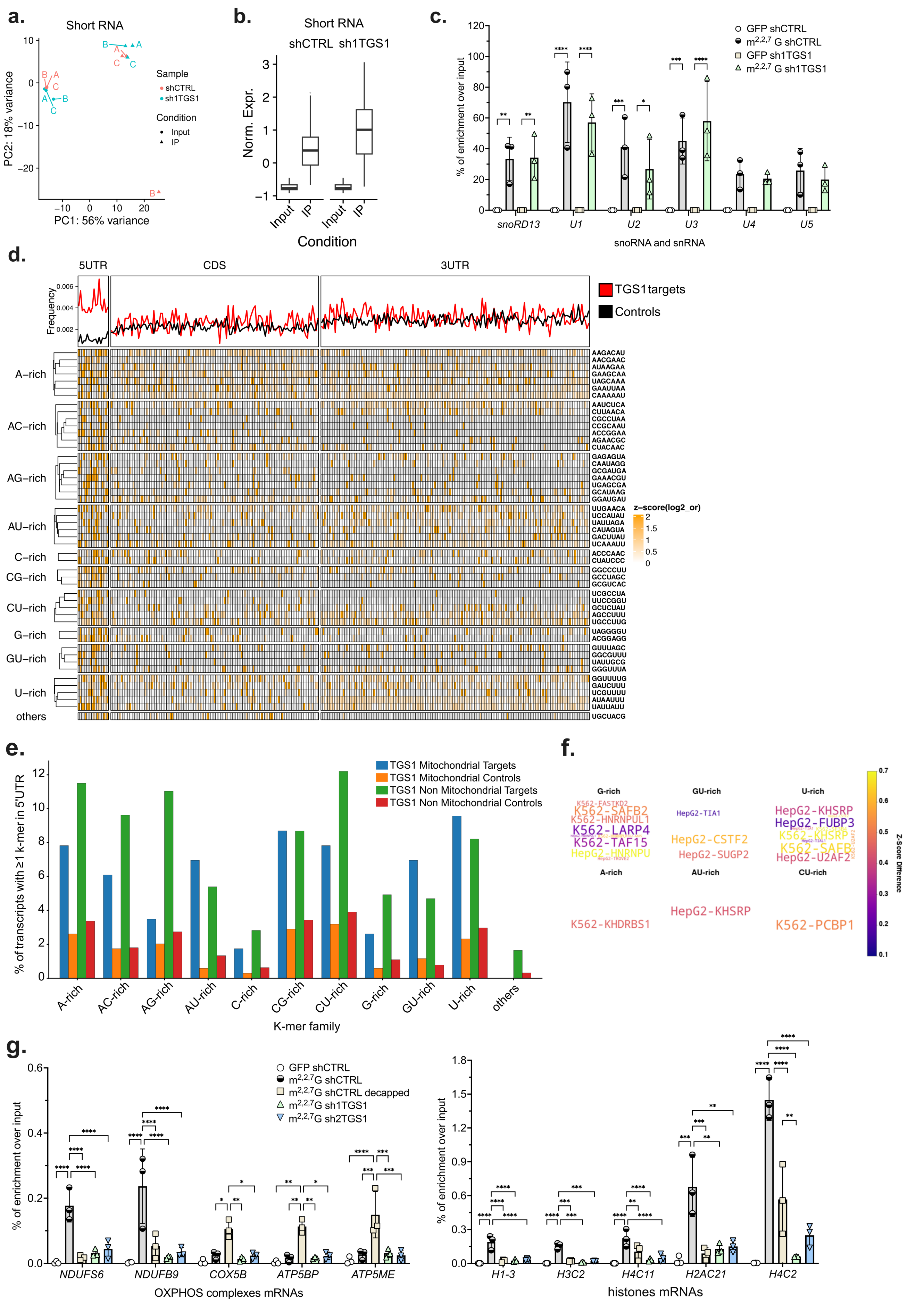


**Supplementary Figure 3. Validation of the $m^{2,2,7}$G-RIP-seq data by RT-qPCR and RNA motif analysis**

**a.** PCA analysis of the short RNAs (<200bp) $m^{2,2,7}G$ RNA-IP-Seq of both input (circles) and immunoprecipitation (triangles) datasets of 3 biological replicates of shCTRL and sh1TGS1 OCI-AML3 cells following 6 days of DOX treatment. **b.** Normalized enrichment of $m^{2,2,7}G$-modified short RNAs in shCTRL (left) and sh1TGS1 (right) OCI-AML3 cells after 6 days of DOX treatment. **c.** $m^{2,2,7}G$-RIP-RT-qPCR validation on short RNA purified from shCTRL or sh1TGS1 OCI-AML3 cells after 6 days of DOX treatment. An antibody anti-GFP was used as a technical negative control (white circles). **d.** *Top panel*: line plot showing the average frequency of enriched 7-mers along 5′UTR, CDS and 3′UTR regions in TGS1 targets (red) versus length- and mitochondrial-representation–matched controls (black) meta-transcript. *Bottom panel:* heatmap of z-scores of $\log_2$ odds ratio (color scale) for the 49 significantly enriched 7-mers in TGS1 targets group versus controls (rows, hierarchically clustered and grouped by nucleotide composition: A-rich, AC-rich, AG-rich, AU-rich, C-rich, CG-rich, CU-rich, G-rich, GU-rich, U-rich, and others) across individual TGS1-target transcripts (columns), separated by transcript region (5′UTR, CDS, 3′UTR). **e.** Bar plot showing, for each 5′UTR k-mer family (x-axis), the percentage of transcripts containing at least one occurrence of that family, separately for mitochondrial targets (blue), mitochondrial controls (orange), non-mitochondrial targets (green) and non-mitochondrial controls (red). **f.** K-mer RBP binding propensity (K-RBP) analysis of the 5′UTR-enriched 7-mers, grouped by nucleotide-composition family (G-rich, GU-rich, U-rich, A-rich, AU-rich, CU-rich). Word size represents the difference in eCLIP-derived z-scores between the selected k-mers and the reference set, with larger words indicating stronger predicted RBP association. Only RBPs with a z-score difference ≥ 0.47 and Bonferroni-adjusted p ≤ 0.05 are shown; the top 10 RBPs ranked by z-score difference are displayed for each k-mer family. **g.** $m^{2,2,7}G$-RIP-RT-qPCR validation on long RNA purified from shCTRL or sh1TGS1 OCI-AML3 cells after 6 days of DOX treatment. An antibody against GFP was used as a technical negative control (white circles). Values in **c** and **g** are means ± SD of *n* = 3 independent experiments. *P<0.05, **P<0.01, ***P<0.001, ****P<0.0001, two-way ANOVA test with Tukey's multiple comparisons.

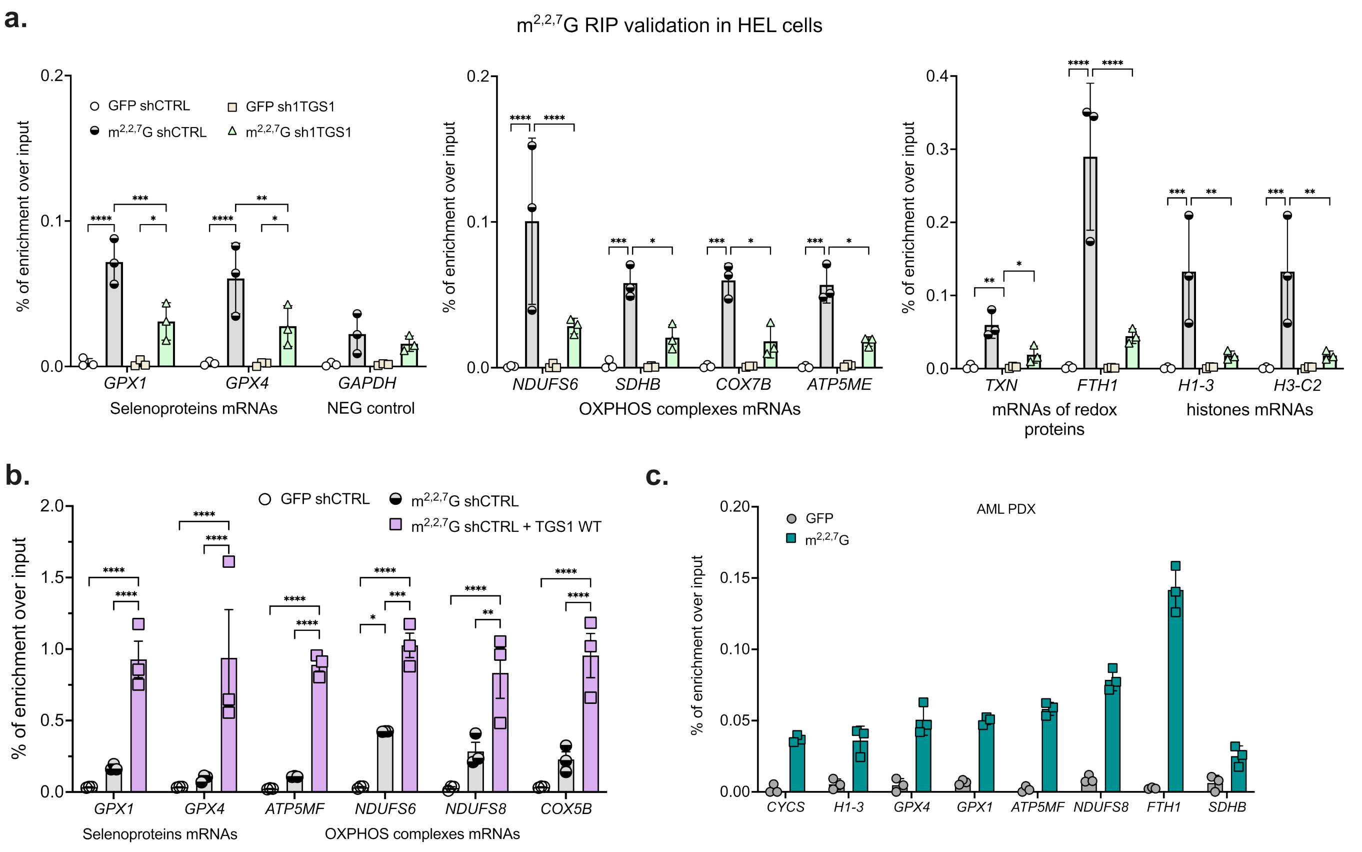


**Supplementary Figure 4. Validation of the m$^{2,2,7}$G-RIP by RT-qPCR in additional cell lines**

**a.** m$^{2,2,7}$G-RIP-RT-qPCR validation on long RNA purified from shCTRL or sh1TGS1 HEL cells after 6 days of DOX treatment. An antibody against GFP was used as a negative control (white circles). **b.** m$^{2,2,7}$G-RIP-RT-qPCR was performed on long RNA purified from OCI-AML3 shCTRL cells and OCI-AML3 shCTRL cells overexpressing TGS1. An antibody against GFP was used as a technical negative control. **c.** m$^{2,2,7}$G-RIP-RT-qPCR was performed on long RNA purified from AML cells derived from an *NPM1c/FLT3ITD/ITD* primary derived xenograft model (PDX2)[31]. An antibody against GFP was used as a technical negative control (triangles). Values in **c** are means ± SD of three independent technical replicates. Values in **a** and **b** are means ± SD of $n = 3$ independent experiments. *P<0.05, **P<0.01, ***P<0.001, ****P<0.0001, two-way ANOVA test with Tukey's multiple comparisons.

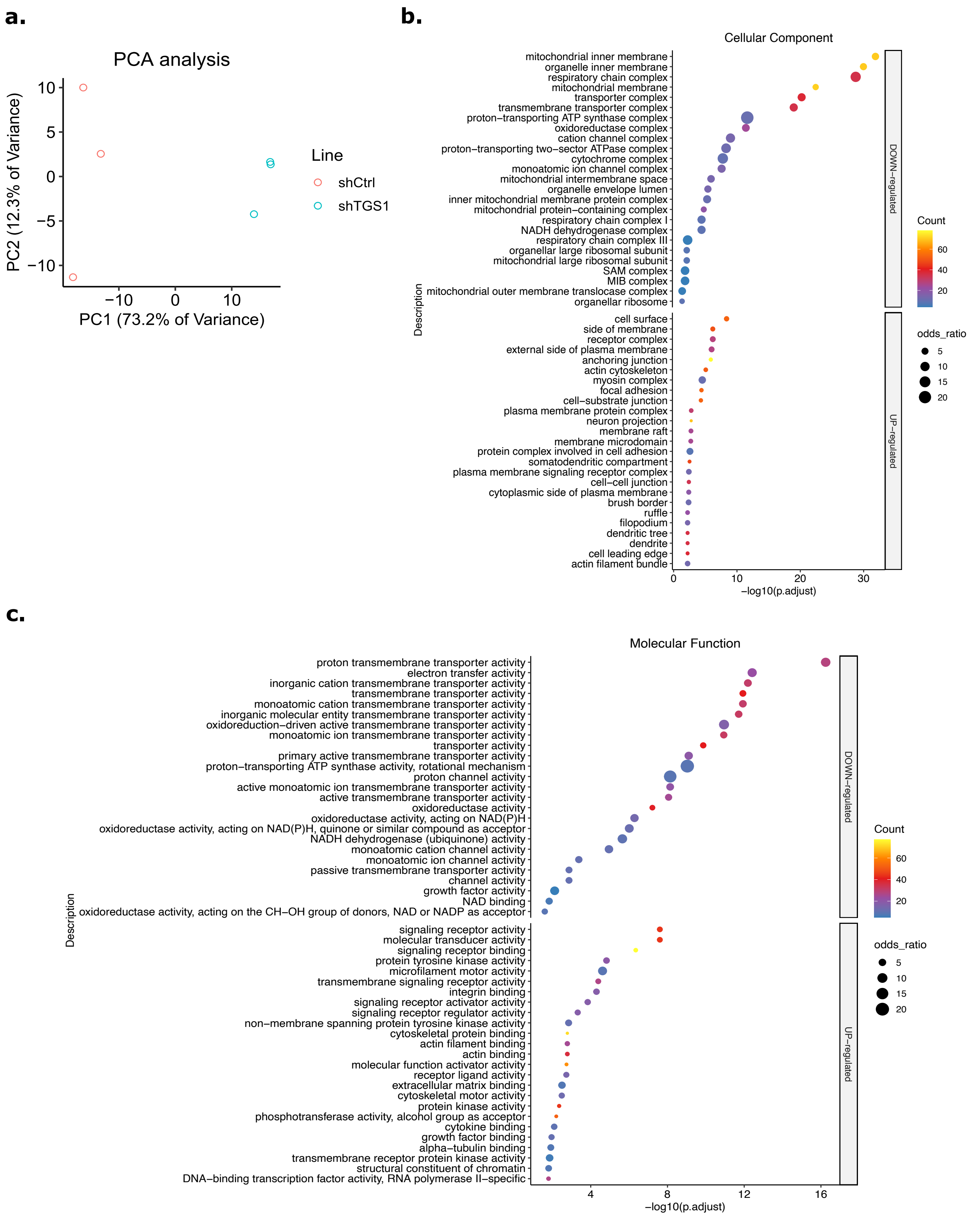


**Supplementary Figure 5. Loss of TGS1 selectively lowers protein expression from m$^{2,2,7}$G-capped transcripts**

**a.** PCA analysis of the proteomic datasets from shCTRL and sh1TGS1 OCI-AML3 cells following 6 days of doxycycline treatment, with three independent replicates for each condition. **b.** Gene

Ontology (GO) analysis of Cellular Components associated with downregulated or upregulated proteins in sh1TGS1 compared with shCTRL OCI-AML3 cells. **c.** Gene Ontology (GO) analysis of Molecular Functions associated with downregulated or upregulated proteins in sh1TGS1 compared with shCTRL OCI-AML3 cells.

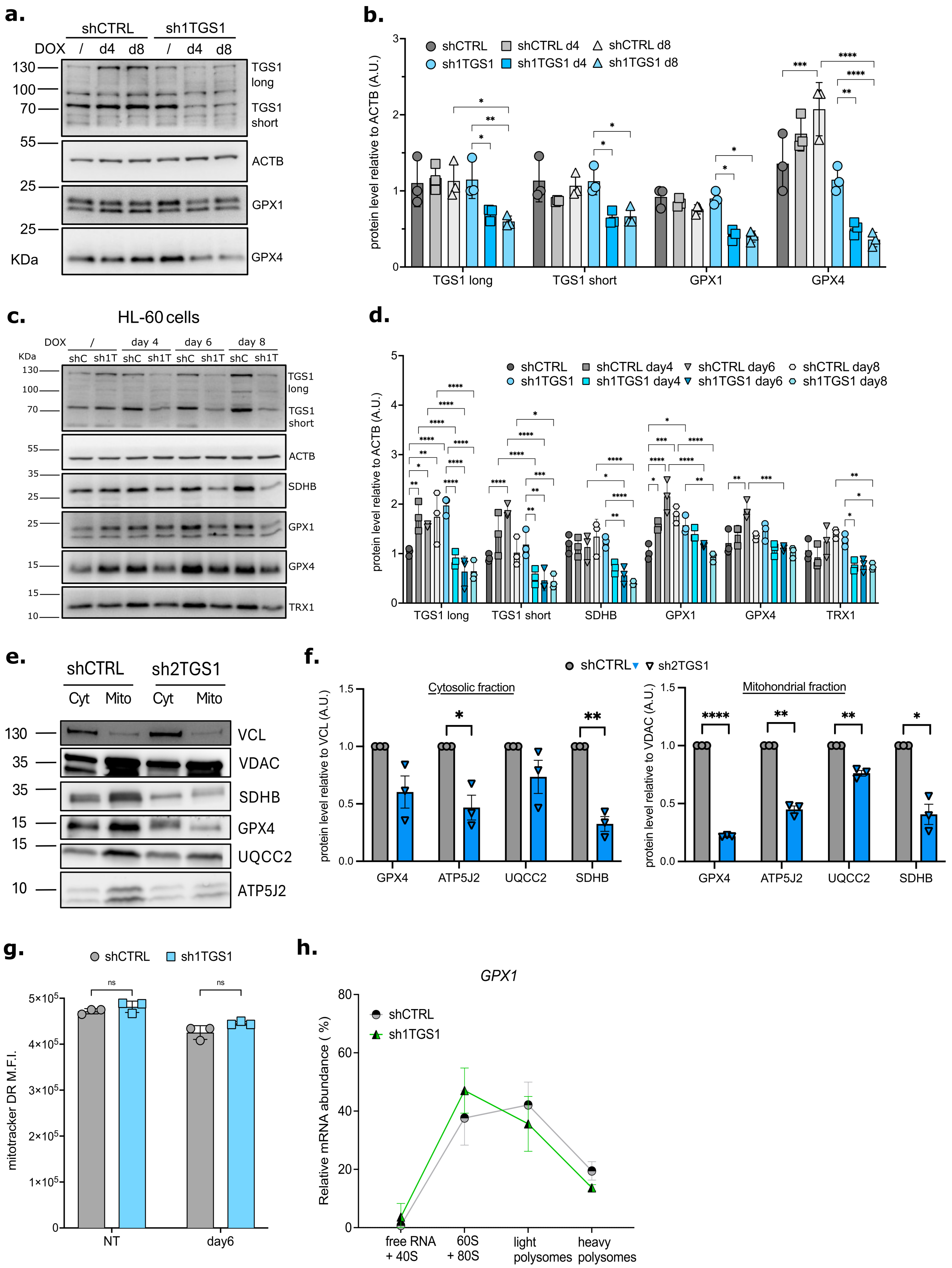


**Supplementary Figure 6. TGS1 silencing affects translation efficiency of modified targets**

**a.** Representative immunoblot of total protein extracts of shCTRL or sh1TGS1 OCI-AML3 cells at the indicated days of DOX treatment with the indicated antibodies. **b.** Densitometric analysis of the immunoblotting shown in panel **a**, normalised to ACTB. **c.** Representative immunoblot of total protein extracts from shCTRL and sh1TGS1 HL-60 cells at the indicated times after DOX treatment with the indicated antibodies. **d.** Densitometric analysis of the immunoblotting shown in panel **c**, normalised to ACTB. **e.** Immunoblot of cytoplasmic (Cyt) and mitochondrial (Mito) fractions purified from shCTRL and sh2TGS1 OCI-AML3 cells after 6 days of doxycycline treatment, using the indicated antibodies. A representative blot of three independent experiments is shown. **f.** Densitometric analysis of the immunoblotting shown in panel **e.** Each fraction was normalized using an internal control (upper panel: VCL for the Cyt fraction; lower panel: VDAC for the Mito fraction). **g.** Mean Fluorescence Intensity (M.F.I.) of MitoTracker™ DR probe measured by flow cytometry in shCTRL and sh1TGS1 OCI-AML3 cells, untreated (NT) or following 6 days of DOX treatment. **h.** *Gpx1* mRNA in each pooled polysome fraction was quantified by RT-qPCR and plotted as a percentage of the total. Values in **b**, **d**, **e**, **f** and **g** are mean ± SD of $n = 3$ independent experiments. Values in **h** are mean ± SD of $n = 3$ independent experiments. *P<0.05, **P<0.01, ***P<0.001, ****P<0.0001 and ns= non-significant two-way ANOVA test with Tukey's or Šidák's (**g**) multiple comparisons, or multiple paired t-test (**e**)

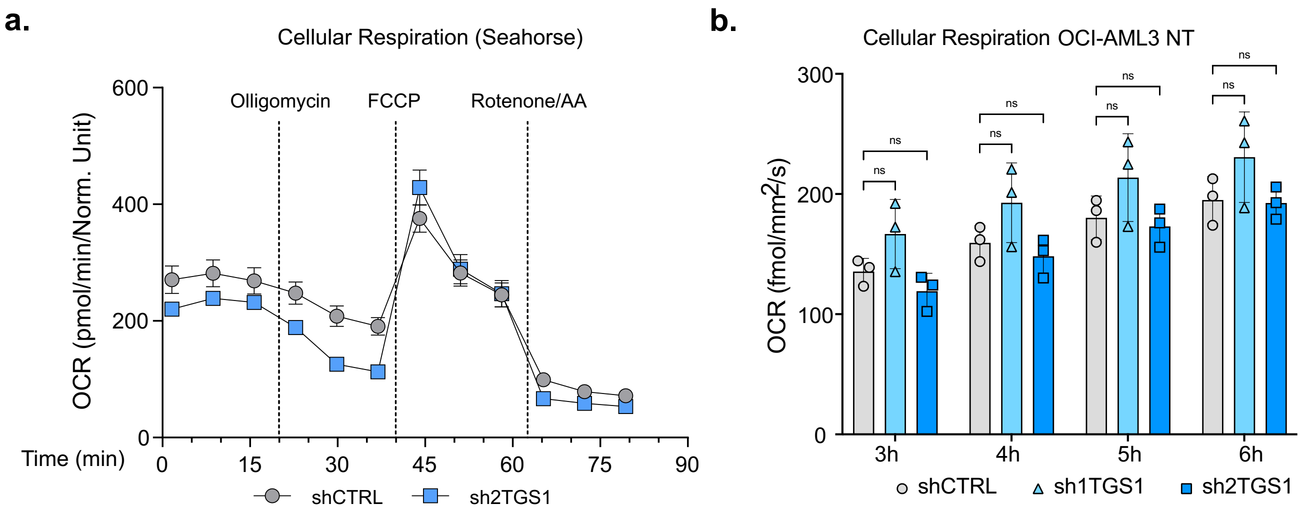


**Supplementary Figure 7. OCR of sh2TGS1 and untreated OCI-AML3 cells**

**a.** OCR profile of shCTRL and sh2TGS1 OCI-AML3 cells determined upon injection of oligomycin, FCCP, rotenone and antimycin A in specific Seahorse medium. Data (mean ± SEM) are normalized by cell count. **b.** Basal respiration of untreated (NT) shCTRL, sh1TGS1 or sh2TGS1 OCI-AML3 cells was measured using Resipher instrument and recorded at the indicated times (hours). Values are mean ± SD of *n*= 3 independent experiments. ns= non-significant, two-way ANOVA test with Tukey's multiple comparisons.

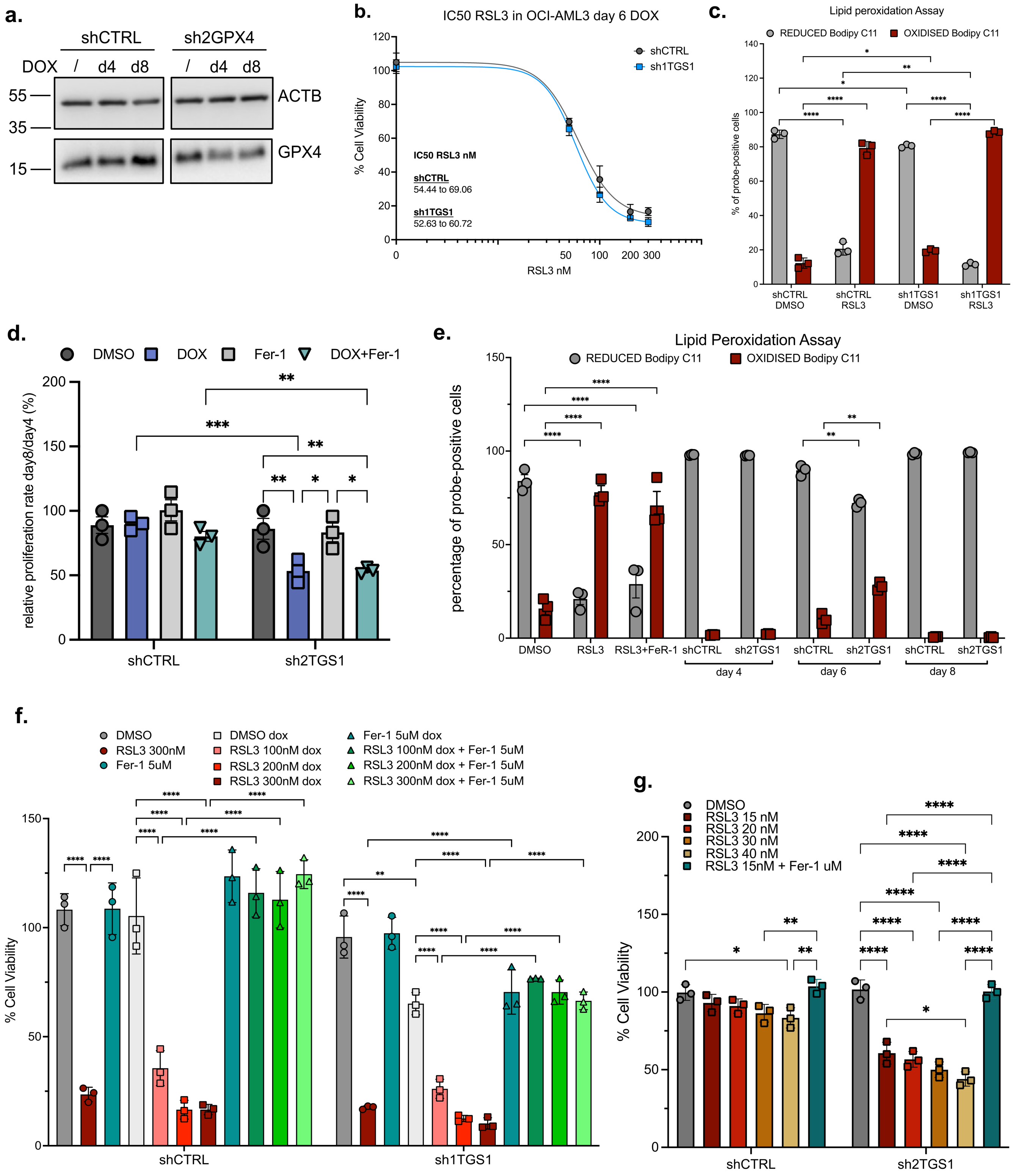


**Supplementary Figure 8. RSL3 sensitivity of shCTRL and shTGS1 OCI-AML3 cells**

**a.** Immunoblot of total protein extracts from shCTRL and shGPX4 OCI-AML3 cells at the indicated days of DOX treatment. A representative immunoblot of three independent experiments is shown. **b.** Dose–response curves of shCTRL and sh1TGS1 OCI-AML3 cells following treatment with RSL3 for 24 hours at the indicated concentrations. Growth inhibition, calculated as half maximal inhibitory concentration (IC50), is shown. Values are mean ± SD of *n*= 3 independent experiments. Absolute IC50 value, 95% Confidence Interval. **c.** Flow cytometry detection of the reduced and oxidised forms of the BODIPY™ 581/591 C11 probe in shCTRL or sh1TGS1 OCI-AML3 cells, following 5 days of DOX treatment and additional 24 h exposure to DMSO or to 0.3 µM RSL3. **d.** Proliferation assay of shCTRL or sh2TGS1 OCI-AML3 cells measured between day 4 and day 8 after DOX treatment. During the four days of the assay, cells were also treated with 5 µM Ferrostatin 1 (Fer-1) alone or together with DOX. **e.** Reduced and oxidised forms of the BODIPY™ 581/591 C11 probe were detected by flow cytometry in shCTRL or sh2TGS1 OCI-AML3 cells, at the indicated days of DOX treatment. Treatment for 2 hours with 0.5 µM RSL3 alone or in combination with 5 µM Fer-1 were used as positive and negative controls of lipid-peroxidation, respectively. **f.** Cell viability of shCTRL or sh1TGS1 OCI-AML3 cells following 5 days of DOX treatment and additional 24 hours treatment with the indicated concentrations of RSL3 (nM) alone or in combination with 5 µM Fer-1. Cell viability following exposure to DMSO (vehicle control) was used as the 100% viability reference for either shCTRL or sh1TGS1 cells, separately. **g.** shCTRL and sh2TGS1 OCI-AML3 cells following 5 days of DOX treatment were incubated for 24 hours with the indicated concentrations of RSL3 (nM) alone or in combination with 5 µM Fer-1. Cell viability following exposure to DMSO (vehicle control) was used as the 100% viability reference for either shCTRL or sh2TGS1 cells, separately. Values in **c**, **d**, **e**, **f** and **g** are mean ± SD of *n* = 3 independent experiments. *P<0.05, **P<0.01, ***P<0.001 and ****P<0.0001, two-way ANOVA test with Tukey's (**c** and **e**) or Šídák's (**d, f** and **g**) multiple comparisons.

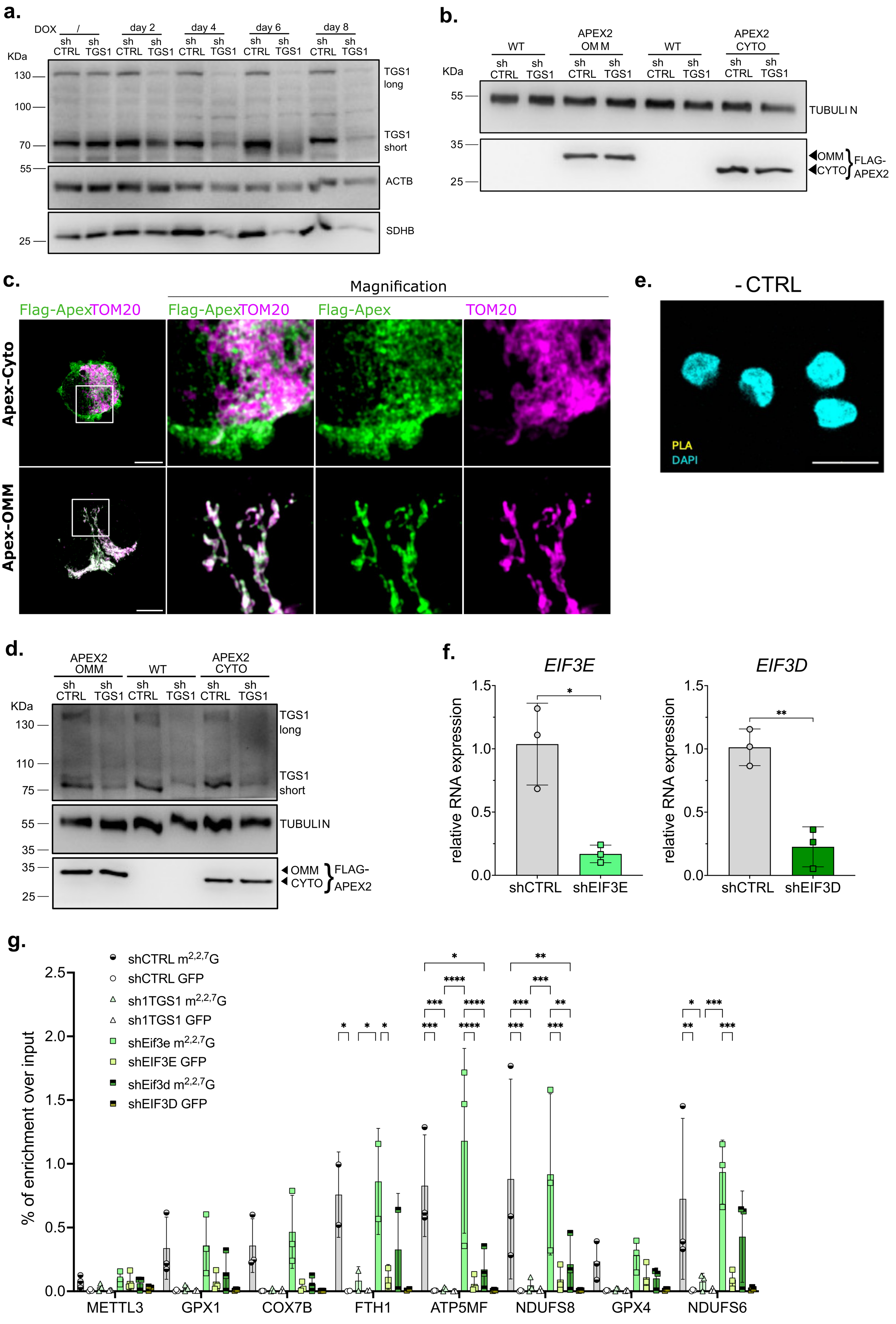

a.
DOX / day 2 day 4 day 6 day 8
sh CTRL sh TGS1
KDa
130
100
70
55
25
TGS1 long
TGS1 short
ACTB
SDHB
b.
WT APEX2 OMM WT APEX2 CYTO
sh CTRL sh TGS1
KDa
55
35
25
TUBULIN
OMM
CYTO
FLAG-APEX2
c.
Magnification
Flag-ApexTOM20
Flag-Apex
TOM20
Apex-Cyto
Apex-OMM
e.
- CTRL
PLA
DAPI
d.
APEX2 OMM WT APEX2 CYTO
sh CTRL sh TGS1
KDa
130
110
75
55
35
25
TGS1 long
TGS1 short
TUBULIN
OMM
CYTO
FLAG-APEX2
f.
EIF3E
EIF3D
relative RNA expression
shCTRL shEIF3E
shCTRL shEIF3D
g.
shCTRL m2,2,7G
shCTRL GFP
sh1TGS1 m2,2,7G
sh1TGS1 GFP
shEif3e m2,2,7G
shEIF3E GFP
shEif3d m2,2,7G
shEIF3D GFP
% of enrichment over input
METTL3 GPX1 COX7B FTH1 ATP5MF NDUFS8 GPX4 NDUFS6

**Supplementary Figure 9. Expression and localisation of APEX2 constructs in HEK-293T cells and knockdown of eiF complex's subunits**

**a.** Representative immunoblot of total protein extracts from shCTRL or sh1TGS1 HEK-293T cells after the indicated days of DOX treatment. ACTB was used as loading control. The images show a representative immunoblot of three independent experiments **b.** Immunoblot of total protein extracts of shCTRL and sh1TGS1 HEK-293T cells transfected with either FLAG-OMM-APEX2 or FLAG-CYTO-APEX2 constructs. A representative immunoblot of three independent experiments is shown. Tubulin was used as loading control. **c.** Representative confocal images of immunofluorescence staining for FLAG-APEX2 (green) and the mitochondrial marker TOM20 (purple) in HEK-293T cells. Scale bar 10 μm (left panels) and indicated magnification (middle, right panels). **c.** Immunoblot of total protein extracts from doxycycline treated (6 days) shCTRL and sh1TGS1 HEK-293T cells transfected with either FLAG-OMM-APEX2 or FLAG-CYTO-APEX2 constructs. A representative immunoblot of three independent experiments is shown. Tubulin was used as loading control. **e.** Representative Proximity Ligation Assay (PLA) images showing the negative control condition (PLA shCTRL), in which one primary antibody was omitted (related to Fig. 9a). No specific PLA signal is detected, confirming the specificity of the assay. **f.** RT-qPCR measure of *EIF3E* and *EIF3D* mRNAs expression in shCTRL and target-shRNAs OCI-AML3 cells following 6 days of DOX treatment; values are mean ± SD of $n$ = 3 independent experiments, *P<0.05 and **P<0.01 unpaired two-tailed t-test. **g.** $m^{2,2,7}$G-RIP-RT-qPCR performed on long RNA purified from shCTRL, sh1TGS1, shEIF3E and shEIF3D OCI-AML3 cells after 6 days of doxycycline treatment. An antibody against GFP was used as a technical negative control. Unmodified *METTL3* mRNA was used as a negative control. The seven TGS1-targets are represented as a global dataset in Figure 9b. Values in **g** are mean ± SD of $n$ = 3 independent experiments, *P<0.05, **P<0.01 and ***P<0.001, two-way ANOVA test with Tukey's multiple comparisons.
